# Single field evolution rule governs the dynamics of representational drift in mouse hippocampal dorsal CA1 region

**DOI:** 10.1101/2024.07.27.605419

**Authors:** Cong Chen, Shuyang Yao, Sihui Cheng, Ang Li, Yusen Yan, Xiang Zhang, Yuanjing Liu, Yumeng Wang, Qichen Cao, Chenglin Miao

## Abstract

How the brain reconciles dynamism with stability to balance learning and reliable memory storage has not yet been fully understood. To address the critical question, we longitudinally recorded place cells in the hippocampal dorsal CA1 region over 7 to 56 days, utilizing multiple goal-oriented navigation paradigms across various environments. We found that over 80% of place cells displayed multiple fields, undergoing complex evolution events including field disappearance, formation, and retention. Place fields from the same neuron showed limited coordination (∼5%), with a preference for synchronized changes. We further uncovered the single field evolution rule: the longer a field remains active, the more likely it is to continue being active; conversely, the longer a field remains inactive, the less likely it is to recover the future fate of a place field depends on its past activity. Mathematical modeling revealed that this rule sufficiently demonstrates the growing stability of the dCA1 spatial representation at the population level.

## Introduction

Dynamism and stability, seemingly incompatible concepts, coexist within the neural network to balance reliable memory storage with the integration of new information. In the hippocampal (HPC) dCA1 region, the exploration of the interrelationship between dynamism and stability has been ongoing since the discovery of place cells^1^, which are pyramidal neurons that encode specific spatial locations known as place fields. Intuitively, the spatial representation of an unchanging environment by the dCA1 should remain stable, providing consistent space coding essential for establishing a cognitive map^2^. However, extensive research has revealed a pronounced dynamic nature of hippocampal spatial representation even in stable environmental conditions, as observed during both novel states^3–7^ and familiar states^8–14^. This phenomenon, termed representational drift (RD), involves substantial changes such as the formation, disappearance, and translocation of place fields and is accompanied by profound refinement of neuronal representation, while still preserving the statistical structure of field properties^12^.

Hippocampal RD can manifest on two distinct timescales: the behavioral timescale and the cross-day timescale. Behavioral timescale RD involves the rapid and robust modulation of HPC representations, governed by behavioral timescale synaptic plasticity (BTSP)^15–17^ and is thought to be experience-driven^7,13^ or learning-related^18^. For instance, HPC spatial representation rapidly changes via BTSP, recruiting more place fields to encode positions associated with recent rewards^18^. As an environment becomes familiar, behavioral timescale RD, indicated by neuronal within-session stability (i.e., representational stability of neurons within a recording session), decreases^3–5,19,20^. However, the overall RD remains prominent, with the correlation of neuronal representation with their initial configuration decreasing to chance levels over weeks^10–13^. In alignment with anatomical evidence from synaptic plasticity^21^ and neurogenesis^22,23^, which function on a week-wise timescale, additional mechanisms might be necessary to drive cross-day RD despite familiarization.

Apart from the hippocampus, cross-day representational drift (RD) has been widely observed in several other brain regions^24–28^, directly invoking intuitive theoretical ideas attributing it to synaptic plasticity. Some have conjectured that cross-day RD may reflect noisy synaptic plasticity during learning^29,30^, such that neural trajectories underlying certain behaviors remain invariant on a low-dimensional manifold^14,31,32^, a framework plausibly supported by the RD with a constant rate observed in the piriform cortex^26^. In the dCA1, however, cumulative studies have reported increasing cross-day stability of spatial representation^6,11,13,33,34^, suggesting factors beyond purely random drift. Despite these qualitative observations, a direct and quantitative measurement of the dynamics of cross-day RD in dCA1 will be essential to shed light on the underlying computational principles, while such estimation has not yet been conducted so far. While synaptic plasticity undoubtedly plays a critical role in representation stability, neuronal impacts should also be considered. Research on memory engrams^35–40^ suggests that stable coding may be confined to particular neuronal ensembles, potentially distinguished by genetic traits^38–41^ or excitability to neuroplastic changes^34^. This raises a critical question: to what extent do dCA1 pyramidal neurons affect the stability of their place fields? If neuronal influence over field stability is substantial, we would expect sibling fields—place fields belonging to the same neuron—to exhibit high coordination in their evolution. Conversely, if place fields maintain independent trajectories, it would indicate underlying principles governing single field stability, rather than being tied to the broader stability of place cells, thereby highlighting a major role for synaptic plasticity.

To address these questions, large-scale studies on place cells with multiple fields (PCmf) are required. Prior studies have predominantly focused on place cells with single fields (PCsf), leaving the long-term dynamics of PCmf largely unexplored. PCmf has been widely reported across various experimental setups such as large-scale linear tracks^12,42,43^, open field^44–46^, and complex mazes^47^, and are proposed to represent a generalized pattern for the dCA1 pyramidal neurons to encode space on a natural scale. With the introduction of additional fields per place cell, latent features of the long-term dynamics of RD, which are obscured by single-field representation, might emerge. By leveraging a spectrum of new analytic methodologies, we dissected the dynamics of cross-day RD with a dataset rich in PCmf and discovered a quantitative rule governing the stabilization of single fields.

We employed longitudinal one-photon calcium imaging to monitor the dCA1 region in freely moving mice engaged in three distinct goal-directed navigation tasks, with or without decision-making processes in complex environments. The primary task was a maze navigation paradigm (MNP) conducted in two complex mazes, Maze A (MA) and Maze B (MB), each incorporating 17 decision points along an 8 to 9-m correct track (Figure 1A). This task was organized into two stages, each lasting approximately 26 days and comprising 13 blocks (Figure 1B). Stage 1 included two 30-min sessions in a familiar open field and one in Maze A. In Stage 2, a session in Maze B was added following the Maze A session (Figure 1C). To assess the directionality of place fields, we implemented a reversed maze paradigm (RMP) where mice navigated back and forth between the entry and exit of Maze A across 7 to 12 sessions spanning 7-15 days (Figures 1H and 1I). Similarly, a hairpin maze paradigm (HMP), without decision points, was employed, mirroring the structure of the RMP (Figures 1H and 1I). This multiplexed design guarantees that our findings are consistent across distinct environments and a spectrum of behavioral conditions, whether involving decision-making or not.

**Fig. 1.**
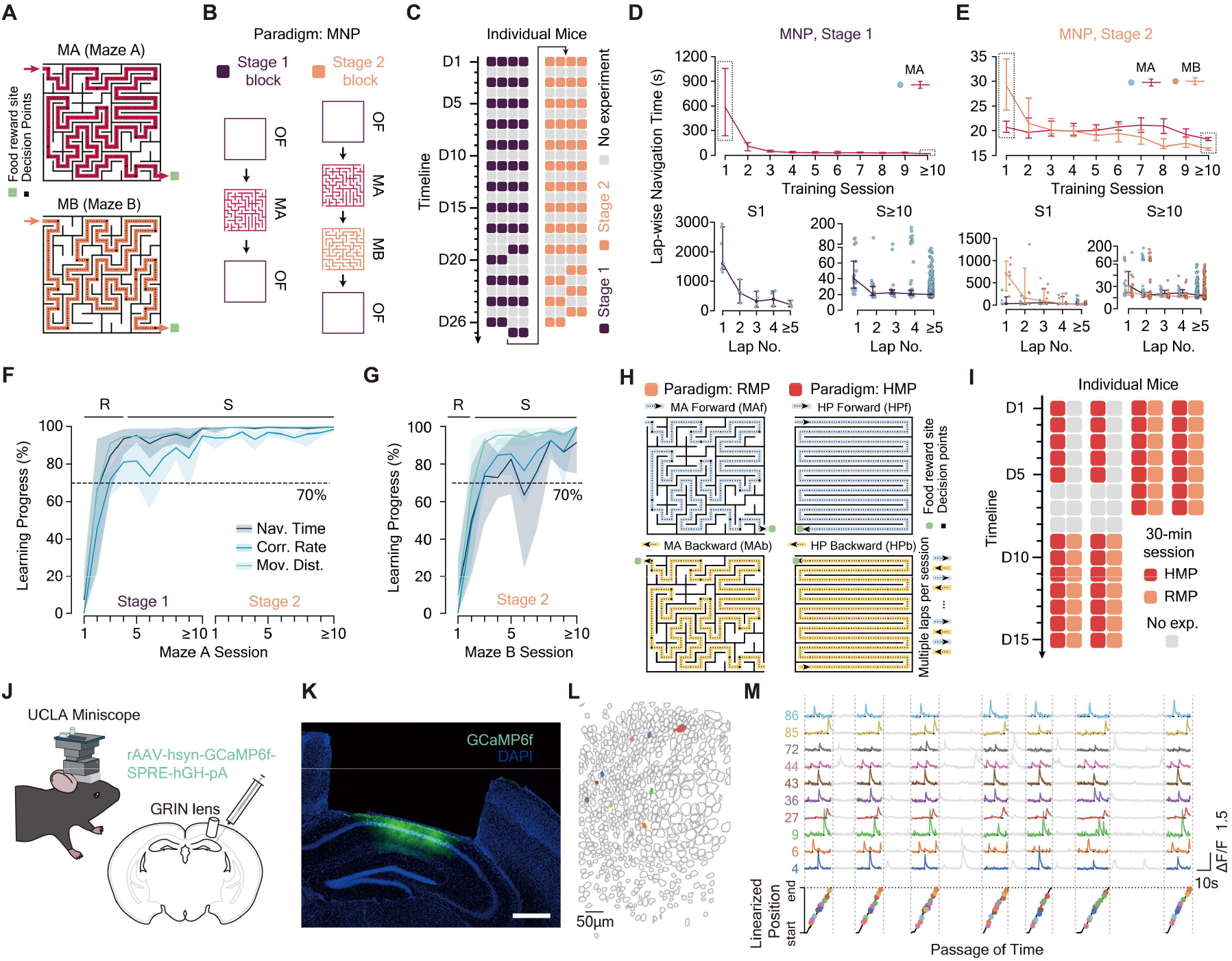
One-photon calcium imaging throughout the engagement of goal-directed navigation in multiple behavioral paradigms. (**A**) Configuration of Maze A (top) and Maze B (bottom). Red and orange lines display the linear representation of the correct paths for Maze A (8.88m) and Maze B (8.08m). MA: Maze A, MB: Maze B. (**B**) Structure of training blocks in Stage 1 (left) and Stage 2 (right). OF: Open Field. (**C**) Timeline of the experiment. Each vertical line represents a mouse, with 13 training blocks conducted over 25∼27 days in each stage. Purple squares: Stage 1 blocks; Orange squares: Stage 2 blocks; Gray squares: no experiments conducted. D: Day. (**D** to **E**) Lap-wise navigation time for Stage 1 (D) and Stage 2 (E). Sessions from ≥10 in each stage are grouped. Bottom panels show navigation times for Session 1 (S1, left) and Sessions ≥10 (S≥10, right) on a per-lap basis. Blue dots: Maze A laps; orange dots: Maze B laps. (**F** to **G**) Learning progress in Maze A (F) and Maze B (G) assessed by lap-wise navigation time (Nav. Time), correct decision rate (Corr. Rate), and lap-wise moving distance (Mov. Dist.). The dashed line at 70% marks the threshold for substantial learning progress, distinguishing the rapid learning phase (R) from the slow improvement phase (S). (**H**) Schematic of the reversed maze paradigm (RMP, left column) and hairpin maze paradigm (HMP, right column), showing moving directions (arrows), decision points (black squares), reward sites (green squares), and the linearized track of Maze A and hairpin maze (dotted line). Light blue represents information related to forward movement, while light yellow represents backward movement. Mice were trained to navigate back and forth between the entry and exit of Maze A for a total of 21-48 laps, with 10-24 laps in each direction. (**I**) The timeline of RMP and HMP training, with 7 HMP sessions for all four mice conducted before the RMP sessions within a day. Each vertical column relates to a mouse. (**J**) Diagram of our surgery and the UCLA Miniscope setup. (**K**) Coronal brain section showing AAV-GCaMP6f expression in the hippocampal dCA1 region. Scale bar: 500 μm. (**L**) An example recording field of view. Gray circles: regions of interest (ROIs); colored, filled circles: example ROIs with activities shown in (M). Scale bar: 50 μm. (**M**) Calcium raw traces from example ROIs demonstrate robust calcium transients and tuning (top) corresponding to correct track on the maze (bottom). Black dots: calcium events; colored dots on linearized position: field centers. Dotted vertical lines indicate the temporal boundary of each lap. Calcium activities within inter-lap intervals were set as gray. Number of mice used: Behavioral training: n = 7 mice in Stage 1, n = 6 mice in Stage 2; Recording: n = 4 mice. Error bars, 95% confidence intervals.

## Results

### The engagement of maze-navigation paradigm

Rodents instinctively excel in navigating within complex environments akin to the subterranean tunnels in where they naturally dwell. In their initial encounter with Maze A, mice averaged 1979.8 ± 645.5 seconds to complete their first lap (Figure 1D) to locate the exit, traversing a path of 203.1 ± 56.7 m (mean ± std.; Figures S1A and S1B), with lighting conditions having no apparent effect on the performance (Figure S1C). During Stage 1 Session 1 (S1), mice exhibited a modest correct-decision rate of 40.3 ± 8.9% at decision points in Maze A (mean ± std; Figure S1D), with an average speed of 11.7 ± 2.8 cm/s (Figure S1E) and a similar visit rate to both correct and incorrect tracks (Figure S1F). A rapid learning phase (R phase) was observed in the initial three sessions, marked by significant improvements exceeding 70% of their behavioral progress (MA, Stage 1, S1 to S3: lap-wise navigation time reduction, lap-wise moving distance, and correct decision rate improvements of 81.5 ± 19.4%, 90.7 ± 8.4%, and 71.1 ± 8.7%, respectively; Figures 1F, S1A, S1B, and S1D). This R phase transitioned into a slow improvement phase (S phase) with continuous refinement of behavioral metrics (Figures 1D-F, S1A, S1B, S1D, and S1G). Upon familiarization with Maze A, mice underwent Stage 2 training in Maze B, exhibiting similar learning patterns (Figures 1E, 1G, S1A, S1B, S1D, and S1G). Key performance indicators, including lap-wise navigation time, moving distance, correct decision rate, and mean speed, all demonstrated significant enhancements (Figures 1E, 1G, S1A, S1B, S1D, and S1G). Ultimately, mice not only mastered the maze exit route but also optimized their path (MA: 6.52 m; MB: 5.88 m; medians), achieving high accuracy in correct decisions (Maze A: 97.2 ± 2.5%; Maze B: 97.5 ± 1.5%, Figure S1D) and mean speed (Maze A: 34.7± 6.5 cm/s; Maze B: 34.7 ± 6.8 cm/s, Figure S1E), showcasing robust and precise spatial learning.

### Enhanced within-session stability and precision of hippocampal dCA1 spatial representation throughout maze learning

To investigate the long-term dynamics of hippocampal spatial representation, we employed the UCLA Miniscope (Figure 1J) to record neural activity in the dCA1 region of freely moving mice. We injected rAAV-hsyn-GCaMP6f-SPRE-hGH-pA virus to express the calcium indicator GCaMP6f (n = 4 mice, Figures 1J and 1K), yielding 447.4 ± 175.4 regions of interest (ROIs) per session (n = 363 sessions; Figures 1L and S2A). These sessions demonstrated robust calcium activities with spatial tuning (Figure 1M). Over 90% of active ROIs in both mazes (Stage 1 MA: 94.7 ± 5.7%; Stage 2 MA 95.3 ± 5.0%; MB 93.0 ± 7.0%; Figures S3A and S3B) qualified as place cells, a significantly higher proportion than observed in the open field (Stage 1: 60.7 ± 13.0%; Stage 2: 66.0 ± 10.2%; Figures S3A and S3B). Moreover, Maze A and B collectively shared 92.5 ± 8.0% of place cells among all tracked active neurons (median 96.3%), (Figure S3C), underscoring the high fraction of place cells across different complex mazes.

The within-session stability of these place cells, as gauged by the half-half Pearson correlation of their event rate maps, showed marked improvement in both mazes over the course of MNP training (Figures S3D-F). This contrasts with the stability in the open field, which remained relatively unchanged (Figures S3D-F). Alongside this increased stability, there was a notable rise in the spatial information provided by place cells (Figures S3G-I), unlike in the open field, where it stayed consistent (Figures S3G-I). Additionally, a slight uptick in mean event rate was recorded (Figures S3J-L). Decoding errors exhibited a significant reduction in both Maze A and B (Figures S3M-O), in stark contrast to the open field, where errors remain unchanged (Figures S3O-Q). These results underscore a notable improvement in the precision and within-session stability of hippocampal spatial coding in complex mazes as mice became familiarized with these environments.

### Canonical statistical structures of multi-field spatial maps in complex mazes and hairpin maze

During MNP training in Maze A, only about 3% of place cells had a single field in both Stage 1 (1.7 ± 1.5%) and Stage 2 (3.2 ± 2.3%) a proportion mirrored in Maze B (3.0 ± 2.5%; Figures S4 and S5A-C). Most place cells in these mazes exhibited multiple fields, averaging 8.0 ± 1.9 fields per cell in Maze A and 7.3 ± 1.2 in Maze B (Figures 2A, 2B, S4 and S5D-F), regardless of the criteria for place fields identification (Figures S5G-K). The RMP revealed that place cells in complex mazes are highly directional, with over 85% of active neurons recruiting as place cells to encode positions of Maze A during both directional movements (MA forward: 90.6 ± 7.2%; MA backward: 85.7 ± 8.5%) and 81.8 ± 10.9% encoding both directions (Figures 5L and 5M). Only 19.6 ± 5.6% of place fields overlapped between directions, a mild but significant difference compared to shuffled results (14.8 ± 4.9%, Paired t-test, P = 1.3 × 10^−13^), suggesting that the spatial maps encoding each direction were nearly distinct. In the hairpin maze, a lower fraction of place cells encoded each direction (HP forward: 76.7 ± 9.9%; HP backward: 73.8 ± 14.8%), with a lower fraction of bi-directional place cells (62.7 ± 15.4%; Figures S5F and S5G) and place fields (17.0 ± 5.2%; significantly higher than shuffle: 12.8 ± 4.4%, Paired t-test, P = 3.9 × 10^−11^). Despite this diversity, all spatial maps displayed extensive multi-field coding, with exceeding 90% place cells exhibiting multiple fields in all paradigms (MA forward: 93.6 ± 3.5%; MA backward: 94.0 ± 3.1%; HP forward: 95.3 ± 2.7%; HP backward: 94.1 ± 3.6%) and an average field number well above one (MA forward: 6.4 ± 1.7; MA backward: 7.3 ± 1.8; Figures 2C, S5N and S5O; HP forward: 8.6 ± 2.6, HP backward: 8.0 ± 2.2; Figures 2D, S5S and S5T). For clarity, place cells’ activities detected during training in Mazes A and B in the MNP, as well as during forward and backward movements in the RMP and HMP, are collectively referred to as six spatial maps: MA, MB, MAf, MAb, HPf, and HPb.

**Fig. 2.**
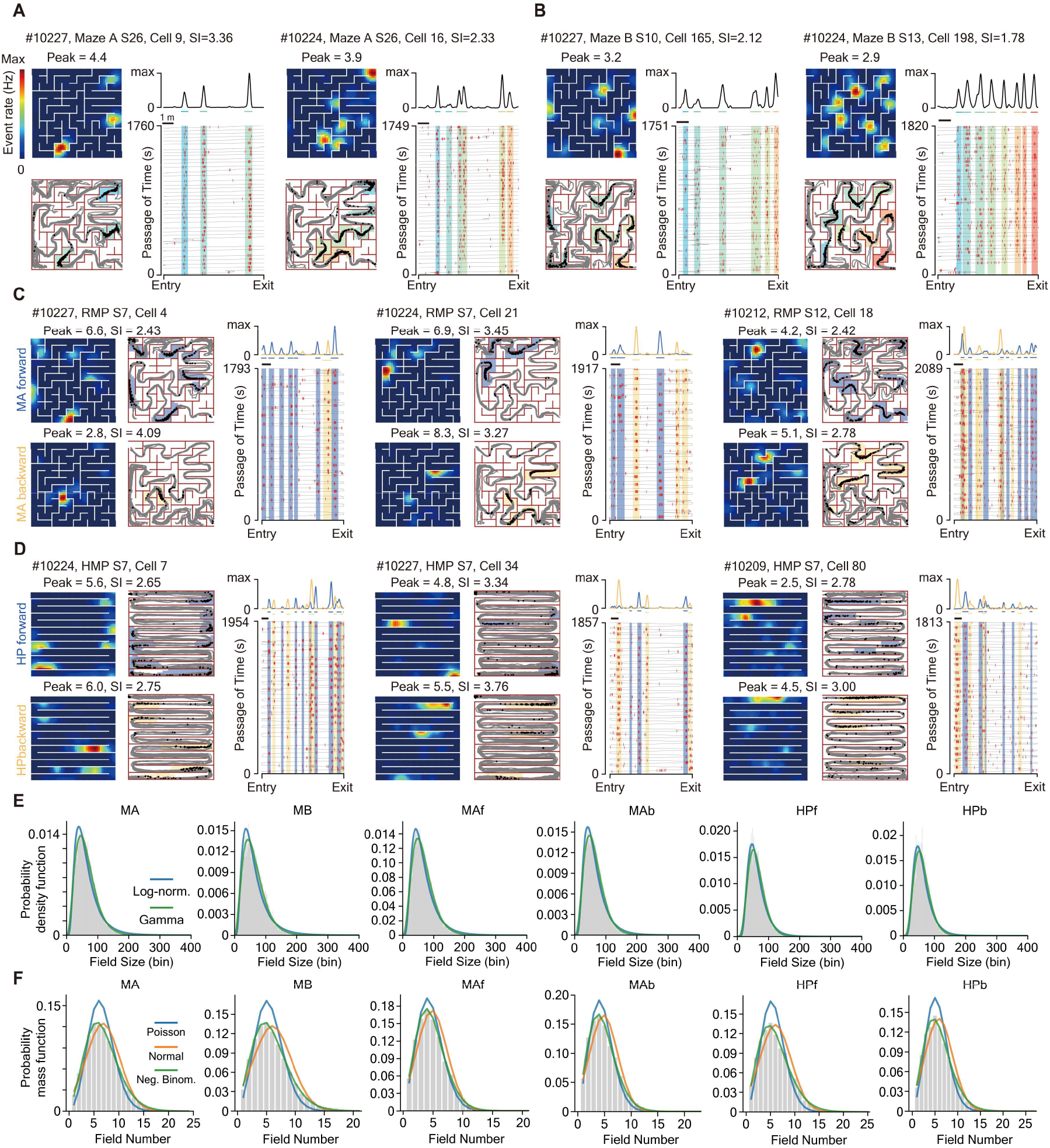
Most place cells in six spatial maps exhibit multiple place fields with canonic statistic structures. (**A** to **D**) Example PCmf detected in the six spatial maps (**A**: Maze A during MNP; **B**: Maze B during MNP; **C**: MAf and MAb during RMP; **D**: HPf and HPb during HMP). Representative place cells in MNP (A and B) are visualized in four subfigures, entitled with the ID of mice, recorded session, cell index, and spatial information (SI, unit: bits per spike). top left: event rate maps, peak: peak event rate (unit: Hz); bottom left: mouse trajectory (gray lines) with spatial distribution of calcium events (black dots); top right: linearized rate map along correct track; bottom right: temporal distribution of calcium events, with red bars indicating calcium events and gray lines representing linearized mouse trajectory in correct path. Colored background shadows in bottom figures and colored bars below the linearized rate map mark the ranges of individual place fields, with the same color using for the same field. Black bars under the linearized rate map mark the length of 1 m along the linearized correct track of Maze A, Maze B, and the track of hairpin maze. Calcium activities on incorrect tracks and backward movements are excluded. For PCmfs detected in RMP and HMP, their event rate maps and trajectory maps were visualized separately, while their activity along the linearized track were shown together. Colored shadows mark the range of individual place fields for both forward (bluish violet) and backward (light yellow) spatial maps, and black bars indicate 1 m length of linearized track of either. (**E**) Examples from mouse #10209 demonstrate a skewed distribution of place field sizes (unit: spatial bin, each equal to 4 cm^2^) in all the six spatial maps, which can be well-captured by gamma distributions (green line) (Lilliefors-corrected KS test, P > 0.26 for all spatial maps; fig. S6). (**F**) Examples from mouse #10227 demonstrate a Poisson-shape distribution of field number per cell in all the six spatial maps, with negative binomial distribution (neg. binom., green line) outperforms the Poisson (Blue line) and normal distribution (orange line) (Lilliefors-corrected KS test, P > 0.64 for all spatial maps; fig. S7).

To ensure that our multi-field spatial maps were not artifacts produced by flawed field identification criteria, we corroborated that these maps conformed to canonical statistical structures for field sizes^43^ and field numbers^12,42^. Specifically, field sizes across six spatial maps adhered to skewed distributions^43^ (Figures 2E and S6). Moreover, the distribution of field counts per neuron throughout all goal-directed navigation paradigms was precisely captured by a negative binomial distribution (Figures 2F and S7). This observation dovetails with the Gamma-Poisson hypothesis (see Methods), suggesting each dCA1 pyramidal neuron harbors a distinct intrinsic propensity, denoted as *λ* (average number of fields per meter), to generate place fields across various contexts and environments. While prior analyses highlighted a significant variance in this propensity across neurons— evidenced by a negative binomial distribution that typically dwindles monotonically^12,42^— our findings depict a Poisson-like distribution for all examined spatial maps (shape parameter r > 1 in all 26 spatial maps from 6 mice, with a minimum value of 3.09; Figure S7). Such an outcome hints at a potentially constrained variance in neuronal propensity for place field formation.

Subsequent examinations verified that the field numbers of identical neurons exhibit significant correlations across distinct mazes (Pearson correlation, MA vs. MB, 0.39 ± 0.11; Figures S8A and S8B) and different movement directions (MAf vs. MAb, 0.53 ± 0.08; HPf vs. HPb, 0.51 ± 0.14; Figures S8C-F). These findings align with previous research, underscoring the existence of a neuronal propensity to form fields^12,43^. Our analysis to determine the true distribution of this neural propensity (see Methods) revealed a consistent adherence to a gamma distribution across all four mice (Kolmogorov-Smirnov test, P > 0.15; Figures S8G and S8H), with coefficient of variation (CV) more than 3 times lower than previously reported values. This denotes a more constrained heterogeneity in the propensity across the neural population. Hence, we ascertained that the multi-field spatial maps we documented not only showcase canonical statistical structures but also align with the principles of the Gamma-Poisson model. This confirmation solidifies the basis for our subsequent analyses on the dynamics of multi-field representations.

### The properties of sibling fields are highly independent within sessions

To assess the long-term autonomy of sibling place fields, a first estimation of the independence across their properties such as size, firing strength, and within-session stability, provides essential insight. The Gamma-Poisson model posits that individual place fields are independent events, randomly assigned to neurons to form multi-field spatial maps through a homogeneous spatial Poisson process^42^. This model implies that sibling fields should exhibit properties largely independent from each other, akin to non-sibling fields. However, this assumption has not yet been rigorously validated. To understand the extent to which the properties or functional processes of place fields are influenced by their parent neuron, we employed three distinct analytical approaches: probabilistic principles, information theory (Figures S9A-C), and shuffle tests (Figures S9F and S9G). We utilized three specific metrics— *χ*^2^, mutual information, and standard deviation—to compare between sibling and non-sibling field groups. Notable differences of these metrics between sibling and non-sibling field group would suggest potential interactions among sibling fields’ properties. Our analyses confirmed a high degree of independence in the properties of sibling fields. Both *χ*^2^ and mutual information metrics (Figures S9D and S9E) indicated no significant differences between sibling (Sib) and non-sibling (Non) groups in most comparisons (two-sided two-sample t-test, P > 0.3 for field center rate and sizes in both mazes and both metrics), except for a nuanced relationship in their within-session stability (*χ*^2^, P < 1 × 10^−6^ for both mazes; MI, P > 0.13 for both mazes). Likewise, analyses of standard deviation did not reveal any significant differences, in terms of either mean or distribution, between sibling and non-sibling groups in most sessions (Figures S9F-N). These findings corroborate the high independence of field properties among sibling fields, while the slight dependency observed in within-session stability hints at some extent of influence of the parent neuron on the stability of its fields.

### Sibling fields evolve with significant but limited coordination

The prevalence of PCmf identified in our dataset provides a unique opportunity to directly quantify the influence of parent neurons on the long-term stability of individual place fields. We meticulously tracked neurons across six spatial maps for extended periods (Figure S2). Notably, a substantial number of place fields demonstrated remarkable durability (Figures 3A and S10), with 255 fields continuously active for at least 40 days (18 sessions; from n = 2 mice) and 87 fields retained for a minimum of 50 days (23 sessions; from n = 2 mice; Figure 3B) in Maze A. These long-lasting fields often coexisted with sibling fields that exhibited comparatively shorter lifespans, highlighting a diversity in field longevity within the same neuron (Figures 3A and S10). Despite a great number of highly stable place fields, the evolution of multi-field place cells involves complex and dynamic processes, with fields emerging and disappearing over time (Figures 3A and S10). To dissect these knotty dynamics, we decomposed the evolution of each place field into three elementary categories of evolution events across consecutive sessions: the disappearance, formation, and retention of field (duration = 2 sessions, Figure 3A). To confirm if we extracted these evolutionary events correctly, we checked the changes of event rate within the place fields across successive sessions and observed significant reductions, increases, and mild changes respectively (Figures 3C-E). Notably, 15∼45% weakened or even disappeared place fields have a significantly greater than chance probability of recovery within the next five sessions (cumulative probability subtracted with chance level: MA: 38.8 ± 3.4%; MB: 36.4 ± 1.9%; MAf: 29.4 ± 9.2%; MAb: 28.3 ± 9.4%; HPf: 25.7 ± 2.6%; HPb: 25.2 ± 2.0%; Figures 3F and 3G). Given the variety of evolutionary events occurring within the same place cell, the evolution of sibling fields is qualitatively less likely to be highly coordinated over extended periods.

**Fig. 3.**
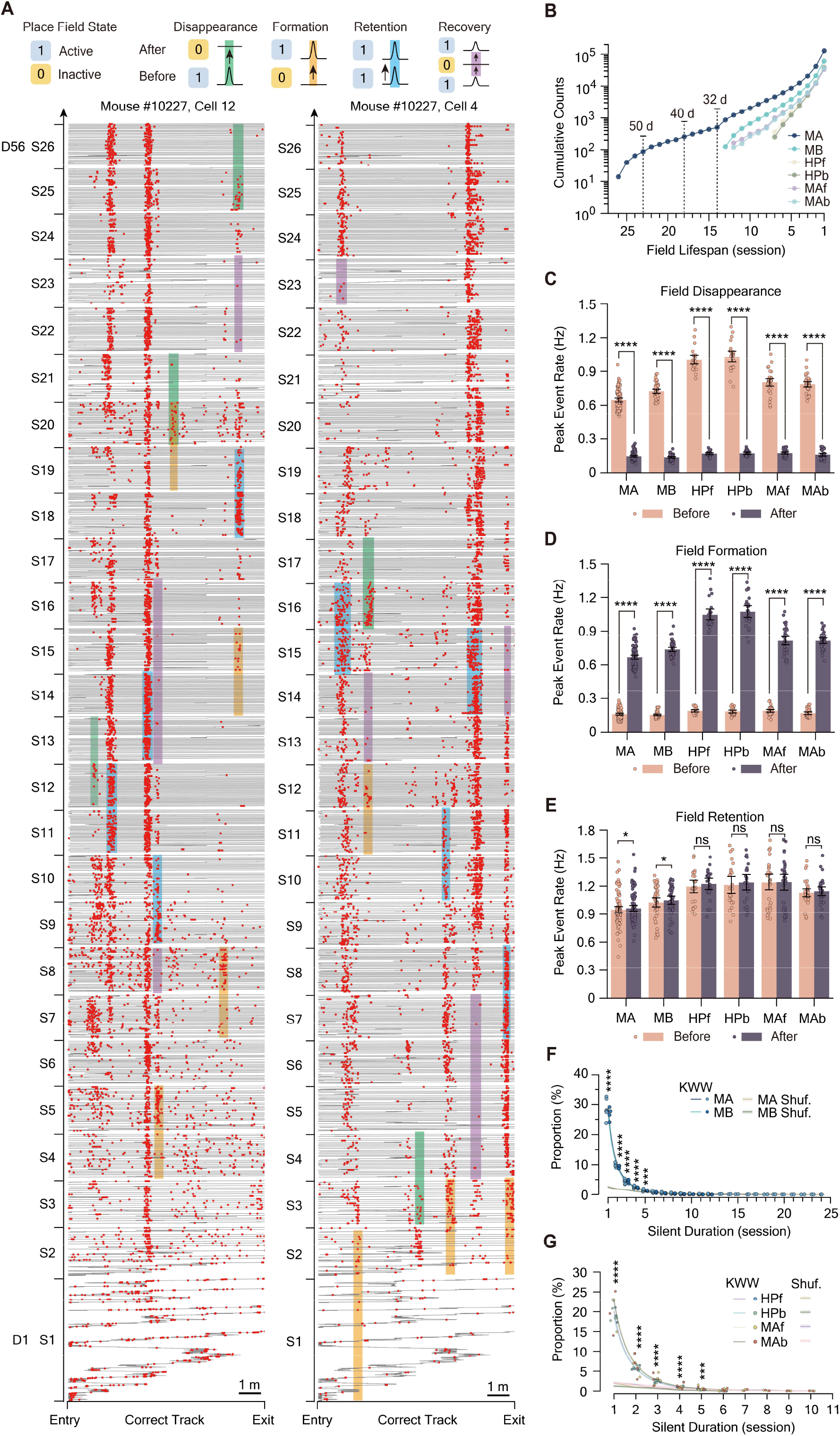
Diverse evolution events in the long-term dynamics of hippocampal multi-field spatial maps. (**A**) Throughout MNP training lasting 26 to 56 days, our observations revealed complex changes in the spatial representation of PCmf. Two exemplary place cells demonstrate a spectrum of intricate evolution of place fields in their spatial maps. These changes were classified into three fundamental categories of evolution events: disappearance, formation, and retention of place fields, marked with green, yellow, and blue shadows, respectively. Beyond these basic categories, we documented the recovery of previously weakened place fields, highlighted by purple shadows to indicate their periods of dormancy. Red bars indicate individual calcium events, and the background gray lines depict the mice’s trajectories on the correct track of Maze A throughout the maze navigation paradigm training. Sessions are labeled as S1 to S26 (corresponding to Stage 2 S13). (**B**) Lifespans of all tracked place fields across six spatial maps are displayed. The graph shows the cumulative count of place fields that exceed specific session durations, summed across data from four mice. Lifespans of 14, 18, and 23 sessions correspond to approximately 32, 40, and 50 days, respectively. (**C** to **E**) Changes of peak event rate (unit: Hz) within place fields during field disappearance (C), formation (D), and retention (E). A two-sided Paired t-test was used to compare the peak rates within place fields before and after evolution events, with the following degrees of freedom (df): MA, 96; MB, 47; HPf and HPb, 23; MAf and MAb, 33. (**C**) Peak rates within place fields drastically decreased (Two-sided Paired t- test, P < 4 × 10^−20^ for all comparisons) during disappearance events. (**D**) Peak rates within place fields significantly increased (P < 2 × 10^−19^ for all comparisons) during formation events. (**E**) Peak rates within place fields showed only mildly significant changes (MA, P = 0.023; MB, P = 0.015) or were non-significant (P > 0.26 for HPf, HPb, MAf, and MAb) during retention events. Peak rates within fields undergoing specific evolution events were averaged across each pair of successive sessions and presented as individual dots, allowing for the analysis of changes in event rates before and after specific evolutionary occurrences. (**F** and **G**) Fraction of recovered place fields among all once weakened fields with respect to their continuous silent duration (recovery interval) observed across six spatial maps. Chance levels computed by re-locate shuffle test (see Methods) are indicated by lines with shadows, and significance levels are denoted by colored stars. Significance levels: ns, P ≥ 0.05; *, P < 0.05; **, P < 0.01; ***, P < 0.001; ****, P < 0.0001. Calcium activities on incorrect tracks and backward movements are excluded. Error bars, 95% confidence intervals.

Nevertheless, we proposed two alternative hypotheses to account for the observed dynamics: the independent drift hypothesis, which assumes high autonomy among sibling place fields, and the coordinated drift hypothesis, which suggests some level of coordination influenced by the parent neuron. To quantitatively assess potential coordination over short (2 to 3 days) or long (∼8 days) durations, we expanded our definition of evolution events to include contiguous 3 to 5 sessions (Figure 4A). This approach allowed us to analyze these extended events alongside the basic three categories of evolution that are defined over a duration of 2 sessions. We applied statistical methods previously used to assess field properties’ independence to now explore the coordination in the evolution of sibling place fields. Joint probability matrices computed from sibling fields, non-sibling fields, and simply multiplication of marginal probability (expected joint probability) (see Methods; Figure 4B) were used to analyze whether evolutionary events such as field retention, disappearance, or formation cooccurred more or less frequently in sibling fields than expected under independence or compared to non-sibling fields (Figures 4C and 4D). Both *χ*^2^ and mutual information metrics revealed significantly higher values for sibling groups compared to non-sibling groups across all six spatial maps (Welch’s t test, *χ*^2^, MI: P-values < 0.0005; Figures 4E, 4F, S11A and S11B), demonstrating notable coordination. This coordination was evident irrespective of the duration used to define evolutionary events (*χ*^2^, MI: P-values < 0.0002; Figures 4E, 4F, S11A and S11B). Although coordination is significantly present, its degree is limited, with only about 5% coordination observed (Figure S11C and S11D). This level of coordination was consistently low across the six spatial maps, regardless of the behavioral paradigms and duration of evolution events (Figure S11C and S11D), and it persisted over time (Figure S12), indicating a subtle but enduring form of interrelation in field evolution.

**Fig. 4.**
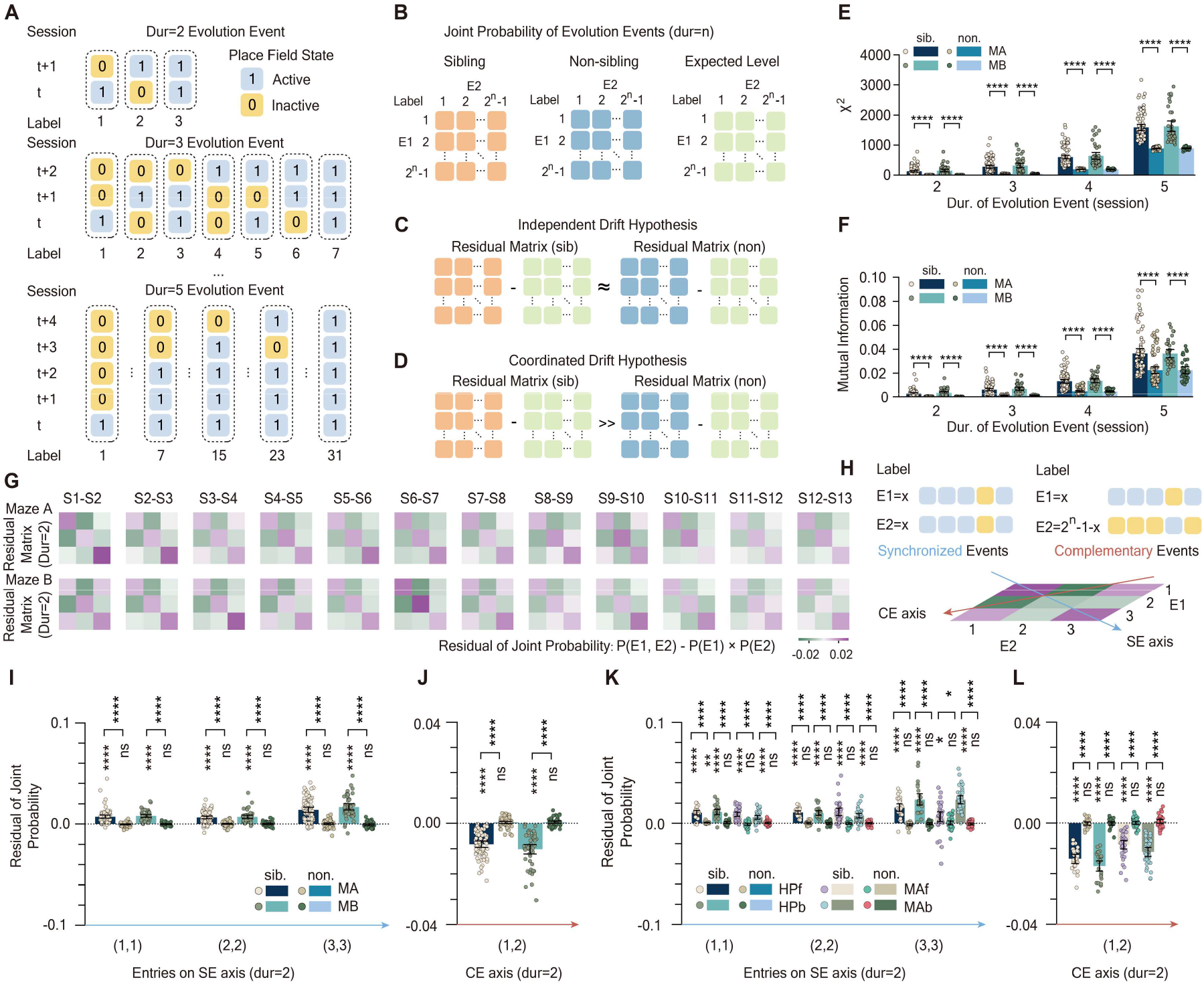
Sibling fields evolve with limited coordination, preferring synchronized changes. (**A**) Evolution event schematics for durations of 2 to 5 sessions. St denotes session t. Each evolution event has a unique label (see Methods). Blue squares: active place fields; yellow squares: inactive place fields. With a duration of 2, evolution events labeled as 1, 2, 3 represent disappearance, formation, and retention of fields, respectively. (**B**) Co- occurrence probability matrices for evolution events with a duration of *n*, computed for sibling fields (orange matrix, top), non-sibling fields (blue matrix, middle), and expected level (green matrix, lower). E1, E2 denote random variables (see Methods). (**C** and **D**) Schematic illustration of the independent drift hypothesis (C) and coordinated drift hypothesis (D). Residual matrices are computed by subtracting either the orange matrix or the blue matrix with the green matrix displayed in (B). ≈: no significant distinction; >>: significantly greater than. (**E** and **F**) Analysis of residual matrices using *χ*^2^ statistics (E) and mutual information (F). Two-sided two-sample t-test was applied. (**G**) Examples of residual matrices with a duration of 2 over 12 sessions for both Maze A (top line) and Maze B (bottom line). S1: Session 1. Colored bars represent residual probabilities. (**H**) Illustration of synchronized event pairs (SE, top left) and complementary event pairs (CE, top right). A SE pair consists of two events with identical states on each day, whereas a CE pair consists of two events with opposite states on each day. The bottom example residual matrix shows that the residual probability of SE pairs is distributed along the main diagonal (blue arrow) of the residual matrix, while CE pairs are distributed along the counter diagonal of 2^*n*^ − 2 degrees (dark red arrow). (**I** to **L**) Statistical analysis identifying which evolution events contribute to coordination within the spatial maps of MA, MB (I and J), MAf, MAb, HPf, and HPb (K and L), along the SE axis (I and K) and CE axis (J and L). Abbreviations: MA, Maze A; MB, Maze B; HP, hairpin maze; f, forward; b, backward. Non.: non-sibling group. Sib.: sibling group. Statistical tests under the brackets: two-sided one-sample t-test with the alternative hypothesis that values are greater than 0 (I and K) or less than 0 (J and L). Above the brackets: two-sided two-sample t-test comparing sibling and non-sibling groups. Significance levels: ns, P ≥ 0.05; *, P < 0.05; **, P < 0.01; ***, P < 0.001; ****, P < 0.0001. Only sibling fields with inter-field interval greater than 0.5 m were considered. Error bars, 95% confidence intervals.

**Fig. 5.**
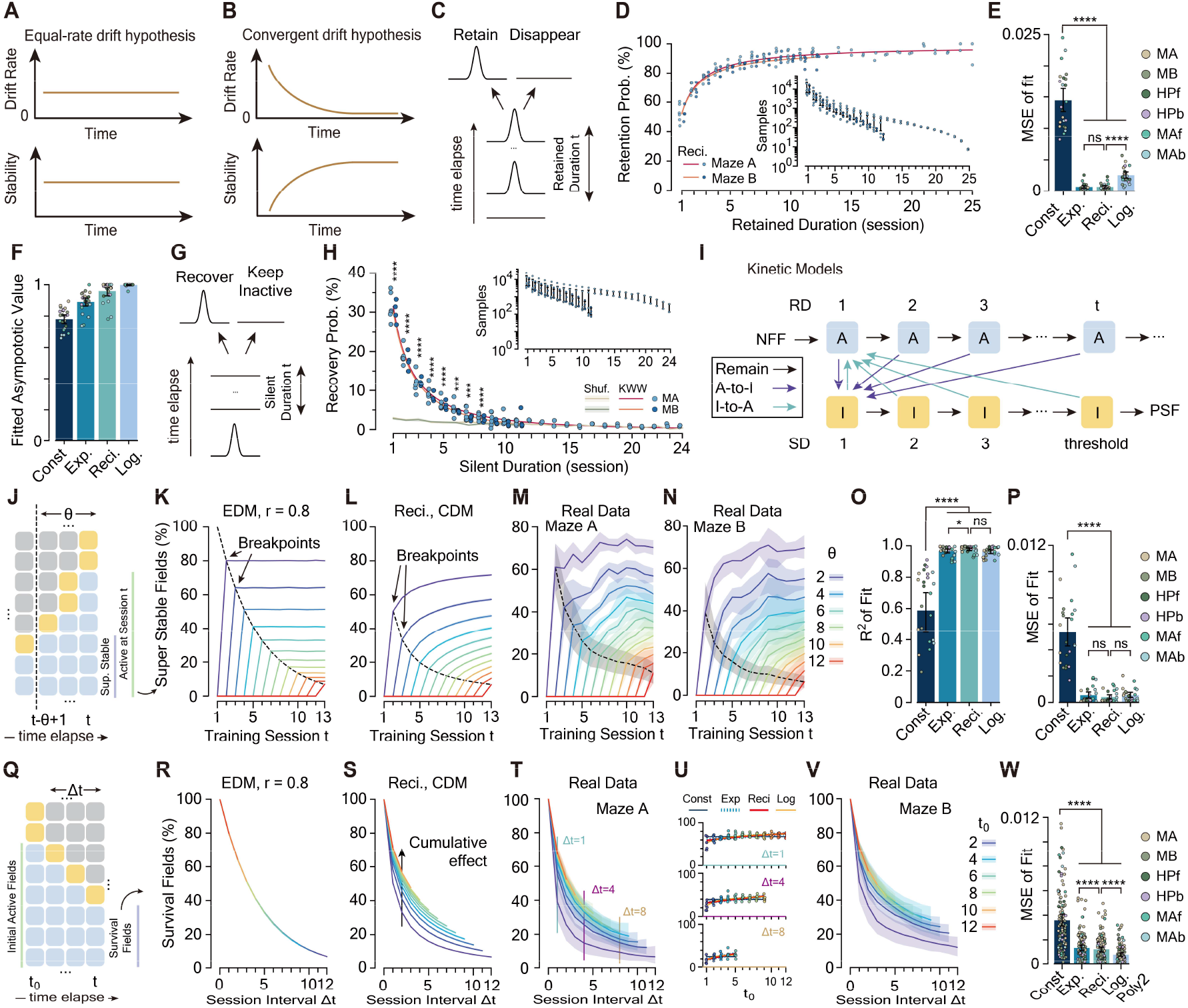
Single field evolution rules are sufficient in description of dCA1 representational dynamics. (**A** and **B**) Schematic illustration of the equal-rate drift hypothesis (A) and the convergent drift hypothesis (B). Solid line: trend of drift rate (top), defined at the single field level; and stability (bottom), defined at the ensemble level. (**C**) Schematic of conditional retention probability. (**D**) Measured field retention probability (dots with black outlines) with respect to retained duration (unit: session), fitted with the ‘reci.’ function (red and orange solid lines, see Methods). Inset: number of samples used for each retained duration. Dots with fewer than 5 samples were excluded. (**E**) field retention probability was fitted with either constant function (‘const.’) or convergent functions (‘exp.’, ‘reci.’, ‘log.’) on a per mouse basis and mean square error (MSE) of fit was adopted as a metric to assess the goodness of fit (GOF). Two-sided paired t-test with Bonferroni correction was used to compare the GOF between constant and convergent functions (P < 0.0001 for all comparisons, df = 23). ‘Reci.’ Provides the best fit with the least average MSE (0.00067 ± 0.00052). (**F**) The fitted asymptotic values to which convergent functions approach with respect to retained duration. (**G**) Schematic of conditional recovery probability. (**H**) Measured field recovery probability (dots with black outlines) with respect to silent duration (unit: session), fitted with the Kohlrausch-Williams-Watts (KWW) function (red and orange solid lines) on a per mouse basis. Wild-green and light-yellow lines: chance levels from field re-location shuffle test (Methods) for Maze A and B, respectively. (**I**) Schematic of kinetic models. Blue squares: active fields; yellow squares: inactive fields. Black arrows: state retention; purple arrows: A-to-I transitions; cyan arrow: I-to-A transition. CDMs and EDMs differ only in A- to-I transition probabilities. RD: retained duration; SD: silent duration; NFF: newly formed fields; PSF: permanent silent fields. (**J** to **N**) Dynamics of the percentage of super stable fields in EDMs (K), CDMs (L), and real data in Maze A (M) and B (N). (J) Illustration of super stable field fraction calculation. Squares indicate active fields (blue), inactive fields (yellow), or either (gray), the same in (*q*). Dotted vertical line sets threshold *θ*, with super stable fields (purple line) and all active fields (green line) marked in session t. Dynamics with varied *θ* in (J to N) were shown in rainbow colors. Black dotted line and arrows indicate predicted breakpoints distribution (Methods). Simulation: 50 runs. Shadows indicate 95% confidence intervals. Black dotted line with shadows shows actual breakpoints distribution. **(O** and **P**) The breakpoints observed in the real data were fitted with either EDM- or CDM- predicted functions, and the GOF, in terms of the coefficient (*r*^2^) (O) and MSE (P), demonstrates that CDMs provide a better description of the trend of percentage of super stable fields (Two-sided Paired t-test with Bonferroni correction, P < 0.0001 for all comparisons, df = 23). (**Q** to **V**) Dynamics of survival field fraction in EDMs (R), CDMs (S), and real data in Maze A (T and U) and B (V). (Q) Illustration of survival field fraction calculation. Dotted vertical line sets session interval Δt, and *t*_0_ is the initial session. Green line marks initial active fields, purple line marks survival fields. Arrow in (S) highlights cumulative effect predicted by CDMs with fixed Δ*t* and increasing *t*_0_, absent in EDMs. Rainbow colors in (R to V) represent lines with different *t*_0_ values, with shadows indicating 95% confidence intervals. (U) visualizes the trends of the cumulative effect observed in Maze A (T) for Δ*t* = 1 (light blue, top), 4 (purple, middle) and 8 (beige, bottom). The cumulative effect was fitted with both EDM- and CDM-predicted functions (n = 4 mice), whose GOF was assessed by MSE in (**W**) (Two-sided Paired t-test with Bonferroni corrections, P < 0.0001 for all CDM-EDM comparisons). Significance levels: ns, P ≥ 0.05; *, P < 0.05; **, P < 0.01; ***, P < 0.001; ****, P < 0.0001. Error bars, 95% confidence intervals.

To determine which evolution events contribute to the observed limited coordination among sibling fields, we analyzed residual matrices computed by subtracting the joint probabilities of sibling field groups to expected values (see Methods; Figures 4G and S13-15). Our analysis revealed that entries along the main diagonal (SE axis), which represent the probability of synchronized evolution events (Figure 4H), consistently exhibited mild yet significant positive residuals (Figures 4I, 4K, S14C, S14F, S14I, S15B, S15E and S15H), a level significantly higher than non-sibling group and the expected level. These findings suggest that synchronized events occur among sibling fields more frequently than expected by chance, indicating that such preference significantly contributed the observed coordination. In contrast, the entries along the secondary counter diagonal (CE axis), which represent complementary evolution events (Figure 4H), consistently showed negative values for durations of 2 to 4 sessions (Figures 4J, S14D and S14G), suggesting a persistent disfavor towards alternating changes. However, for longer durations (5 sessions), most of these entries no longer exhibited significant negative values (Figure S14J), remaining the possibility that the observed negative values along the CE axis may be a consequence of the positive values on the SE axis. This pattern of preference for synchronized evolution and disfavor of alternating evolution among sibling fields was consistent across the other four spatial maps (MAf, MAb, HPf, and HPb; Figure 4K, 4L, S15C, S15F and S15I). Although these preferences have been identified, they might only partially contribute to the observed coordination. Given the already limited overall level of coordination, these preferences are consequently unlikely to significantly impact the long-term evolution of sibling fields. These findings endorse the coordinated drift hypothesis, yet they also represent a compromise with the independent drift hypothesis by acknowledging that place fields maintain a high, albeit not entire, autonomy in their long-term fates.

### Probabilistic description of rules governing the evolution of single place field

With high degree of place field’s autonomy, it becomes possible that the mystery of the dynamism-stability duality of the dCA1 spatial representation is hidden behind some potential rules governing the evolution of single place fields. To explore this possibility, we began by addressing a transformative question: considering the diverse evolution events observed, how do place fields switch between active (A) and inactive (I) states? We initially concentrated on the pivotal A-to-I transition, which fundamentally determines the drift of individual fields and influences the overall stability of the ensemble representation. We proposed two hypotheses: the equal-rate drift hypothesis, which suggests that the probability of an A-to-I switch *P*_*A*→*I*_ remains constant regardless of the field’s retained duration (R_t_), maintaining a consistent level of system stability (Figure 5A); and the convergent drift hypothesis, which posits that *P*_*A*→*I*_ monotonically decreases and converges to a low level as a place field’s retained duration extends, leading to increasing stability within the system (Figure 5B). We assessed the conditional retention probability (1 − *P*_*A*→*I*_), representing the likelihood of a place field remaining active in the subsequent session given its current retained duration (Figure 5C). We observed a significant increase in this probability as the retained duration lengthens (from R_t_ = 1 to 12, MA: 50.9 ± 4.9% to 94.2 ± 1.8%, Paired t test P = 3.5 × 10^−5^, df = 4; MB: 52.2 ± 3.6% to 85.0 ± 4.0%, P = 0.002, df = 3; Figure 5D). A similar trend was evident across the other four spatial maps, contradicting the equal-rate drift hypothesis (from R_t_ = 1 to 6, MAf: 55.1 ± 4.9% to 90.8 ± 2.7%, P = 0.002; MAb: 52.4 ± 4.3% to 89.4 ± 4.2%, P = 0.003; HPf: 46.8 ± 5.3% to 82.7 ± 8.7%, P = 0.012; HPb: 45.0 ± 5.0% to 77.3 ± 3.3%, P = 0.003; df = 3; Figure S16A).

Moreover, fitting this retention probability with monotonically converging functions (‘exp.’, ‘reci.’, ‘log.’; see Methods) as posited by the convergent drift hypothesis, consistently outperformed the constant function model of the equal-rate hypothesis across all maps, with the ‘reci.’ function showing the highest fit (r^2^ = 0.947 ± 0.048; Figure 5E). The fitted asymptotic values for ‘reci.’ function approached one (0.96 ± 0.06; Figure 5F), indicating a decreasing and converging drift rate within the hippocampal dCA1, indicating a transformative stability over time. To investigate whether the trend in retention probability is learning-related, we calculated the retention probabilities for place fields formed in each session, either during the R phase (i.e. session 1 to 3) or S phase (remaining sessions). Most comparisons revealed no significant differences (Figures S17A-F), suggesting that this trend is a learning-independent feature.

In addition to A-to-I transitions, we analyzed the recovery of fields that had previously become inactive (Figures 3A, 3F and 3G). We assessed the conditional recovery probability (*P*_*I*→*A*_), reflecting the likelihood of an inactive field becoming active again given its silent duration (S_t_). this probability significantly declined with longer dormancy periods (from S_t_ = 1 to 8, MA: 31.3 ± 3.2% to 3.3 ± 0.8%, Paired t test P = 9 × 10^−6^, df = 5; MB: 30.2 ± 2.4% to 2.8 ± 0.7%, P = 0.0005, df = 3; Figure 5G and 5H). Despite the decline, recovery probabilities remained significantly above chance level for up to 8 sessions. Similar trends were observed across all other spatial maps (from S_t_ = 1 to 5, MAf: 26.3 ± 5.2% to 4.6 ± 2.7%, P = 0.002; MAb: 25.4 ± 6.1% to 4.0 ± 1.6%, P = 0.007; HPf: 23.8 ± 3.2% to 5.6 ± 1.1%, P = 0.002; MAb: 22.4 ± 2.1% to 3.1 ± 0.7%, P = 0.0004; df = 3; Figure S16B). The declining trend in recovery probability is well-described by the Kohlrausch-Williams-Watts function, a stretched exponential decay function with a slower convergence rate. Similarly, the recovery probability is also learning-independent (Figures S17G-L). This analysis allowed us to define the Single Field Evolution Rule (SFER): the longer a field remains active, the more likely it is to continue being active; conversely, the longer a field remains inactive, the less likely it is to recover.

### Kinetic models with SFER account for the growing cross-day stability in dCA1

SFER delineates the rules governing state transitions for individual place fields. To assess whether SFER alone could explain the relationship between dynamism and stability at the ensemble level, we simulated two classes of kinetic models: the Equal-Rate Drift Model (EDM) and the Convergent Drift Model (CDM), based on distinct trends in field retention probabilities: EDMs assume a constant retention probability, CDMs integrate the dynamically decreasing probabilities predicted by SFER (see Methods, Figure 5I). We evaluated these models by comparing the dynamics of the proportion of super stable fields (defined as fields active for θ sessions) (Figures 5J-P) and the fraction of survival fields across sessions (Figures 5Q-W) with actual data. In EDMs, the proportion of super stable fields remains constant (Figures 5J, 5K and S18A), contrasting with CDMs where it increases over time (Figures 5L and S18B), mirroring the growing stability observed in our data (Figures 5M and 5N). This increase in super stable place fields indicates a rising stability of representation, aligning with previous findings^34^. The differentiation in model performance, particularly the location of breakpoints, was more accurately depicted by the SFER-based functions, with the ‘reci.’ function showing the best fit (see Methods; Figures 5O and 5P). The fraction of survival fields, defined as the proportion of fields that withstand daily drift over a session interval Δt (Figure 5Q), demonstrated exponential decay in EDMs, irrespective of the starting measurement point (Figure 5R and S18C). In contrast, CDMs revealed cumulative effects, with the survival fraction increasing based on the initial measuring point (Figures 5S and S18D). This cumulative effect, indicative of extended average lifespans of individual fields, suggests an increasing stability in the dCA1 representations. This finding aligns with our empirical observations across both maze configurations (Figures 5T-V). Moreover, SFER-predicted functions closely matched the observed cumulative effects across all six spatial maps (Figure 5W), underscoring that kinetic models informed by SFER aptly capture the diminishing dynamism and escalating stability at the ensemble level.

## Discussion

In this study, we first confirmed that the multi-field place cells recorded in our settings displayed canonical statistical structures, consistent with previous findings. By employing multiple analytical tools, we then revealed that the properties and stability of sibling fields are largely independent, despite exhibiting subtle but significant coordination. Finally, by leveraging the advantages of PCmf, we discovered that the longitudinal dynamics of individual place fields are governed by environmental- and learning-independent statistical rules, which we referred to as the Single Field Evolution Rule (SFER), providing a quantitative description.

Discussions about the long-term dynamics of place fields have typically focused on two main aspects: the formation and stabilization of place fields. While our findings primarily anchor into the stabilization process, it is necessary to contextualize them within a broader framework. From a computational perspective, an initial configuration of HPC neuronal representation is rapidly established across the entorhinal-hippocampal circuits^9,48^. A Gamma-Poisson model^12,42^ assumes the formation of this initial representation as a spatial-Poisson process based on the heterogeneous neuronal propensity of dCA1 pyramidal neurons, a concept associated with neuronal excitability. Our dataset, according to the presented analysis, conforms to the Gamma-Poisson model despite differing in the mean neuronal propensity measured in the complex mazes (Figures S7 and S8). Parallel studies on memory engrams^49^ and subthreshold activity during field formation^15–17,20,50–53^ converge to indicate the central role of neuronal excitability in the rapid establishment of HPC representation. Additionally, neuronal plasticity, such as BTSP^16^, contributes significantly to the generation and rapid modification of at least some place fields, although it remains contentious whether these mechanisms account for the formation of all place fields^15,20^. Place cells in novel environments have long been known to be less stable^3,19^, less precisely tuned^19^, and less distinguishable across different environments, consistent with our dataset (Figure S3), suggesting that the initial neuronal representation is provisional and requires further reorganization and refinement.

Cross-day RD is possibly a manifestation of this memory refinement process, during which mechanisms on the cellular level and synaptic level are both required to function in their respective manners and together reconfigure the overall neural network over time. For instance, the expression level of immediate-early genes (e.g., Fos) has been reported to affect the cross-day stability of place fields in pyramidal neurons^40^. To elucidate the underlying mechanisms of cross-day RD, it is essential to determine which mechanisms play a major role. Cellular-level factors, from genetic expressions to day-to-day fluctuations in cellular properties, nonspecifically impact all sibling fields of a neuron, thereby synchronizing their evolutionary trajectories, either enhancing or weakening them together. Conversely, synaptic-level mechanisms are field-specific, resulting in highly diverse fates for sibling fields, although they are expressed by the same neuron. Our analysis confirmed significant coordination (i.e., some interdependence) among the representational evolution of sibling fields (Figures 4A-F, S11 and S12) and a preference for synchronized change (Figures 4G-L and S13-15), indicating the presence of nonspecific factors affecting sibling field stability. However, the ubiquitous variation in the lifespans of sibling fields (Figures 3A and S10) and the low degree of coordination (∼5%) among them (Figures S11C and S11D) provide compelling evidence that field-specific mechanisms are of major importance. This finding aligns with long-standing computational theories on the relationship between synaptic mechanisms and mnemonic functions, suggesting a potential framework for understanding the long-term dynamics of RD at the single-field level.

Investigation of PCmf allows us to reliably capture the switches of place field states and accurately calculate conditional probabilities, given the expanded number of field samples provided by PCmf (Figure 5), laying a solid foundation for the discovery of the Single Field Evolution Rule (SFER). SFER exhibits several compelling properties that collectively elucidate the dynamics of cross-day RD in the dCA1: 1) SFER is a field-specific rule. Given the high independence of fields’ evolution, SFER is adequate to approximately predict the evolutionary trajectory of individual fields, although the exact probability values are averaged over all field samples. 2) SFER applies regardless of whether a field is formed during an animal’s novel states (training sessions 1 to 3) or familiar states (training sessions ≥ 4) (Figure S17), suggesting that learning-related neuronal plasticity like BTSP alone cannot conceivably explain SFER. 3) SFER operates on a cross-day timescale, displaying a slow dynamic far beyond the rapid learning time window (Figures 1D-G and S1). 4) SFER remains consistent across different behavioral paradigms, environments, and movement directions, reflecting inherent network properties in the dCA1 (Figures 5D-H and S16). 5) SFER inherently defines a convergently diminished drift rate and growing stability, capturing long-term features observed at the ensemble level. It provides a coherent explanation for the highly varied lifespans of place fields, such as why only a small fraction persist long-term, aligning with findings from both electrophysiological recordings and calcium imaging studies^8,10,11,13,54^ (Figures 5J-P). Notably, our studies in complex mazes show a substantial number of place fields with lifespans exceeding 35 days (394 fields in Maze A, from 2 mice over 36 days) (Figure 3B), a level rarely observed in 1.5-m linear tracks^11^. This could be attributed to the greater denominator number of fields and the heightened demands for reliable spatial coding in complex mazes^55^. It dismisses the idea that two separate groups of place fields in dCA1, defined by their lifespans, are responsible for transient and persistent coding respectively^34^. SFER also explains how the average lifespan of place fields gradually lengthens, a key feature for representational stabilization (Figures 5J-W). Its convergent dynamics resonate with recent computational frameworks proposing cross-day RD as a consequence of continuous representation refinement^56,57^, a distinct dynamics compared to RD observed in other brain regions^31,58^, suggesting that SFER is confined to the entorhinal-hippocampal circuit and results from characteristics unique to these circuits.

However, it is essential to emphasize several considerations regarding SFER: 1) The validation of SFER is currently limited to a two-month period, and its applicability beyond this timeframe remains uncertain. It is unlikely for the retention probability to eventually reach 100%; it is more plausible that it will plateau, allowing a baseline level of drift due to the slowing changing upstream inputs^9,22,23,48^ and assumed noisy synapses^29^. Nonetheless, our estimation indicates that the baseline drift rate, if present, is very limited (Figures 5C, 5D and S16). 2) The efficacy of SFER has only been tested in environments that remain unchanged and are regularly revisited with intervals of 24 and 48-72 hours, under consistent experimental paradigms. Although previous studies have shown that cross-day RD is unaffected by short intervals (2 or 4 days) but sensitive to the number of visits to the environment^13^, longer intervals (e.g., 10 days or more) can cause significant RD^11^. The drift rate in the piriform cortex has also been reported to depend on the intervals^26^. Future studies are necessary to address the impact that passive elapse of time has on SFER.

Several features are of potential importance for future studies to screen the mechanisms governing SFER. 1) It should be associated with synaptic plasticity at the CA3-to-CA1 projections^9^, as neuronal representations in upstream regions, such as the dentate gyrus^9^ and CA3^9,48^, have demonstrated greater stability compared to the highly dynamic representations in dCA1. Long-term recordings from dCA1 perisomatic dendrites have also identified recurrent spines, providing an anatomical basis for the recovery of place fields^21^. Consequently, further research linking SFER to CA3-to-CA1 synaptic plasticity is crucial to unraveling the underlying principles of SFER. 2) It should bidirectionally affect, either strengthen or weaken, the stability of place fields. Although BTSP has the capacity to bidirectionally adjust synaptic strengths^59^, this mechanism is more likely related to mnemonic processes that profoundly reconfigure the HPC network during rest or sleep^60– 62^. One reason is that SFER is observed on a cross-day timescale, while BTSP functions primarily on a behavioral timescale. Another reason is that mechanisms such as memory consolidation and reconsolidation can display bidirectional effects. For instance, memory becomes liable^63,64^ or strengthened^65^ during reconsolidation, mirroring the large-scale disappearance of place fields and the enhancement of retention probabilities. 3) Although the plasticity occurs on a cross-day timescale rather than a behavioral timescale, such mechanisms should be experience-driven. It is already known that experience, specifically the number of visits to an environment, triggers the modulation of dCA1 spatial representation^13^, and SFER is also demonstrated based on the number of visits (Figures 5C-D and 5G-H). The reactivation of memory driven by revisits significantly affects the storage of memory itself^63–65^, although more research is required to determine if reconsolidation genuinely drives RD in the hippocampus. 4) It should differentially affect place fields with distinct retained durations to result in the convergently growing retention probability (Figures 5C and 5D). Memory must be selected for long-term storage, with more valuable experiences, such as those related to rewards^66–68^, being more likely to be stabilized. From a computational perspective, RD delineates the process of neuronal representations optimizing toward an optimal solution^31^, which theoretically exhibits convergent dynamics^56^. Therefore, it is possible that SFER reflects the innate property of this optimization, with the underlying neural mechanisms serving to implement it.

## Acknowledgements

This work was supported by the National Key R&D Program of China (2019YFA0802400). Open project for Chenglin Miao, State key laboratory of Membrane biology. This work is supported (funded) by Qidong-SLS Innovation Fund, the Lingang Laboratory, Grant No. LG-TKN-202204-01 and Beijing Government Outstanding Youth Scientist for Chenglin Miao. We thank the National Center for Protein Sciences at Peking University in Beijing, China, for assistance with imaging and histology.

## Author contributions

Q.C, C.M. designed research, C.C, S.Y, S.C, collected the data, S.Y, C.C, S.C, A.L, X.Z, Y.Y, Y.W, analyzed the data. Q.C, C.M supervised the project, S.Y, C.C, C.M. wrote the paper with input from all authors.

## Declaration of interests

The authors declare no competing interests.

## Supplemental information

Document S1. Figures S1–S17

## Materials and Methods

### Animals

Mice (C57BL/6N, male and female) were group-housed in a controlled environment under a reverse 12-hour light/dark cycle (9:00-21:00). Prior to undergoing surgery, they had ad libitum access to standard laboratory chow, water, and a running wheel for enrichment. Surgeries for implanting the necessary devices were conducted at approximately 10-14 weeks of age. Post-surgery, mice were singly housed to prevent implant damage and facilitate recovery. We expressed GCaMP6f in the hippocampal dCA1 region of four mice (IDs: #10209, #10212 (female), #10224, #10227), which were utilized for all behavioral paradigms and imaging studies. Two additional mice (IDs: #11092, #11095) expressing GCaMP6s were trained exclusively for the maze navigation paradigm; their imaging data were used solely to validate the Poisson distribution of field numbers. Another mouse (#11094, female) was trained only in Stage 1 of the maze navigation paradigm and was therefore included only in the behavioral analysis. All procedures were approved by the Institutional Animal Care and Use Committee at Peking University and adhered to ethical guidelines for animal research.

### Surgery

#### Virus injection

Virus Injection: Animals were initially anesthetized with 5% isoflurane in air in a chamber and subsequently placed in a stereotactic apparatus (Kopf Instruments) with 1-2% isoflurane. The skin was sterilized with betadine, and erythromycin ointment was applied to protect the eyes. The fur over the skull was removed, and the head window was carefully cleared. Using a stereotactic drill, holes were made at the designated injection sites (AP: - 1.82 mm, ML: -1.25 mm; AP: -2.5 mm, ML: -2.28 mm). rAAV-hsyn-GCaMP6f-SPRE-hGH-pA (titer: 1×10^12^) was injected via a beveled steel needle, positioned 1.4 mm below the dura, at a rate of 100 nL/min, delivering 200-250 nL at each site. The needle was retained in situ for an additional 10 minutes to facilitate even distribution of the viral vector. The incision was then sutured, and the mice were allowed to recover for 2-3 weeks to ensure optimal viral expression.

#### Lens implant

Anesthetic procedures were identical to those described for virus injection. The skull was re-exposed, and a craniotomy of approximately 2.0 mm in diameter was performed with a hand-held drill, centered over the previous virus injection sites. After removing the skull, the brain tissue was exposed. The cortex was gently aspirated using a blunt needle attached to a vacuum pump until the stripy corpus callosum became visible. The first layer of the corpus callosum was carefully removed until the fiber direction changed and the tissue appeared transparent. During aspiration, 0.9% saline was continuously applied. A hemostatic sponge was then placed in the cavity until significant bleeding ceased. Three set screws were affixed to the skull for added stability and attachment of the cement. The GrinTech lens, attached to the UCLA miniscope and connected to the DAQ box and computer, was slowly lowered into the craniotomy cavity until clear cell signals or blood vessels were visible. The optimal viewing position was determined, and the lens was secured in place with Metabond dental cement. A tube cap was placed over the lens for protection. Postoperatively, ceftriaxone sodium (1.25 mg/kg) was administered intraperitoneally to the animals for 3-7 days. The animals were then returned to their home cages for a recovery period of 1-2 weeks.

#### Baseplate

The miniscope was connected to the DAQ software and affixed to the aluminum baseplate at the bottom of the miniscope. The miniscope was then mounted above the lens, and its position and focal length were adjusted to obtain the optimal field of view. The miniscope was secured to the holder, allowing the baseplate to be cemented just above the optimal viewing position. After the cement had dried, the miniscope was detached, leaving the baseplate attached to the animal’s head for subsequent reattachment and imaging sessions. Finally, a Lego block was attached as a protective cap for the lens.

#### Histology

Mice were sedated intraperitoneally with avertin. Subsequently, they were perfused transcardially with 0.9% saline followed by 4% paraformaldehyde (PFA). Brains were extracted and postfixed in 4% PFA at 4°C for 24 hours. Afterwards, the fixed brains were transferred to 30% sucrose for dehydration. The tissues were rapidly frozen using instant freeze spray and sectioned into 40 μm slices. Brain sections were mounted directly onto slides and washed with PBS three times for 5 minutes each at room temperature. Sections were blocked for at least 1 hour in PBS containing 10% goat serum and 0.1% Triton X-100. They were then incubated overnight at 4°C with the primary antibody Chicken-anti-GFP IgY (1:1000, Invitrogen, A10262) to label GCaMP6f-positive neurons. After primary antibody incubation, sections were rinsed three times in PBS and incubated for 2 hours with the secondary antibody Alexa Fluor 488 Goat-anti-Chicken IgG (1:1000, Invitrogen, A11039) and DAPI (1:1000, Sigma, D9542) for nuclear staining. Slides were then sealed with a coverslip using Fluoromount-G (B2623-NJ03, SouthernBiotech). Finally, the sections were imaged using the Olympus Virtual Slide System (VS120-S6-W).

### Behavioral Training

Mice were singly housed with a running wheel and Lego in their home cage post-surgery, with free access to food and water until three days before training. During training, food was restricted to maintain their weight at ∼80% of their baseline to ensure motivation.

#### Open Field

The open field arena is squared (1m × 1m × 0.5m, length × width × height). The environment was marked by distinct black shapes and stripes on four white walls, illuminated by dim white light, with three sides surrounded by black curtains. The recording computer was placed on the left wall side. Mice were anesthetized to attach the miniscope and allowed to recover before being placed in the center of the arena. Recording began when the mice actively explored the environment. Crumbs of butter cookies, sometimes mixed with chocolate, were used as incentives. A helium balloon was attached to balance the weight of the recording cable and miniscope. Sessions lasted ∼30 minutes or until the mice covered the entire area. Mice were subjected to a pre-training phase within open field for 20∼27 30-min sessions over 3 weeks, to habituate movement with miniscope and stabilize the imaging quality. During the Stage 1 and Stage 2, mice were exposed to this open field for two 30-min sessions before and after the maze session(s), respectively. This accompanied open field random exploration task was conducted for 26 sessions over 26 days in each Stage.

#### Maze Navigation Paradigm (MNP)

Both mazes are squared (0.96 m × 0.96 m × 0.5 m, length × width × height). Maze A was constructed as a 12 × 12 lane grid with 8 cm wide lanes, featuring a total path length of 8.88 m and 17 decision points. The walls were adorned with distinct shapes and stripes to serve as distal cues. Maze B, while identical in size and the number of decision points, had a shorter correct path measuring 8.08 m. The environmental setup, including visual cues and lighting, remained consistent for both mazes.

Training was divided into two stages, each comprising 13 blocks spread over 26-29 days. In Stage 1, a block consisted of three successive sessions: open field, Maze A, and open field again. This sequence aimed to familiarize mice with Maze A while reinforcing their acclimation to the open field. In Stage 2, a Maze B session was added after Maze A, introducing a novel challenge while retaining familiarity with the previous environments.

A ‘lap’ in maze navigation is defined as the interval between a mouse entering and exiting the maze. Mice were required to complete at least 5 laps in the first session of both mazes and at least 10 laps in subsequent sessions. Each maze session lasted approximately 30 minutes, except for the initial maze sessions, which extended over an hour to accommodate the time-consuming process of locating the exit for the first time. A reward box containing cookies was placed outside the exit, and mice were manually transferred from the reward box back to the entry for subsequent laps. Laps without any incorrect decisions or turn-around events were labeled as ‘perfect’ laps.

### Reversed Maze Paradigm (RMP)

The environmental setup was consistent with previous settings. Mice previously familiarized with forward navigation in Maze A were trained to traverse back and forth between the entry and exit. A lap, whether forward or backward, was defined similarly. Each 30-minute session comprised 21-48 laps in total, with 10-24 laps in each direction. Training lasted for 7 to 12 sessions over 7 to 15 days, with 2 mice completing the shorter duration and 2 mice completing the longer duration.

### Hairpin Maze Paradigm (HMP)

We implemented a linear navigation task in a hairpin maze, designed to exclude decision-making while retaining the characteristics of the complex mazes. This included identical dimensions, material, and wall height, but with a different arrangement of the internal barriers. The total path length within the maze was 11.52 meters. Environmental cues and the setup of the room mirrored those used in the maze training sessions. Rewards were strategically placed at both ends of the track. Each 30-minute training session comprised 15-48 laps, spanning 7 sessions conducted over a week.

The environment setups were cleaned carefully by 75% ethanol every time after ending a recording session.

### Data analysis

#### Behavioral data

Animal behavior in our experimental setups was captured using a top-view webcam at a frame rate of 20 Hz. We programmed to automatically concatenate multiple videos into one integrated file and provide a user-friendly graphical interface for manual labeling of the start and end frames of each lap.

For tracking mouse positions, we employed DeepLabCut^69,70^ to track the mice positions. The model was trained using an automatic extraction method that down-sampled the videos with a k-means algorithm and a sliding window of 25 frames. We annotated four points along the rostro-caudal axis of the mice: the head (marked with a red LED light), the neck, the back, and the tail. The neck point was chosen to represent the animal’s position due to its proximity to the centroid and comparatively lower tracking noise.

The behavior data underwent several processing steps for analysis: 1) Removal of all NaN and erroneous values, with errors defined as points with a transient speed exceeding 100 cm/s. 2) Retention of behavioral data only within defined laps, discarding inter-lap information. 3) Rectification of visual perspectives caused by filming angles through an affine transformation, which required manual identification of the four corners of the mazes or open fields to create a transformation matrix using the getPerspectiveTransform() and perspectiveTransform() functions from the OpenCV package. 4) Binning of the transformed trajectory into 2 cm bins for each dimension. 5) Application of a cross-wall correction to adjust or remove points that cross walls, which is rarely necessary due to the well-lit environments and strong visual contrast between the mice and the ground. Furthermore, the neck’s position crosses walls less frequently than the head’s position, and meticulous labeling of the corners minimizes cross-wall points.

To linearize the correct paths of our mazes, we divided the correct paths into large square bins (111 for Maze A, 101 for Maze B, and 144 for hairpin mazes), each with an 8 cm edge. These bins are spatially organized, with each containing 16 smaller 2 cm x 2 cm bins (Figures 1A and S6J). This approach treats all small bins within a large bin as a single unit, resulting in approximate path lengths of 8.9m for Maze A, 8.1m for Maze B, and 11.5m for the hairpin maze. For visualization, jitter is added to the position of each small bin relative to its corresponding large bin. This approach doesn’t perfectly capture the trajectory that is highly curved and optimized for navigation, which measures approximately 6.3m for Maze A, 5.7m for Maze B, and 10.3m for the hairpin maze.

#### Behavioral indexes

To evaluate behavioral performance, we defined three basic indexes for each lap: 1) the duration spent by the mouse, 2) the distance traveled to complete the lap, and 3) the mouse’s mean speed during the lap. For each session, we defined additional indexes: 1) the correct decision rate, defined as the ratio of correct decisions at decision points to the total number of decision points encountered while moving forward, and 2) occupation time on correct or incorrect paths, calculated as the total time spent on each path divided by the number of bins in that path.

The behavioral learning progress (LP) in each session is determined by dividing the current progress by the total progress. The total progress is calculated by taking the absolute value of the difference between the maximum and minimum values of the animal’s behavioral index.

#### One-photon calcium imaging and signal preprocessing

The UCLA miniaturized micro-endoscope (Miniscope V3)^71^ was custom-built following a previous design. Calcium imaging data collected by the miniscope were synchronized with behavioral data from a top-view webcam at 20 Hz using Miniscope DAQ software. Downsampled AVI videos underwent motion correction with NoRMCore^72^. CNMF-E^73^ was employed to delineate regions of interest (ROIs), extract the dF/F signals of each neuron, and deconvolve calcium signals. Spatial (ssub) and temporal (tsub) downsampling factors were set to 1 and 5, respectively. Neuron diameter (gSize) was constrained to 15 pixels, and a Gaussian kernel (gSig) with a width of 3 pixels was used for data filtering. A ring radius (ring_radius) of 18 was applied in our background model (bg_model). ROIs with signal-to-noise ratios above 5-fold (smin=-5) and minimum peak-to-noise ratios of 8 (min_pnr) were selected. Data from all sessions were included in the analysis, with the exception of Stage 1, Session 6 in Maze A for mouse #10212. This session was excluded due to the mouse’s abnormal behavioral performance, characterized by physical weakness and evident disengagement from the navigation task.

#### Signal Processing

Deconvolved signals were binarized, with frames having deconvolved values greater than three times the standard deviation of the deconvolved signals considered as calcium events. Frames with corresponding behavioral speeds below 2.5 cm/s were discarded to ensure activity detection during active movement. Only within-lap neural activities were retained for analysis, excluding inter-lap signals. Unless stated otherwise, analysis in the MNP paradigm included all within-lap calcium activities. To account for potential directional tuning of spatial maps and occasional backward movement during navigation, we also isolated purely forward spatial maps on the correct path by removing calcium events detected during backward movement or on incorrect paths. This treatment will be explicitly mentioned when applied. Signals recorded during the RMP and HMP were categorized by movement direction (forward or backward) to analyze direction-specific neural activity. Additionally, neural activities from incorrect paths in Maze A during RMP were excluded from the analysis. In the all paradigms, neurons with fewer than 10 calcium events were excluded from analysis but counted in the total neuron number.

#### Calculation of spatial rate maps

We computed calcium event rate maps to visualize neural activity. Our mazes were divided into 48×48 square spatial bins (numbered 1 to 2304), each with a side length of 2 cm. We calculated the occupation time spent by mice in each bin, setting bins with occupation times under 50 ms as NaN to prevent abnormally high event rates. All NaN values of this raw event rate maps of *N* neurons, *R*_*r*_ ∈ ℝ^*N*×2304^, were later set as 0 for smoothing purposes. For smoothing, we employed a modified Gaussian kernel that accounts for the internal walls of the mazes. The Gaussian smoothing matrix *W* ∈ ℝ^2304×2304^ is defined such that its entry at row *i* and column *j, w*_*i,j*_, is given by:

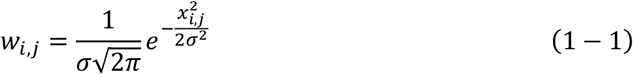

where *σ* represents the standard deviation, set as 2, and *x*_*i,j*_ denotes the shortest theoretical distance between bin *i* and *j* that does not cross any internal walls. The smoothing matrix *W* was normalized across columns to ensure that the sum of weights for each bin equals 1. The smoothed event rate map *R*_*s*_ was computed as:

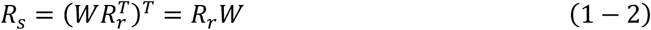

All visualized smoothed event rate maps displayed in the main figures were purely forward or backward, based on the direction of the mouse’s movement in the maze. The linearized rate map utilized 12×12 bins, each 8 cm wide, aligned with the maze’s correct path as described in the ‘behavioral data’ section. Bins were reordered according to their sequence along the correct path for clearer visualization and analysis of neural activity distribution.

#### Spatial Information

The calculation of spatial information (SI) follows the methodology established in previous research^74^. Briefly, SI is computed using the formula:

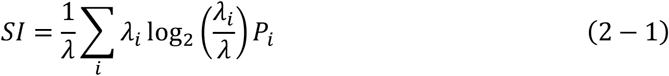

where *λ* is the mean firing rate of a neuron across all bins, *λ*_*i*_ is the firing rate of the neuron in bin *i*, and *P*_*i*_ represents the fraction of time mice spent in bin *i*. This calculation quantifies the amount of spatial information conveyed by the neurons’ firing pattern.

#### Identification of place cells

Neurons were classified as place cells when passing all the three shuffle tests: 1) Temporal Shuffle Test, which dissociates calcium events’ timestamps from the mouse’s positions, 2) Inter-Event Interval Shuffle Test, which rearranges calcium events by shuffling their inter-event intervals^75^. 3) Random Shuffle Test: which randomly rearranges the calcium events. Each test was repeated 1000 times, and a neuron passed if its real SI exceeded 95% of the shuffle test SI values. Stability criteria were not applied to avoid limiting the exploration of dynamic neural representations, yet neurons meeting our criteria showed sufficient stability at the population level within sessions (Figures S3D-F).

#### Within-session stability

To assess neuronal within-session stability, we employed half-half and odd-even correlation methods. The former compares smoothed event rate maps between the session’s first and second halves, determined by lap count for equal position coverage. The latter correlates maps from odd and even laps. Both serve as reliable stability indicators, with half-half correlations typically lower, possibly due to the time needed for place cells to reestablish spatial representations or new field formations in later laps. Unless specified, we discuss stability using half-half correlations.

#### Identification of place field

We defined a place field with multiple requirements. 1) Spatial bins with a smoothed event rate above 0.4 Hz were considered as candidates for place fields. 2) A minimum of 10 calcium events were required to be detected on the bins constituting a place field to avoid inclusion due to occasional fluctuations. 3) To ensure reliable encoding of positions, calcium events must be detected in at least five laps within a maze or during five separate visits in an open field, with a minimum interval of 15 seconds between each visit. 4) We addressed closely spaced place fields to avoid miscounting field numbers and overestimating average field sizes. We split a place field with multiple peaks into multiple fields if the fraction of the inter-field rate to the peak rate was lower than 0.2 (split threshold).

These criteria are referred to as ‘rigorous criteria’, which may miss some weak and dynamic place fields due to the high requirement for the number of calcium events and be less sensitive to separate two spatially close and occasionally merged place fields due to the low split threshold. For our long-term studies on the evolution of place fields, which requires minimizing false-negative results as much as possible for reliable tracking of place field states, we also applied an alternative set of criteria, i.e., the ‘loose criteria’: a 0.2 Hz threshold and 5 calcium events across at least 5 laps or visits, and the split threshold was set as 0.5. Most of our analysis adopted the loose criteria, and our analysis confirmed that place fields identified by loose criteria display canonic statistical structure at the ensemble level.

#### Within-field stability

To analyze place cells with multiple fields, we focus on within-field stability rather than overall neuronal representation. This is calculated as the Pearson correlation (either half-half or odd-even) of event rates within place field bins. For fields smaller than 16 bins, neighboring bins are included to ensure a minimum of 16 bins for the correlation calculation, reducing variance in the correlation values.

#### Naïve Bayesian classifier

To reconstruct the positions of mice, denoted by spatial bin IDs, from their neural activity, we employed a modified naive Bayesian classifier. We have a Δ*F*/*F* activity matrix:

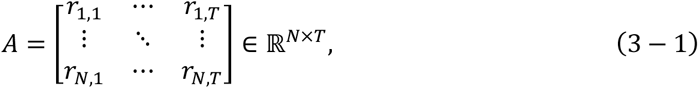

where *N* represents the number of neurons used for decoding, and *T* is the total number of frames. The entry of *A* at row *i* and column *t*, i.e.,

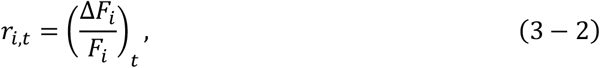

is the Δ*F*/*F* value of neuron *i* ∈ {1, 2, …, *N*} at temporal frame *t*. We applied neuron-specific indicator functions to binarize ***r***_*t*_ and obtain a binarized population vector.

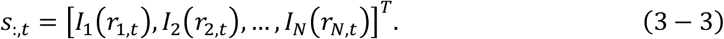

The indicator function of neuron *i* is defined as:

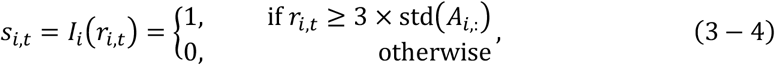

with std(*x*) returning the standard deviation of vector *x*. Each *s*_:,*t*_ corresponds to a unique spatial bin ID, denoted as *y*_*t*_ ∈ {1, …, *Y*}, where *Y* is the number of spatial bins and is 2304 in our content. For each neuron *i* we could compute its firing probability at each position *j*, namely a normalized tuning curve^76^, yielding an *N* × *Y* matrix *P* whose element at row *i* and column *j* is *P*(*s* = 1|*y* = *y*_*j*_). This matrix was smoothed by a modified 2D Gaussian smoothing matrix, *A*, which have described in formula (1 − 2):

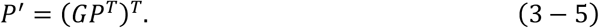

Traditional naive Bayesian classifiers typically minimize a zero-one loss function. However, in our case, we consider the loss as a function of the distances between predicted and real bins. Our modified naive Bayesian classifier minimizes the expected risk function:

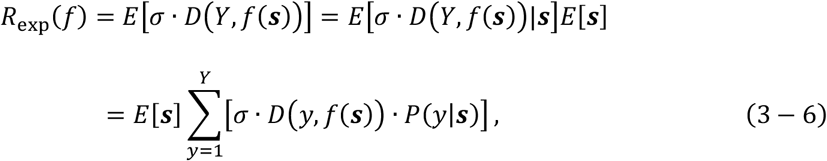

where *D*(*a, b*) is a distance function, *σ* is a scale parameter, *f*(***s***) is the decision function. This leads to the decision rule:

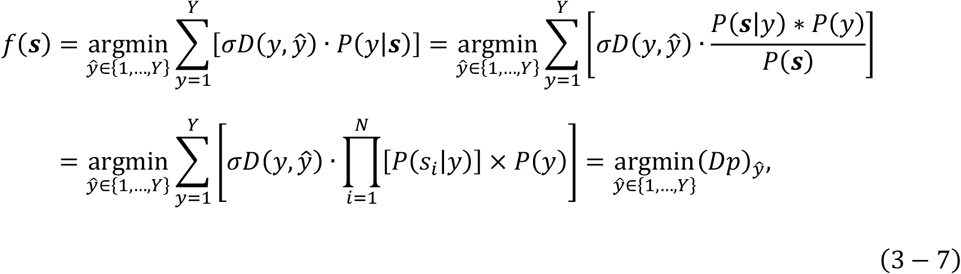

where *ŷ* is the predicted bin ID, *s*_*i*_ is the binarized value of neuron *i, D* ∈ ℝ^*Y*×*Y*^ is a matrix containing the scaled distances between bins, and *p* ∈ ℝ^*Y*^ is a vector calculated from *P*′:

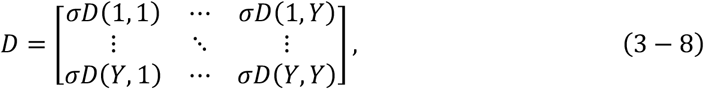

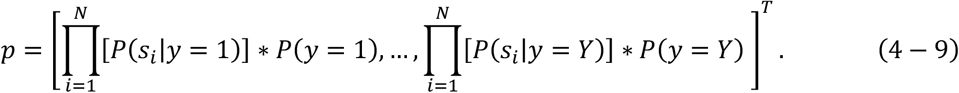

Our modified classifier decodes the animal’s position for each recording frame. The median absolute error (unit: cm) is used to assess the decoding results for each lap. We trained the classifier using *L*′ ′ laps (approximately 60% of all *L* laps) and performed lap-wise decoding. Basically, we trained the decoder with the last *L′* laps data to decode the first *L* − *L* ′ laps. To decode positions in the last *L*′ laps, we used a rolling approach, training the decoder with the remaining laps from the last *L*′ + 1 laps for each lap that required decoding.

#### Log-normal distribution and gamma distribution

Field sizes, measured by the number of bins, were fitted with either a Log-normal distribution or a gamma distribution as reported in a prior study^43^. The sizes of fields collected across familiar sessions (MA: Stage 1 S≥4 and Stage 2 S≥1; MB: Stage 2 S≥4) in the maze navigation paradigm and all the sessions in reversed maze paradigm and hairpin maze paradigm were examined together using Lilliefors-corrected Kolmogorov– Smirnov tests. Given the large number of field, which exceeded the validity range of the KS test, we conducted bootstrap resampling to down-sample the number of fields for goodness-of-fit test to 1621, a level reported in the prior study^43^, to mitigate over-sensitivity of KS test. The Log-normal distributions were fitted using the scipy.stats.lognorm.fit function, and the gamma distributions were fitted using the scipy.stats.gamma.fit function, with the location parameter forcibly set to 0.

#### Gamma-Poisson model for hippocampal multi-field coding

One key aspect of hippocampal multi-field coding is understanding how fields are assigned to individual neurons. Previous research^77^ suggests that the spatial distribution of place fields follows a homogeneous spatial Poisson process with a certain rate *λ* (unit: field per meter), leading to exponentially distributed inter-field intervals. This rate *λ*, indicative of a neuron’s propensity or inertia to form place fields, is assumed to be an innate, environment-independent property of each neuron. If this neuronal propensity is homogeneous or nearly so across the neural ensemble, meaning *λ* is relatively constant, then the number of fields, denoted as *X*, follows a Poisson distribution with rate parameter *λ*:

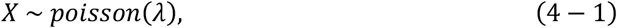

which has a probability mass function (pmf) given by:

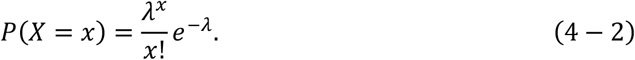

The Gamma-Poisson model was introduced by Rich et al.^77^ to explain an observed negative binomial distribution of place cells’ field number, which significantly deviates from Poisson distribution. This model assumes a highly heterogenous rate parameter *λ* across neurons, with some forming more fields and others fewer.

Unlike an approximately fixed *λ* in a Poisson distribution, the Gamma-Poisson model treats *λ* as a random variable *Λ*, following a gamma distribution with a shape parameter α and a scale parameter *σ*:

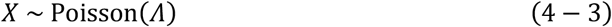

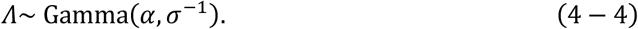

Under this model, *X* is governed by a mixture of Poisson and gamma distributions and ultimately follows a negative binomial distribution characterized by a shape parameter *r* and a scale parameter *p*:

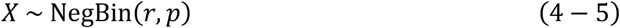

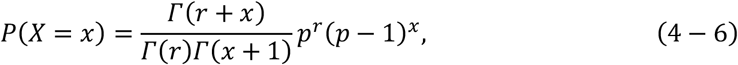

where

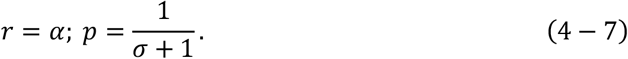

When the shape parameter r is less than 1, the probability mass function (PMF) of a negative binomial distribution monotonically decreases with a long tail as the value of X increases. When r is greater than 1, the PMF has a Poisson-like shape. As the scale parameter σ approaches zero, the heterogeneity of neuronal propensity decreases, and the negative binomial distribution becomes mathematically equivalent to a Poisson distribution. To fit these distributions, we used the ‘scipy.optimize.leastsq’ function for the Poisson and negative binomial distributions, and ‘scipy.stats.gamma.fit’ for the gamma distribution. We also fitted the field number with discrete normal distribution by ‘Scipy.stats.norm.fit’ function. The goodness of fit was assessed by Lilliefors-corrected Kolmogorov-Smirnov test with a bootstrap sample size of 500.

### Estimation of individual neuron’s propensity

To assess potential discrepancies between our observed Poisson-distributed field number and the Gamma-Poisson model, we quantified the distribution of potential heterogeneity of neuronal propensity. Although the real distribution of *λ* is unobservable, it can be approximated using the statistical property of the Poisson distribution:

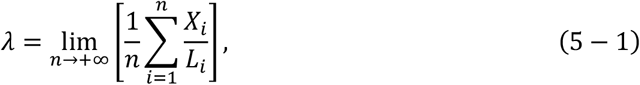

where *X*_*i*_ ∼ Poisson(*λL*_*i*_) for *i* ∈ {1,2, …, *n*}, and *L*_*i*_ is the path length. The mean of a neuron’s field numbers per meter across different spatial maps can approximate its propensity. We used n = 4 spatial maps (MA forward, MA backward, HP forward, HP backward) for estimation. Neurons that were place cells in all four maps were used for estimation. Field numbers were divided by the approximate real trajectory lengths of 6.3 meters in Maze A and 10.3 meters in hairpin mazes to obtain observations of propensity.

We conducted seven experimental blocks over ten days, analyzing neuron pairs from each mouse together. We fitted the estimated values for each mouse with a gamma distribution and assessed their conformity using the one-sample Kolmogorov-Smirnov test. The heterogeneity of the neuronal propensity across the ensemble can be quantified by the coefficient of variation (CV), which assesses the extent of dispersion of a distribution:

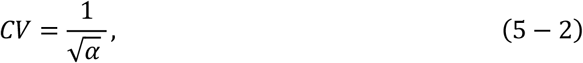

where *α* is the fitted shape parameter of Gamma distribution. The lower CV is, the lower relative variance the neuronal propensity is.

### Lilliefors-corrected Kolmogorov–Smirnov test

The one-sample Kolmogorov–Smirnov (KS) test can be limited when reference distribution parameters are estimated from the sample. The Lilliefors correction^78^ provides a method to mitigate this limitation, leading to more accurate statistical results. The process involves the following steps: 1) Estimate the initial parameters of the reference distribution from the original data and compute the initial KS statistic, *D*_0_, using:

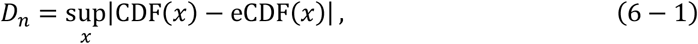

where x represents all possible sample values, CDF(*x*) is the cumulative density function of the reference distribution with fitted parameters, and eCDF(*x*) is the empirical cumulative density function computed from the samples. 2) Generate a bootstrapped sample with size N from the original data^79^ and obtain a new set of parameters param_b_ for the reference distribution from the bootstrapped sample. 3) Generate N random observations from the reference distribution with fitted parameters and obtain a new set of parameters param_r_ for the reference distribution from the randomly generated observation. 4) Compute the KS statistic, D, between the randomly generated observations and the reference distribution with param_r_. 4) Repeat steps 2 and 3, i.e., Monte Carlo simulation, 10,000 times to obtain 10,000 KS statistics, D. 5) Determine the significance level (P-values) as the fraction of random statistics exceeding the real statistic, *D*_0_.

The Lilliefors-corrected approach addresses the limitations of the one-sample KS test when reference distribution parameters are estimated from the sample. Additionally, bootstrap resampling overcomes the over-sensitivity of the KS test in cases of substantial sample sizes, ensuring a more accurate evaluation of distributional conformity.

### Analysis of sibling fields’ properties independence

In our study, we explored the independence of properties within sibling place fields, which are place fields belonging to the same neuron. Our null hypothesis suggests that sibling fields exhibit similar properties, such as stability, size, and firing rate, in the level significant over non-sibling fields. To assess this, we employed three approaches, each offering a different perspective on the hypothesis:

Firstly, we employed a probabilistic approach, drawing from probability theory’s definition of independence, which states that two events *A* and *B* are independent if and only if

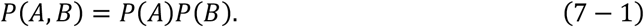

In our context, *A, B* are discrete random variables representing a certain property of place fields, such as field size. This property, denoted as *X*, can take on *n* possible values {*x*_1_, *x*_2_, …, *x*_*n*_} and is measured across a set *V* of all place fields:

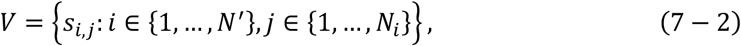

where *s*_*i,j*_ is the value of this property of place field *j* of place cell *i, N*′ is the total number of place cells, *N*_*i*_ is the field number of place cell *i*. We then consider all possible combinations of field pairs, forming a set *S* of paired property:

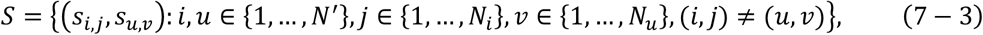

where *s*_*i,j*_, *s*_*u,v*_ ∈ *V* are two values of the given property (e.g., field size) of field *j* of place cell *i* and field *v* of cell *u* respectively, *N*_*i*_ and *N*_*u*_ are the field number of cell *i* and *u*, respectively. We have a related joint probability:

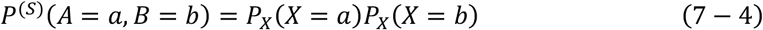

for any *a, b* ∈ {*x*_1_, *x*_2_, …, *x*_*n*_}, where *P*^(*S*)^(*A, B*) is computed from all pairs (*s*_*i,j*_, *s*_*u,v*_) ∈ *S*. Now, we divided this set into two subsets:

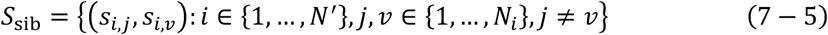

for sibling field pairs and

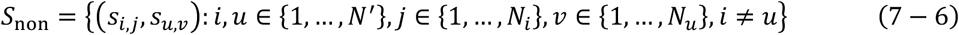

for non-sibling field pairs.

We computed the joint probability functions for each subset, denoted as 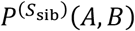 and 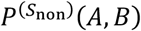 respectively. Generally, the number of elements in *S*_non_will be hundreds to tens of thousands of folders more than that in *S*_sib_, depending on the number of place cells and the average field number. This imbalanced set size will result in distinct accuracy for estimating the joint probability. Therefore, we randomly and non-repeatedly drew *N = min*(|*S*_sib_|, |*S*_non_|) paired properties of field pairs from both sets to construct joint probability matrix 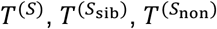 to represent the distributions of properties within these sets:

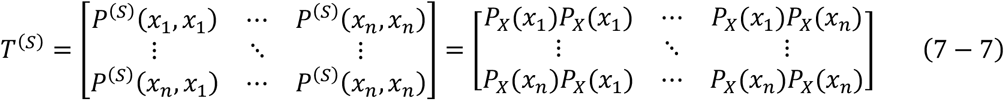

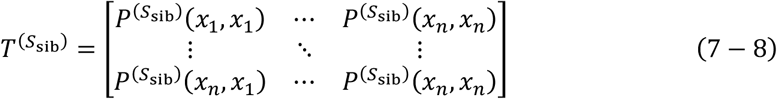

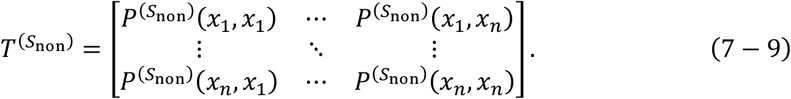

If our null hypothesis holds and there is a tendency in sibling fields’ properties, 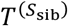 significantly deviate from *T*^(*S*)^ more than the deviations between 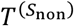 and *T*^(*S*)^. We quantify these deviations using the *χ*^2^ statistic:

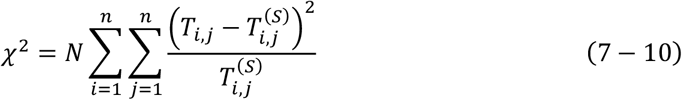

where *T* is either 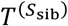 or 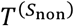. The joint probability matrix and *χ*^2^ statistics are computed with the binned_statistic_2d() and chi2_contingency() functions provided by SciPy.

The second method is based on information theory, which utilizes the concept of mutual information *I*(*A, B*) ^80^ to quantify the dependence between two discrete random variables *A* and *B*:

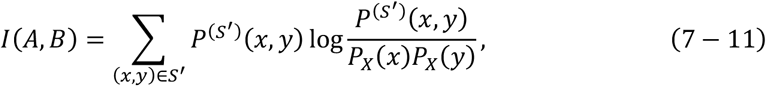

where *S*′ is either *S*_sib_ or *S*_non_, and the summation is applied to *N* values randomly and non-repeatably drawn from both sets to ensure a balanced sample amount for computing the joint probability. We use the function ‘mutual_info_score’ from scikit-learn to compute mutual information^81^.

We must emphasize that these methods are applied to discrete random variables, while some of the field properties are continuous. To accommodate this, we binned all continuous variables into 20 bins to satisfy the assumptions of both methods. This discretization allows us to effectively apply the concepts of mutual information and *χ*^2^ statistics, originally defined for discrete variables, to our continuous field properties by treating each bin as a discrete category.

The third method focuses on the variance of sibling field properties. It assumes that if neurons tend to select place fields with certain properties, and if such a tendency prevails, it will result in a significantly lower standard deviation of sibling field properties than a chance level. The standard deviation among sibling fields forms a vector:

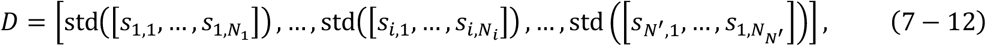

where std(*x*) returns the standard deviation of vector *x, N*′ is the total number of place cells, and *N*_*i*_ is the number of fields of cell *i* ∈ {1, …, *N*′}. Then for each place cell *i*, we randomly draw *N*_*i*_ place fields from *N*_*i*_ other place cells to constitute a group of virtual sibling fields:

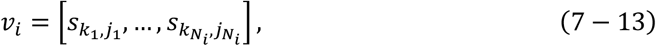

where 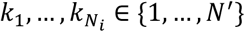 are *N*_*i*_ mutually distinct numbers denoting *N*_*i*_ distinct place cells, and 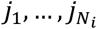 are the indexes of fields drawn from the corresponding neurons. We compute the corresponding standard deviation vector *D*_shuf._ with the same length as the chance level:

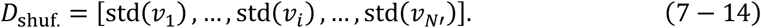

We compare the mean value and distribution between *D* and *D*_shuf_ using the two-sided Paired-t test and the two-sample KS test, respectively. Only place fields with more than 2 fields were included in this analysis.

### Cell registration

We tracked neurons across days and environments using the CellReg algorithm^82^. First, we computed pairwise correlations of the spatial footprint (SFP) for all sessions to be aligned and selected the session with the highest mean correlation as the optimal reference session (*s*_*opt*_). Second, we used the Fiji plugin MultiStackReg^83^ to generate a transformation matrix for affine transformation, correcting linear shifts in the field-of-view (FOV) due to miniscope reattachment. After pre-correction, CellReg was used to register neuron indexes with a P_same threshold of 0.5, micron per pixel set to 2.3, and maximal distance set to 14 microns.

CellReg performs rotation and movement corrections on the SFPs based on a reference SFP. The choice of reference SFP affects alignment results, as neurons may match differently depending on the reference session. To optimize final alignment, we applied a re-match algorithm, named ‘re-match optimization’, based on CellReg’s outputs with distinct reference footprints.

The index map returned by CellReg, denoted as, denoted as *N* ∈ ℝ^*S×Q*^ where *S* is the number of sessions and *Q* is the number of registered neurons, varies accordingly when the SFP alters. We denote the index map with respect to a reference SFP *k* ∈ {1, …, *S*} as *N*^(*k*)^. Each column of 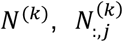 represents a registered neuron *j*. The *t* entries of 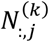 denote its indexes in *S* sessions, respectively. If this neuron was not detected in a session *i* ∈ {1, …, *S*}, then its entry 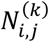 is 0. Now we selected *S*^*′*^ ′distinct SFPs as references. The set of all the SFPs that we chose as references is denoted as *R*, where

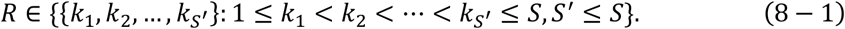

Then we obtained *t*′ distinct index maps:

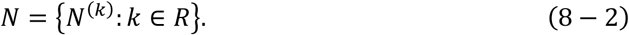

Practically, for the 13-session stage, we selected session 1,3,7,10, 13 as references. For the 26-session stage, we selected session 1, 3, 7, 10, 13, 17, 20, 23, 26 as references. The index map with respect to the optimal reference SFP *k*_*opt*_ is 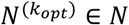. Any further improvements and optimizations were conducted on this index map.

For any registered neuron *j* of 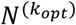, there will be at least one registered neuron *j*^(*k*)^ in any other index map *N*^(*k*)^ such that 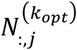 and 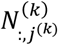 share most identical items at the same positions. These registered neurons form a *S* × *S*′ candidate matrix *C*:

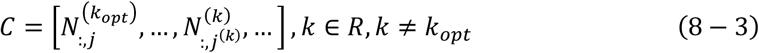

In some positions of the registered neuron, the neuron index will not change despite alterations in the reference SFP, resulting in identical elements across columns. In uncertain positions, neuron indexes highly depend on the reference. Our aim is to find an optimal registered neuron *J*_*opt*_ = [*I*_1_ … *I*_S_]^T^, where 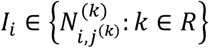 and *T* is the transpose.

For a given candidate matrix *C*, we consider all finite combinations of candidates, forming a set *S*_*J*_:

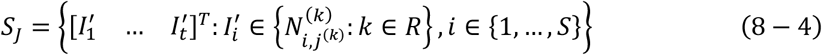

For any 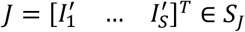, we have a uniquely determined matrix *P* ∈ ℝ^*S×S*^, the likelihood matrix, that defines the pairwise likelihood of neurons being registered:

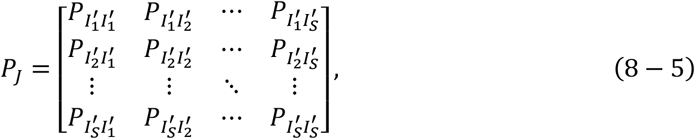

where 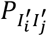 represents the likelihood that the neurons indexed 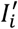 and 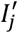 are the same neuron. We define a registered score, *RS*(*J*; *α*), based on the version described by Sheintuch et al., 2017, but with an additional structural item to maximize the number of neurons registered together:

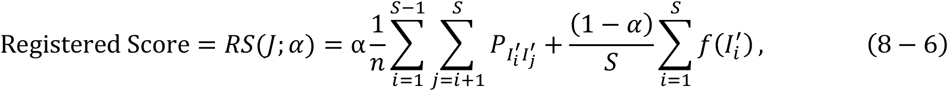

where *α* ∈ [0, 1] is a fraction parameter set as 0.5 in our analysis *n* is the number of non-NaN values in the upper triangle of *P*_*J*_, and *f*(*x*) is an indicator function returning one if the input 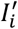 is larger than 0 (indicating it is detected in session *i*) and zero otherwise. This modified version of the registered score balances the need for maximum likelihood that the registered neurons are the same and the need for more neurons to be registered together to provide adequate samples for computing precise probabilities determining the fate of place fields.

As the total number of combinations is generally very large, exceeding what we can compute within a limited time, we used a greedy strategy for optimization. The initial registered neuron *J*^(O)^ = *J*_*opt*_. In iteration *t*, we update all *S* entries of *J*^(*t-*1)^ one by one. For each update on entry 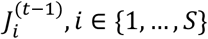, the remaining entries are not changed. The update is conducted to maximize registered scores by replacing 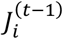 with all the candidates:

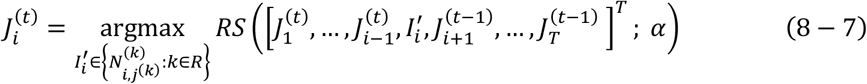

we iteration this process until *J*^(*t+*1)^ *= J*^(*t*)^ or *t* ≥ 20. Only registered neurons with more than 6 non-zero elements are optimized. Additionally, if a candidate neuron may have already been registered into another registered neuron *J*″, we compare the spatial rate map correlation of this candidate neuron with its companions in both registered neurons. If the spatial correlation between 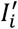 and the members of the targeted registered neuron is higher than the members of *J*″, we manually decide to re-match this neuron based on rule-based suggestions from our criteria.

It’s crucial to clarify that all candidates are derived from CellReg’s results without the introduction of new neurons or raw candidates. The optimization process is applied solely to spatial maps from Maze A and B, whereas maps from the reversed maze paradigm (RMP) and hairpin maze paradigm (HMP) tasks—including MAf, MAb, HPf, and HPb—are not optimized. Our findings demonstrate consistency across all data, confirming that our optimization of CellReg’s outputs does not compromise the validity of our conclusions. Moreover, our re-match algorithms maintain the cross-day pair-wise spatial correlation between place cells (Figures S2G-I) and, in some cases, even show a slight albeit non-significant improvement.

We conducted RMP and HMP training between Stage 1 and 2 of the MNP training for mice #10209 and #10212. Given that RMP and HMP training may influence the spatial representation in Maze A, we separately tracked place cells detected in Stages 1 and 2 for these mice, each spanning 13 sessions. For mice #10224 and #10227, RMP and HMP training occurred after all MNP training was complete, allowing us to track place cells in Maze A collectively across all 26 sessions from both stages. Consequently, this methodology resulted in six long-term spatial maps of Maze A, derived from our cohort of four mice.

### Place field registration

In RMP and HMP, place fields were considered bidirectional if they overlapped by 60% or more. The related chance level is computed by shifting the backward spatial maps, thereby dissociating the one-to-one relationships between forward and backward maps across place cells. To analyze the evolution of individual place fields across multiple days, we implemented a multi-step field tracking approach: 1) Initially, place fields were independently extracted from each session’s place cells using loose criteria. 2) Preliminary tracking was then performed based on field overlap, grouping fields with at least 75% overlap. 3) Recognizing that mere overlap might not account for positional fluctuations that could lead to tracking errors, we calculated and normalized the summed event rates, which report the average activity of each tracked field, for preliminarily grouped fields. Areas exhibiting average rates exceeding 0.5 (half of the peak rate) were identified as consistently encoded positions. Fields sharing these consistently encoded positions were considered identical and consequently registered together. This tracking process ensures that each tracked place field is distinctly separated, facilitating precise analysis of individual field evolution.

Technically, this tracking process forms a field index map 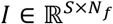, where *N*_*f*_ is the number of all registered field across *S* sessions, typically measured more than ten thousand. The entry at row *i* and column *j* of the field index map represents the state of field *j* at session *i*, where *j* is its unique index among all the fields detected across *S* sessions. The field state has three values: 0 for inactive, 1 for active, and NaN for cases where the neuron is either not detected or not successively registered in session *i*. We computed the field index map for place cells recorded in all the paradigms. We defined a place field’s lifespan as the maximum number of continuous sessions during which it remained active. Place fields that underwent recovery events were counted multiple times, treating each recovery as a new lifespan. However, this measurement is necessarily conservative due to potential issues such as non-detection or registration failures of cells across sessions. For instance, a place field observed for 10 sessions may still be active beyond this period, but without definitive evidence in subsequent sessions, its lifespan is conservatively reported as at least 10 sessions. Therefore, we only present the minimum estimated lifespans of place fields and provide cumulative counts of fields with lifespans exceeding specific thresholds, rather than exact durations.

### Analysis of the coordination of field evolution

To estimate the degree of coordination in the evolution of sibling fields, we defined a discrete random variable, the evolution events *E*, over our field index map. We considered field states across two continuous sessions, *t*_0_ to *t*_0_ + 1, with the following combinations:

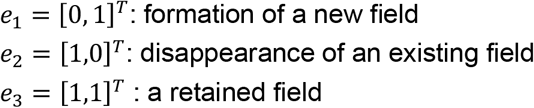

We excluded the case *e*_0_ = [0,0]^*T*^. The subscript of *e*_*i*_ is determined by converting the binary vector to a decimal representation, and these subscripts *i* ∈ {1,2,3} are referred to as ‘labels’ for each evolution event. This allocation of subscripts is for convenience when generalizing to cases with longer durations.

The evolution events *E* can be considered a property of a place field that reflects its evolution or fate over a future period. We hypothesize that the representations of sibling fields may drift coordinately, meaning that the evolution event *E* occurs dependently across sibling fields at a level significantly higher than it occurs across non-sibling fields. This implies that the fate of sibling place fields is more closely linked compared to fields from different neurons. We applied methods from probability theory and information theory to test such dependencies, as described in the “Analysis of Sibling Fields’ Properties Independence” section.

Additionally, we define the degree of coordination (CD) to quantitatively measure the extent of coordinated drift among sibling fields. Perfect coordination, where sibling fields evolve in a completely synchronized manner, corresponds to an expected joint probability matrix, 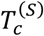, with diagonal entries defined by the marginal probabilities *P*_*X*_ (*x*_*i*_) and off-diagonal entries as zero:

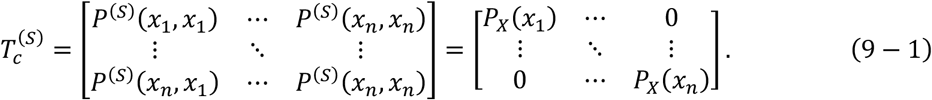

Conversely, the matrix corresponding to 0% coordination is *T*^(*S*)^ (formula 7-7). The CD is then defined as:

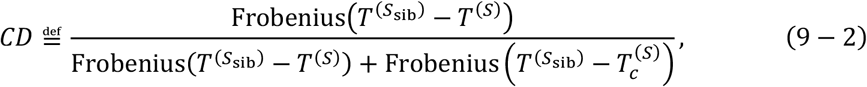

where the Frobenius norm is used to quantify the magnitude of residuals in these matrices. Intuitively, a value of CD close to zero indicates that the joint probability of evolution events among sibling fields are closer to 0% coordination than the 100% coordination, suggesting a minimal coordination among sibling fields.

For longer durations, we defined evolution events with continuous durations *d*, starting from session *t* and ending by session *t* + *d* − 1, the states of a certain place field have *2*^*d*^ − 1 situations (excluding the all-zero case):

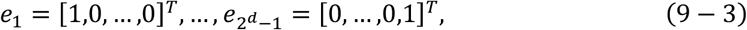

However, as the duration extends, the number of values increases exponentially, while the number of field samples declines. Therefore, if the duration is too long, the resulting contingency table will be too sparse, and the accuracy of the joint or marginal probability of evolution events *E* will decrease. Thus, we only considered durations *d* ∈ {2,3,4,5}, with continuous five sessions over eight days being considered a relatively long-term period. To compute the probability mass function of evolution events, *P*_*E*_(*e*) for 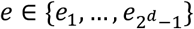, we used all state pairs that do not contain any NaN values during sessions *t*_0_ to *t*_0_ + *d* − 1.

To investigate which evolution events contribute to the observed coordination among sibling fields, we analyzed the residual probability matrices. This matrix is obtained by subtracting the expected joint probability 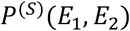 from the joint probability of either sibling field pairs 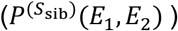 or non-sibling field pairs 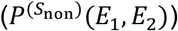, where *E*_1_ and *E*_2_ denote evolution events of a certain duration:

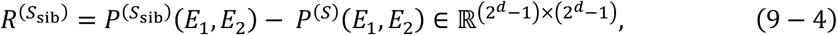

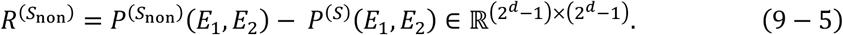

The entry 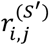 at row *i* and column *j* of residual matrix *R*^(*S′*^), where *S*^*′*^ ∈ {*S*_sib_, *S*_non_ }, represents the extent to which the probability of evolution events *e*_*i*_ and *e*_*j*_ occurring together exceeds the expected probability (*P* (*E* = *e*_*i*_)*P* (*E* = *e*_*j*_)). If the value 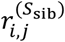 is significantly more than 0 and 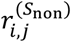, it indicates a statistically significant tendency for the co-appearance of *e*_*i*_ and *e*_*j*_. Conversely, if the value 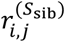 is significantly less than 0 and 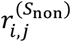, it suggests a statistically significant tendency for the mutual exclusivity of *e*_*i*_ and *e*_*j*_.

### Calculation of A-to-A and I-to-A conditional probability

We define the retained duration *T*_*a*_ as a continuous active period of a given field from the first session it becomes active (*t*_0_) to a determined session (*t*), with *T*_*a*_ = *t* − *t*_0_ + 1. It is important to note that *t* is not the session in which the given field transitions to the inactive state. The conditional probability of retained duration for *T*_*a*_ is 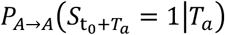, where 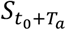 is the field state in the next session after *T*_*a*_ continuous active sessions. This probability is measured using fields continuously detected *T*_*a*_ sessions and is active for the first *T*_*a*_ sessions, with the fraction of active in session *T*_*a*_ + 1 representing the conditional probability for *T*_*a*_.

Similarly, the inactive duration *T*_*i*_ is the continuous inactive period from a session detecting an active-to-inactive transition (*t*_0_) to a determined session (*t*), with *T*_*i*_ = *t* − *t*_0_ + 1. We exclude continuous inactive periods starting from the first training session. The conditional probability of field recovery for *T*_*i*_ is 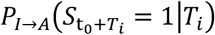, where 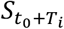 is the field state in the next session after *T*_*i*_ continuous inactive sessions. This probability is calculated similarly to *P*_*A*→*A*_.

The chance level of *P*_*I*→*A*_ was calculated by relocating field positions before field registration. In this relocation shuffle test, the index map and field sizes remain unchanged, but the field centers are randomly redistributed over the correct tracks of both mazes and the hairpin maze, as well as everywhere in the open field. This chance level reflects the minimal probability that two randomly distributed fields may overlap and thus appear to recover. Our goal was to determine whether recurrent events are due to such random occurrences. The computation of *P*_*I*→*A*_ is not necessary because it requires fields to be retained for many continuous sessions, and the likelihood that such continuous retention is caused by random events is extremely low.

### Kinetic models

#### Fitting *P*_*A*→*A*_ and *P*_*I*→*A*_

To model the dynamics of multi-field representations, we developed two kinetic models: the equal-rate drift model (EDM) and the convergent drift model (CDM). In EDM, the drift occurs at a constant rate, with place fields drifting at a static probability regardless of their active duration. In CDM, the drift rate reduces and converges to a low level 1 − *c*_∞_.

For EDMs, the conditional probability *P*_*A*→*A*_ is assumed to be constant (1 − *r*), independent of the retained duration *T*_*a*_, with *r* explored in the range {0.5, 0.6, 0.7, 0.8, 0.9}:

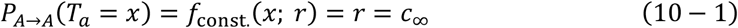

In CDMs, *P*_*A*→*A*_ is approximated by functions that monotonically increase and gradually converge to *c* _∞_ ∈ (0, 1]. We selected three candidate functions (‘exp.’, ‘reci.’, and ‘log.’) to fit our empirical data.

The ‘exp.’ function is defined as:

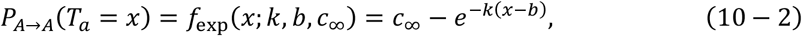

with parameters *k* and *b* estimated from experimental data:

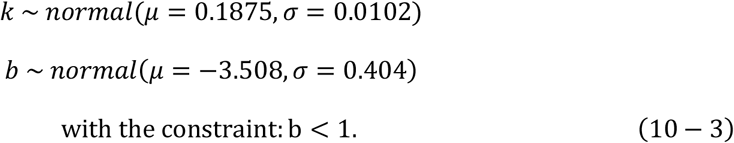

The ‘reci.’ function is defined as:

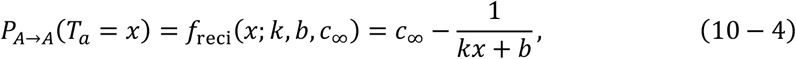

with parameters *k, b, c*_∞_ are measured from our data:

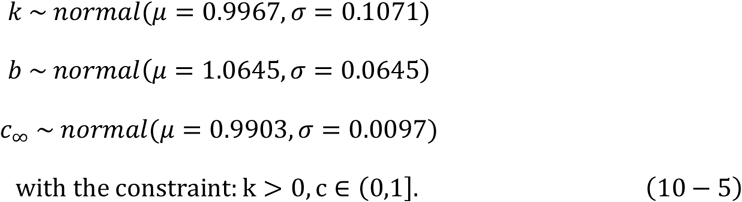

The ‘log.’ function is defined as:

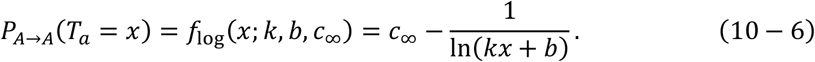

The initial value of all *c*_∞_ is set as 0.8. We used the ‘curve_fit’ function from SciPy to fit our data with these convergent functions. The metrics for assessing the goodness of fit (GOF) include the coefficient of determination (*r*^2^):

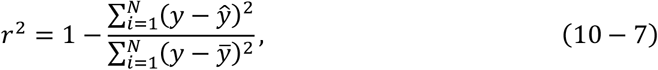

where *y* represents the real values of the conditional probability, 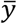 is the mean of all real values, *y* is the predicted value corresponding to the real values, and *N* is the number of values. Additionally, mean squared error (MSE) is also applied as a metric for goodness of fit.

Henceforth, we will use *f*(*x*; *k, b*) to denote all the three candidate functions.

Both models account for recurrent place fields. The probability *P*_*I*→*A*_ was approximated as a Kohlrausch-Williams-Watts function, a stretched exponential decay function with a lower convergence rate:

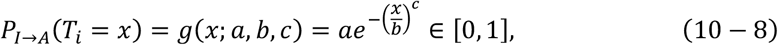

with parameters *a, b, c* estimated from real data:

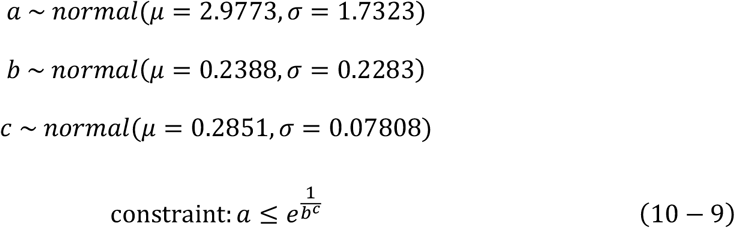

This function ensures that the recovery probability does not exceed a certain threshold, thereby preventing over-stabilization of the dynamic system. We also assumed that newly recovered fields behave similarly to newly formed fields and thus follow a similar transition probability *P*_*A*→*A*_.

#### Model equations

For each session *t*, let *A*_*t*_ denote the number of newly formed place fields, including recovered fields that that were inactive before session *t*, and *S*_*t*_ denote the number of place fields that survive from previous *t* − 1 sessions. The recursive expressions of *A*_*t*_ and *S*_*t*_ are:

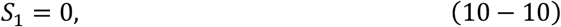

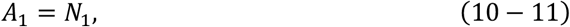

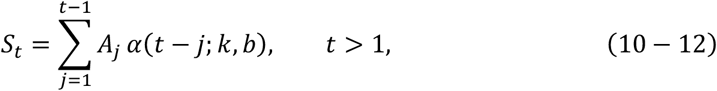

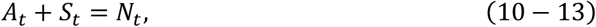

where *N*_*t*_ is the total number of place fields in session *t*, and *α*(*x*; *k, b*) is the cumulative productivity of *f*(*x*; *k, b*), given by

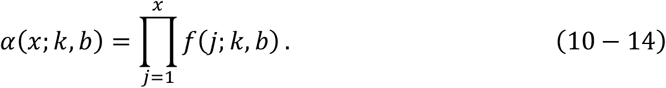

In Equal-Rate Drift Models (EDMs),

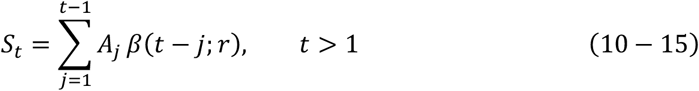

where *β*(*x*; *r*) is the cumulative productivity of the drift rate constant *r* and thus it is an exponential function:

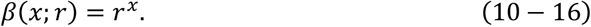

In the following context, we use *h*(*x*) to denote both *f*(*x*; *k, b*) and *β*(*x*; *r*), with *h*(0) = 1. Additionally, for simplicity, we assume that the total number of fields does not change over time, although we observed a significant decline:

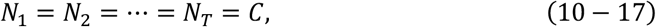

where *T* is the total number of sessions. This assumption simplifies the derivation of our models.

### Additional evidence for kinetic models

To assess whether the kinetic model based on single field evolution rules (SFER) is sufficient to account for representational dynamics at the ensemble level, we investigated two indices: the fraction of super stable place fields and survival fields.

#### 1. Fraction of super stable place fields (*SupS*(*t*; *θ*))

This index measures the fraction of super stable place fields among all active fields in session *t*. A super stable field is defined as one that has retained for a duration equal to or longer than a threshold *θ* up to session *t*. The fraction of super stable fields is thus a function of *θ* and *t*. The active fields at session *t* are defined as those fields that have been continuously detected, whether inactive or active, from session *t* − *θ* + 1 to *t*, and are active in session *t*. Fields that are not detected or registered in one or more sessions within the range from session *t* − *θ* + 1 to *t*, even if they are active in session *t*, are excluded from this definition.

The expression of *SupS*(*t*; *θ*) derived from our models is:

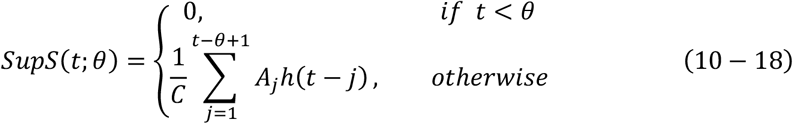

This function is discrete, but we defined its gradient as:

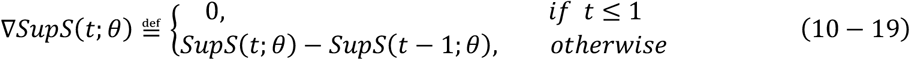

For both models and any *θ* ∈ ℕ^+^ and *θ* > 1, *SupS*(*t*; *θ*) will display a drastic elevation from zero at point *t* = *θ*, which we referred to as a breakpoint. The gradient at this breakpoint is a feature of *SupS*(*t*; *θ*) for both models:

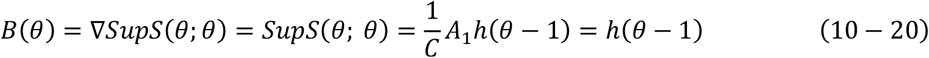

In EDMs, it is an exponential function:

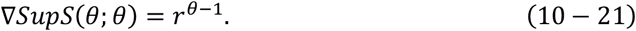

In CDMs,

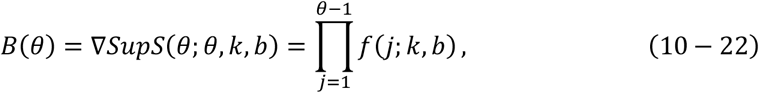

Equation (10 − 22) represents the SFER-predicted function (also the CDM-predicted one) for the fraction of super stable place fields, and we will demonstrate that it more accurately represents the increasing fraction of super stable fields for any given threshold θ than the EDM-predicted function.

#### 2. Fraction of survival place fields (*Surv*(Δ*t*; *t*_0_))

This index demonstrates the fraction of place fields that survived from session *t*_0_ to *t*. The survived fields are defined as those fields that remain continuously active from session *t*_0_ to *t*, Additionally, fields that have been continuously detected and registered, regardless of their active or inactive state, during the same period from session *t*_0_ to *t* are used as the denominator for calculating the survival fraction. The function of the survival index can be derived mathematically:

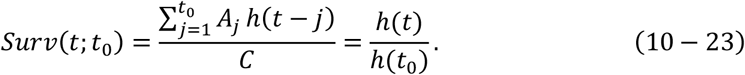

In EDMs, *Surv*(Δ*t*; *t*_0_) can be simplified as:

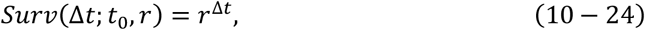

which is an exponential decay with respect to the session interval Δ*t* = *t* − *t*_0_. In CDMs,

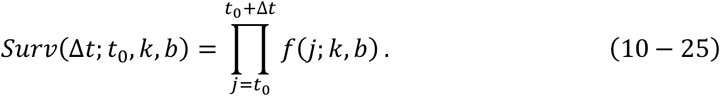

A key distinction between EDMs and CDMs is whether their dynamics display cumulative effects. The cumulative effect is a tendency for the survival fraction to gradually rise with respect to an increasing *t*_0_ for a given and fixed session interval Δ*t*, the value of *Surv*(*t*; *t*_0_) gradually rises with respect to an increasing *t*_0_, due to a cumulative of survived place fields. EDMs do not display the cumulative effect because the values of *Surv*(*t*; *t*_0_) do not change with *t*_0_:

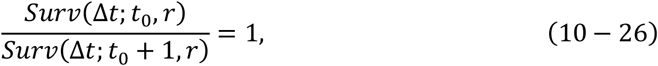

while in CDMs,

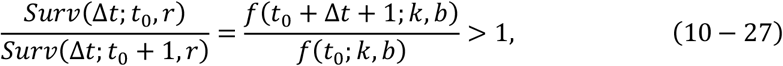

indicating a gradual increase as the candidate functions *f*(*x*; *k, b*) monotonically increase with respect to an increasing *x*. Equation (10 − 27) represents the SFER-predicted function for the cumulative effects observed in real data, and we will demonstrate that it more accurately represents the increasing trend of the fraction of super stable fields for any fixed session interval Δ*t* and any initial session *t*_0_ than the EDM-predicted function.

### Statistical tests

When comparing two sets of data that follow a normal distribution and have no significant difference in their variances, a two-sample t-test is applied. If a significant difference in variances is reported by Levene test, the Welch’s t-test is used. Both tests can be performed using the ‘scipy.stats.ttest_ind’ function. If the two samples have the same length and a one-to-one correspondence, a paired t-test is performed using the ‘scipy.stats.ttest_rel’ function. To test whether the mean of a group of data points is significantly greater or less than a specific value, a one-sample t-test is performed using the scipy.stats.ttest_1samp function. To compare whether two groups of data have the same distribution, a two-sample Kolmogorov–Smirnov test is performed using the ‘scipy.stats.ks_2samp’ function. To determine whether a group of data follows a certain distribution, Lilliefors-corrected KS test is performed. If not explicitly mentioned T-test, Welch’s test or Paired t-test used in the results and legends indicate two-sided test. If not explicitly mentioned, KS test used in the results and legends indicate two-sample KS test.

## Notes

### Competing Interest Statement

The authors have declared no competing interest.

