## Supplemental Figures for "Single field evolution rule governs the dynamics of representational drift in mouse hippocampal dorsal CA1 region"

**The PDF file includes:**

Figs. S1 to S18

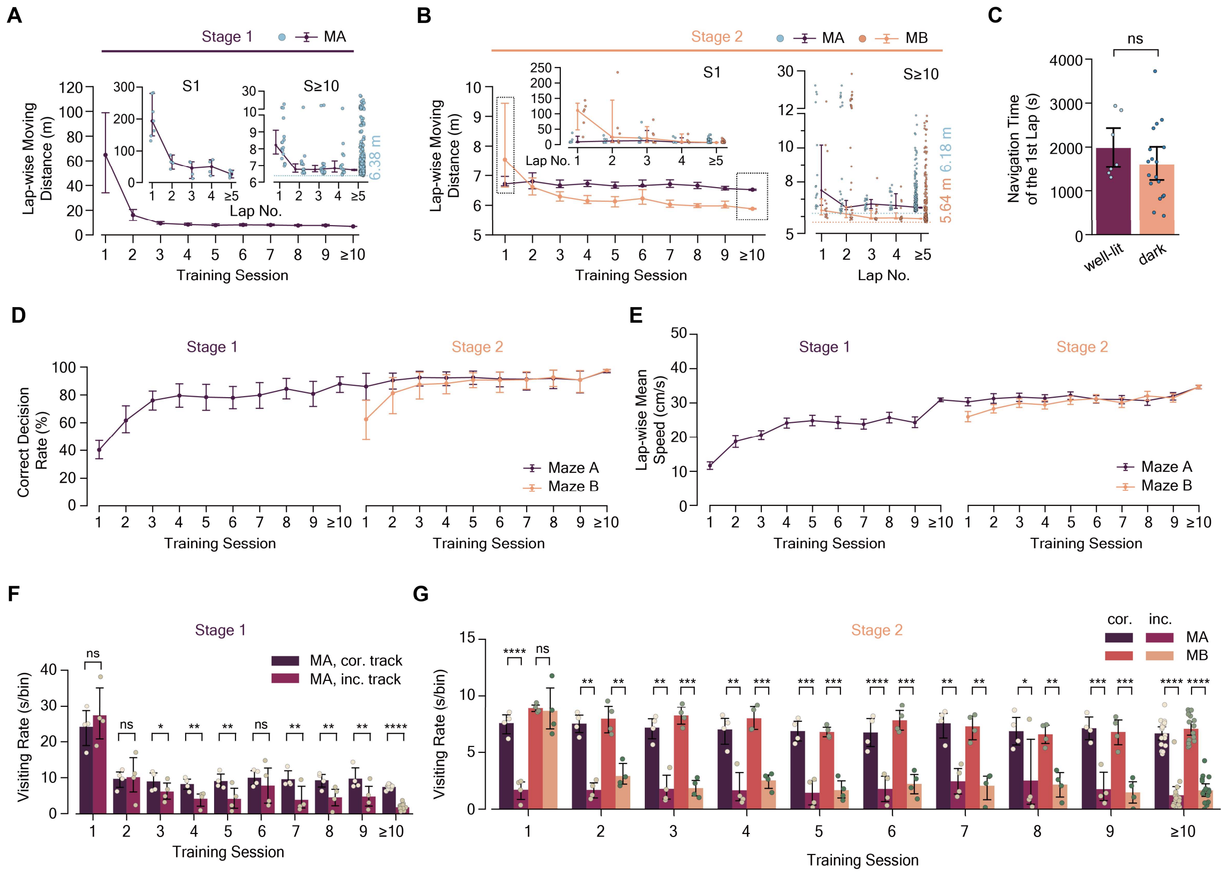


**Figure S1. Robust maze learning process demonstrated by multiple behavioral indexes.**

(**A** and **B**) Lap-wise moving distance (in meters) consistently declined throughout maze learning in Maze A (purple lines and blue dots) and B (orange lines and dots) for both Stage 1 (A) and 2 (B). Estimator: median. Insets show lap-wise moving distances for Stage 1 Session 1 (S1, left) and Session ≥10 (S≥10, right), on a per-lap basis, with blue dashed lines indicating the shortest recorded distances in Maze A (Stage 1: 6.38m; Stage 2: 6.18m ), and the orange dotted line marking the record in Maze B (Stage 2: 5.64m ). Data points represent individual laps, with laps ≥5 grouped together for clarity. Welch's two-sample t-test results: Maze A Stage 1 S1 (mean ± std: 90.6 ± 80.7m; median: 64.9m ; range: 12.5m to 282.5m ) vs. Stage 2 S≥10 (6.98 ± 1.79m ; median: 6.52m ; range: 6.18m to 23.01m ), P = $4.9\times10^{-5}$; Maze B Stage 2 S1 (21.74 ± 38.08m; median: 7.54m; range: 5.82m to 234.71m ) vs. Stage 2 S≥10 (6.52 ± 2.02m ; median: 5.88m ; range: 5.64m to 28.57m ), P = 0.0011.

(**C**) Initial lap time to locate the Maze A exit in well-lit (1979.8 ± 645.5s, range: 1304.0 to 2936.8s) and dark settings (1604.1 ± 819.8s, range: 433.8 to 3725.0s), showing no significant difference (two-sided two-sample t-test, P = 0.85). Individual data points represent mice.

(**D**) Correct decision rate in Maze A and Maze B during Stage 1 and 2 training, showing significant improvements (Welch's t-test): Maze A Stage 1 S1 (40.3 ± 9.3%; median: 36.8%) vs. Stage 2 S ≥ 10 (97.2 ± 2.5%; median: 98.0%), P=$3.7\times10^{-6}$; Maze B Stage 2 S1 (62.5 ± 18.0%; median: 65.0%) vs. Stage 2 S≥10 (97.5 ± 1.5%; median: 98.0%), P=0.007.

(**E**) Lap-wise mean speed (cm/s) increased significantly during maze learning: Welch's t-test, Maze 1 Stage 1 S1 (11.7 ± 2.8 cm/s) to Stage 2 S≥10 (34.7 ± 6.5 cm/s), P=$1.0\times10^{-25}$; Two-sides two-sample t test, Maze 2 Stage 2 S1 (26.0 ± 6.8 cm/s) to Stage 2 S≥10 (34.7 ± 6.8 cm/s), $P = 3.9\times10^{-23}$.

(**F** and **G**) Visiting rate for each session in Stage 1 (F) and Stage 2 (G), expressed as the average time mice spent per bin on the correct and incorrect tracks (n = 4 mice, Paired t-test). The visiting rates on each type of bin gradually diverged, consistent with the increasing correct decision rate. MA: Maze A; MB: Maze B; cor.: correct track; inc.: incorrect track. Error bars represent 95% confidence intervals. Significance levels: ns, P ≥ 0.05; *, P < 0.05; **, P < 0.01; ***, P < 0.001; ****, P < 0.0001.

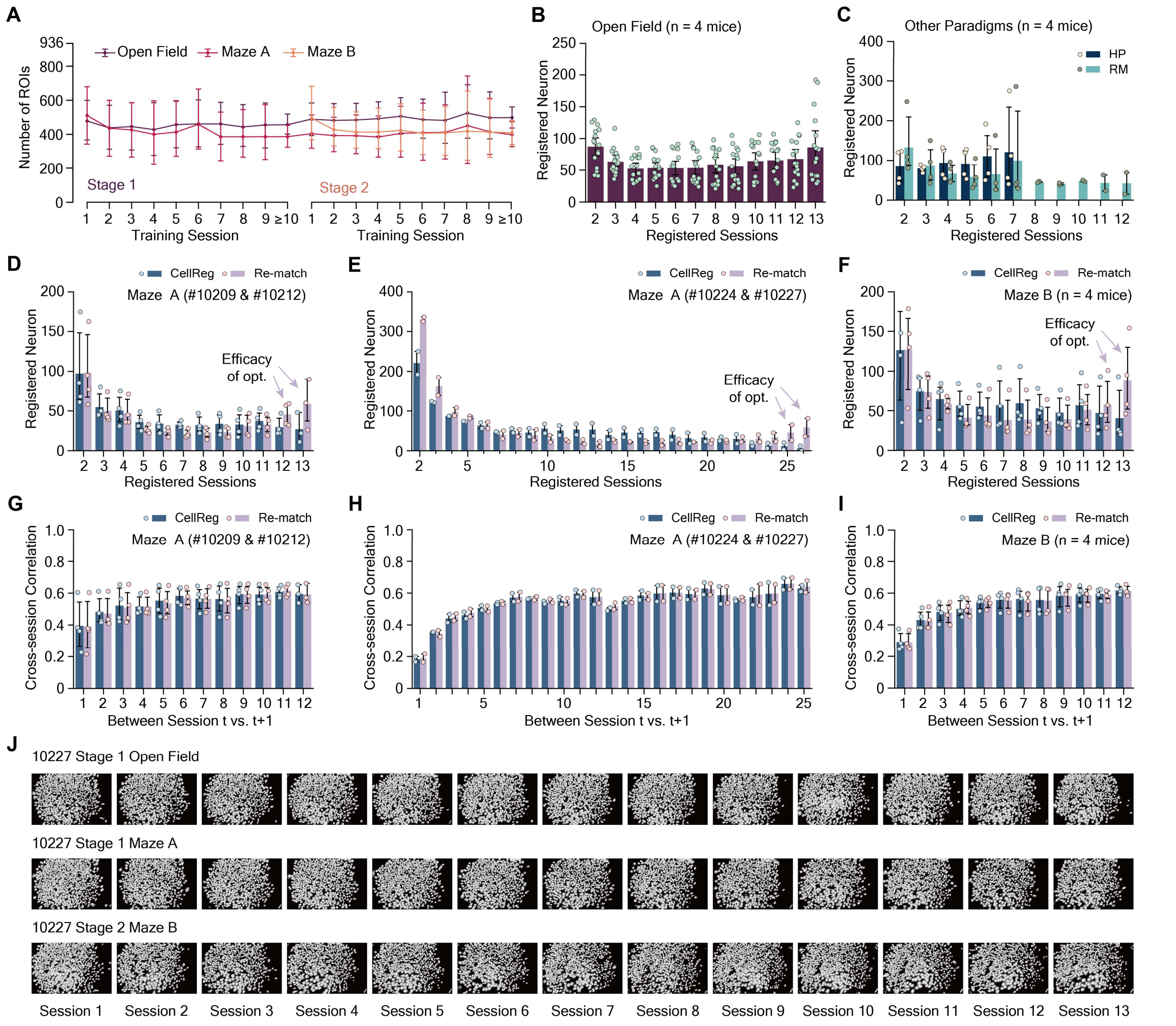


**Figure S2. Tracking and optimization of neuronal registration across multiple days in different behavioral paradigms and environments.**

(**A**) The number of regions of interest (ROIs) detected in the field of views during the maze navigation paradigm (n = 363 sessions from 4 mice). The range of detected ROIs varied from 174 to 936 (Maze A: 174–867; Maze B: 202–764; Open Field: 203–936).

(**B**) Tracking neurons in open field sessions using CellReg. Each mouse underwent two open field sessions for 13 blocks in either Stage 1 or 2, resulting in n = 16 tracked open field spatial maps from 4 mice. The 'Registered session' indicates the number of sessions in which a neuron was registered. Missing positions may be due to registration failure, non-detection, neural silence, etc.

(**C**) Tracking neurons in reversed maze paradigm (RM) and hairpin maze paradigm (HP) using CellReg. Mice #10209 and #10212 underwent RM training for 12 sessions over 15 days, while mice #10224 and #10227 underwent HP training for 7 sessions over 7 days. All 4 mice underwent HP training for 7 sessions over 7 days.

(**D** to **F**) Comparisons between the cell tracking results of CellReg and our re-match optimization (see

Methods) for Maze A sessions (D to E) and Maze B sessions (F). Briefly, the re-match optimization aims to match as many neurons together as possible while maintaining reliability, providing an adequate sample size for precise quantitative measurements. Light purple arrows highlight the increased number of neurons registered over 12 sessions in Maze A (D, mice #10209 and #10212, n = 4 tracked spatial maps, CellReg: 199; Re-match: 358, Paired t-test, P = 0.009) and Maze B (F, n = 4 tracked spatial maps for 4 mice, CellReg: 350; Re-match: 583, Paired t-test, P = 0.02). The optimization showed robust results when tracking neurons across 26 sessions in Maze A (E, n = 2 mice (#10224 and #10227), CellReg: 93 neurons over 23 sessions with only 12 neurons registered in all 26 sessions; Re-match: 338 neurons over 23 sessions with 117 neurons registered in all 26 sessions).

(**G** to **I**) The application of our re-match algorithm on spatial maps of either Maze A or Maze B sessions did not significantly affect the cross-session correlation of spatial rate maps between matched neurons in adjacent sessions, suggesting that the optimization did not introduce more false-positive results than CellReg itself. Paired t-test: Maze A, P > 0.2 for all cross-session comparisons with session interval = 1 (degree of freedom (df) = 5, n = 4 mice); Maze B: P > 0.13 for all cross-session comparisons with session interval = 1 (df = 3, n = 4 mice). (**J**) Example field of views of mouse #10227 in different environments across 13 sessions over approximately 26 days. Width × height: 700 × 450 µm.

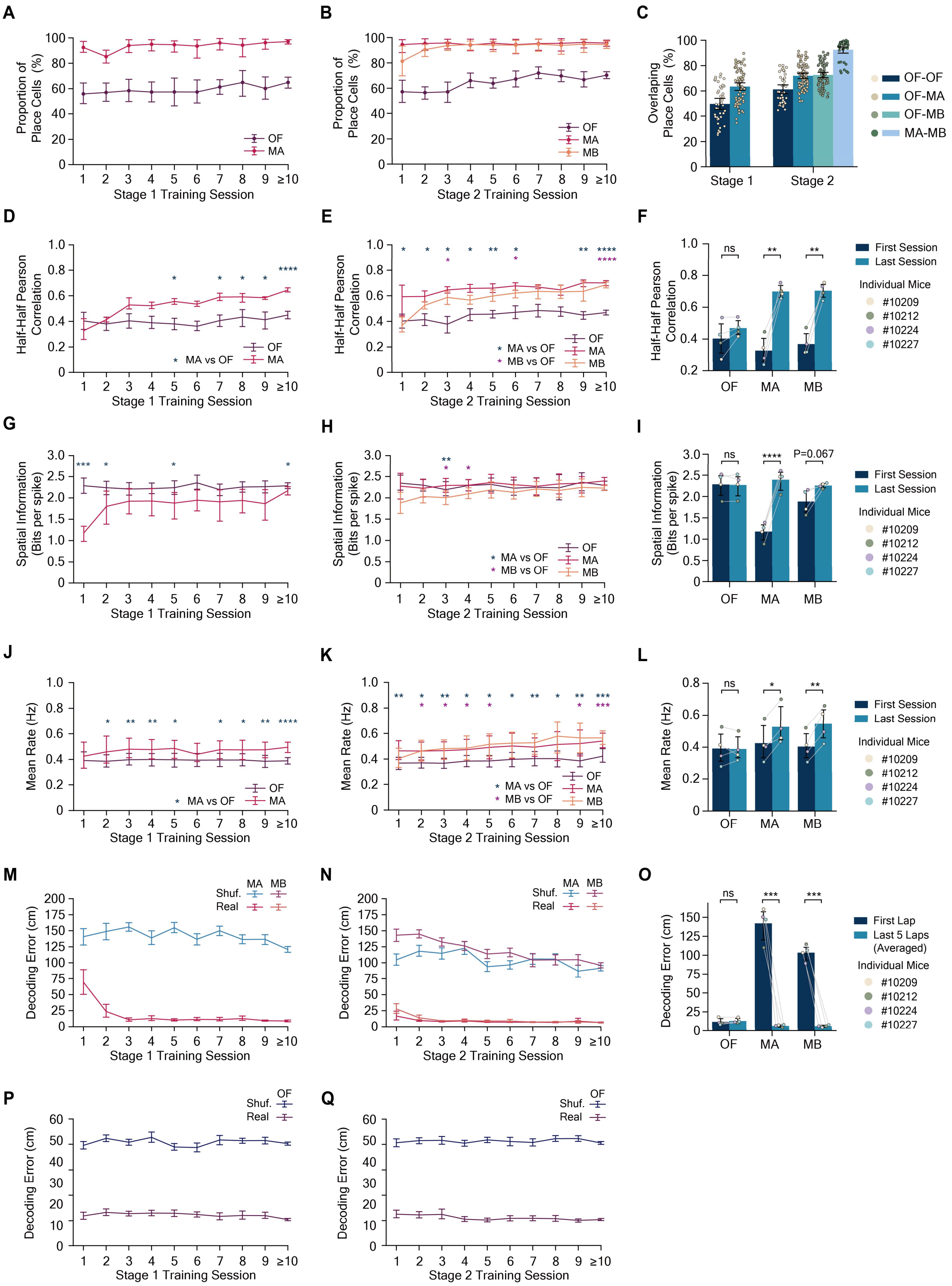
A
Proportion of Place Cells (%)
OF
MA
Stage 1 Training Session
B
Proportion of Place Cells (%)
OF
MA
MB
Stage 2 Training Session
C
Overlaping Place Cells (%)
Stage 1
Stage 2
OF-OF
OF-MA
OF-MB
MA-MB
D
Half-Half Pearson Correlation
* MA vs OF
OF
MA
Stage 1 Training Session
E
Half-Half Pearson Correlation
* MA vs OF
* MB vs OF
OF
MA
MB
Stage 2 Training Session
F
Half-Half Pearson Correlation
ns
First Session
Last Session
Individual Mice
#10209
#10212
#10224
#10227
OF
MA
MB
G
Spatial Information (Bits per spike)
Stage 1 Training Session
H
Spatial Information (Bits per spike)
* MA vs OF
* MB vs OF
OF
MA
MB
Stage 2 Training Session
I
Spatial Information (Bits per spike)
ns
P=0.067
First Session
Last Session
Individual Mice
#10209
#10212
#10224
#10227
OF
MA
MB
J
Mean Rate (Hz)
* MA vs OF
OF
MA
Stage 1 Training Session
K
Mean Rate (Hz)
* MA vs OF
* MB vs OF
OF
MA
MB
Stage 2 Training Session
L
Mean Rate (Hz)
ns
First Session
Last Session
Individual Mice
#10209
#10212
#10224
#10227
OF
MA
MB
M
Decoding Error (cm)
MA MB
Shuf.
Real
Stage 1 Training Session
N
Decoding Error (cm)
MA MB
Shuf.
Real
Stage 2 Training Session
O
Decoding Error (cm)
ns
First Lap
Last 5 Laps (Averaged)
Individual Mice
#10209
#10212
#10224
#10227
OF
MA
MB
P
Decoding Error (cm)
OF
Shuf.
Real
Stage 1 Training Session
Q
Decoding Error (cm)
OF
Shuf.
Real
Stage 2 Training Session

**Figure S3. Within-session stability and coding accuracy of hippocampal spatial representations enhanced throughout maze learning process.**

(**A** and **B**) Proportion of place cells in open field (OF), Maze A (MA), and Maze B (MB) during Stage 1 (A) and Stage 2 (B). Place cells' proportion in mazes consistently higher than in open field (n=4 mice, Paired t-test, df=3 for S1 to S9 and df=15 for S≥10, P<0.032 for all comparisons between open field and maze sessions). No significant difference between Maze A and B (Paired t-test, P>0.09).

(**C**) Percentage of active neurons identified as place cells in two different sessions: OF-OF (49.8±14.2% in Stage 1, 61.2±11.0% in Stage 2), OF-MA (63.5±13.2% in Stage 1, 72.0±9.9% in Stage 2), OF-MB (72.7±9.6%), MA-MB (92.5±8.0%).

(**D** to **F)** Within-session stability of place cells, measured as the half-half session correlation, significantly increased during Stage 1 (D) and 2 (E) in complex mazes (F, n = 4 mice, Paired t-test: MA, 0.33 ± 0.08 (Stage 1 S1) to 0.70 ± 0.04 (Stage 2 S13), P = 0.004; MB, 0.37 ± 0.06 (S1) to 0.70 ± 0.04 (S13), P = 0.005), while remaining unchanged in the open field (F, Paired t-test: 0.40 ± 0.09 (Stage 1 S1) to 0.47 ± 0.05 (Stage 2 S13), P = 0.11). Eventually, the half-half correlation in complex mazes became significantly higher than that in the open field (Paired t-test with Bonferroni correction, P < 1 × 10-8 for both mazes).

(**G** to **I**) Spatial information (unit: bits per spike) of place cells significantly increased during Stage 1 (G) and 2 (H) in complex mazes (**I**, n = 4 mice, Paired t-test: MA, 1.18 ± 0.19 (Stage 1 S1) to 2.39 ± 0.21 (Stage 2 S13), P = $3 \times 10^{-5}$; MB, 1.88 ± 0.25 (S1) to 2.26 ± 0.05 (S13), P = 0.067), while remaining unchanged in the open field (**I**, Paired t-test: 2.29 ± 0.25 (Stage 1 S1) to 2.27 ± 0.23 (Stage 2 S13), P = 0.75). Eventually, spatial information in both mazes showed no significant difference compared to that in the open field (Paired t-test with Bonferroni correction for both comparisons: P > 0.13).

(**J** to **L**) Mean event rate (unit: Hz) of place cells displayed a mild but significant increase during Stage 1 (J) and Stage 2 (K) in complex mazes (L, n = 4 mice, df = 3, Paired t-test: MA, 0.42 ± 0.10 (Stage 1 S1) to 0.53 ± 0.10 (Stage 2 S13), P = 0.012; MB, 0.40 ± 0.08 (S1) to 0.55 ± 0.09 (S13), P = 0.004), while remaining unchanged in the open field (Paired t-test: 0.39 ± 0.09 (Stage 1 S1) to 0.39 ± 0.07 (Stage 2 S13), P = 0.92). The mean event rate in complex mazes eventually became significantly higher than that in the open field (Paired t-test with Bonferroni correction, P < 0.05 for both mazes).

(**M** to **Q**) We employed a modified Naïve Bayesian Classifier (Methods) to decode mice's positions in two mazes (M and N) and open fields (P and Q) based on neural activities. The decoding errors (unit: cm) showed a significant decline in complex mazes (O, n = 4 mice, df = 3, Paired t-test: MA, 142.1 ± 19.2 cm (first lap in Stage 1 S1) to 6.6 ± 1.5 cm (mean of the last 5 laps in Stage 2 S13), P = 0.0009; MB, 103.4 ± 9.2 cm (first lap in S1) to 6.2 ± 1.3 cm (mean of the last 5 laps in S13), P = 0.0003) but remained unchanged in the open field (Paired t-test: 11.8 ± 3.7 cm (first 1/5 of S1) to 13.0 ± 2.9 cm (average of the 5 1/5 pieces of Stage 2 S13), P = 0.32). Shuf.: Decoding results based on shuffled data. Real: Decoding results based on actual data. Error bars represent 95% confidence intervals. Significance levels: ns, P ≥ 0.05; *, P < 0.05; **, P < 0.01; ***, P < 0.001; ****, P < 0.0001. For figures (A to L), data from all within-lap neural activities of place cells were used, while for the decoding analysis (M to Q), only neural activities occurring when mice were moving forward on the correct track were considered. All open field sessions were included for visualization, but for statistical analysis, the two open field sessions per block were averaged for paired t-tests. Bonferroni correction was applied for multiple comparisons across open field (OP), Maze A (MA), and Maze B (MB). Colored dots in (C, F, I, L, O) represent individual mice, and gray lines connect data points from the first and last sessions for each mouse.

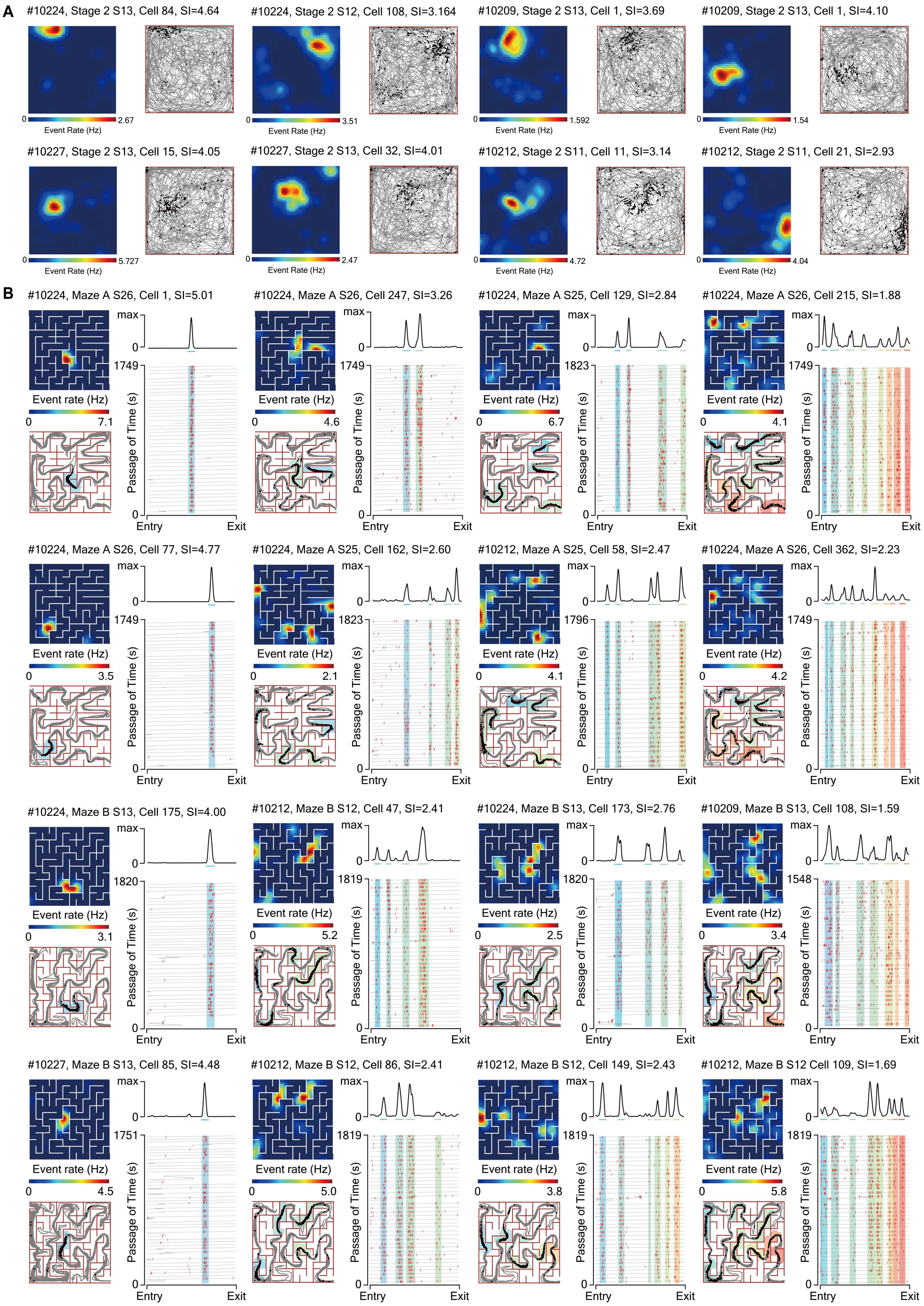
A
#10224, Stage 2 S13, Cell 84, SI=4.64
#10224, Stage 2 S12, Cell 108, SI=3.164
#10209, Stage 2 S13, Cell 1, SI=3.69
#10209, Stage 2 S13, Cell 1, SI=4.10
#10227, Stage 2 S13, Cell 15, SI=4.05
#10227, Stage 2 S13, Cell 32, SI=4.01
#10212, Stage 2 S11, Cell 11, SI=3.14
#10212, Stage 2 S11, Cell 21, SI=2.93
Event Rate (Hz)
B
#10224, Maze A S26, Cell 1, SI=5.01
#10224, Maze A S26, Cell 247, SI=3.26
#10224, Maze A S25, Cell 129, SI=2.84
#10224, Maze A S26, Cell 215, SI=1.88
#10224, Maze A S26, Cell 77, SI=4.77
#10224, Maze A S25, Cell 162, SI=2.60
#10212, Maze A S25, Cell 58, SI=2.47
#10224, Maze A S26, Cell 362, SI=2.23
#10224, Maze B S13, Cell 175, SI=4.00
#10212, Maze B S12, Cell 47, SI=2.41
#10224, Maze B S13, Cell 173, SI=2.76
#10209, Maze B S13, Cell 108, SI=1.59
#10227, Maze B S13, Cell 85, SI=4.48
#10212, Maze B S12, Cell 86, SI=2.41
#10212, Maze B S12, Cell 149, SI=2.43
#10212, Maze B S12 Cell 109, SI=1.69
Event rate (Hz)
Passage of Time (s)
Entry
Exit
max

**Figure S4. Example place cells in open field and complex mazes**

**(A and B)** Representative place cells with either single or multiple fields recorded in the open field (A) and both mazes (B). Each example cell is labeled with the animal ID, recording session, neuron index, and spatial information (SI).

(**A**) For visualizing place cells in the open field, two subfigures are used: the event rate map (left) and the spatially organized events-trajectory map (right). The event rate map with Gaussian smoothing shows the event rate of the neuron, while the spatially organized events-trajectory map displays the mouse's trajectory (gray lines) and the spatial distribution of calcium events (black dots).

(**B**) For visualizing place cells in complex mazes, four subfigures arranged in a 2 × 2 grid layout are used: the event rate map (top left), the spatially organized events-trajectory map (bottom left), the linearized event rate map (top right), and the temporally organized events-trajectory map (bottom right). The events rate maps and the spatially organized events-trajectory map are similar to those for the open field. The linearized event rate map visualizes the event rate along the correct track, highlighting the peak(s) of activity. The temporally organized events-trajectory map shows the temporal distribution of calcium events along the correct track, providing evidence of reliable fields repetitively responding to positions. Red bars indicate calcium events, while gray lines represent linearized mouse trajectory in correct path. Colored background areas in the spatially organized events-trajectory map, colored bands in the temporally organized events-trajectory map, and colored bars below the linearized event rate map mark the ranges of individual place fields. The same color is used for the same field. For all visualized place cells in complex mazes, only neural activities occurring when mice were moving forward on the correct track were considered.

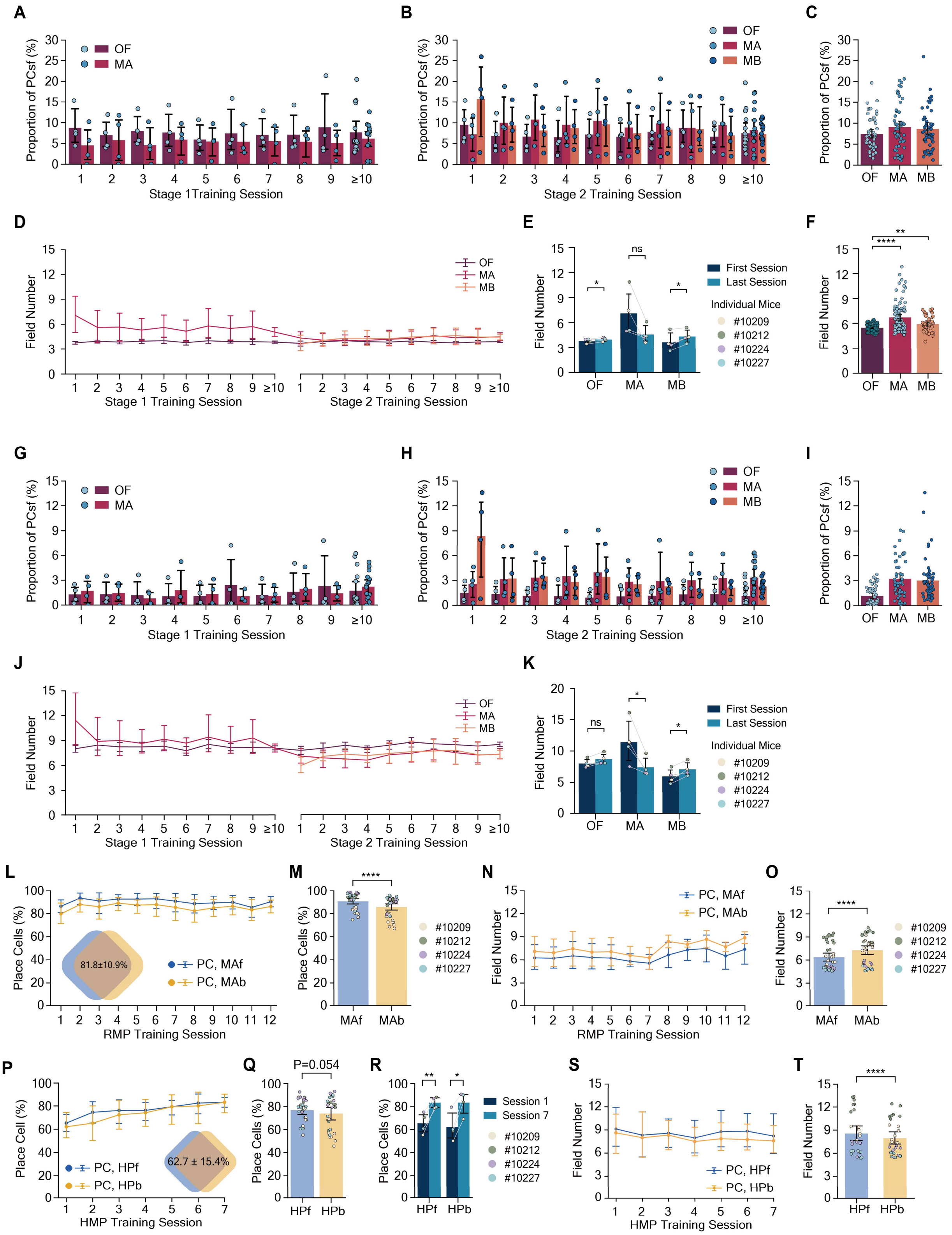

A
Proportion of PCsf (%)
OF
MA
Stage 1Training Session
B
OF
MA
MB
Stage 2 Training Session
C
D
Field Number
Stage 1 Training Session
Stage 2 Training Session
E
First Session
Last Session
Individual Mice
#10209
#10212
#10224
#10227
ns
F
G
H
I
J
K
L
Place Cells (%)
81.8±10.9%
PC, MAf
PC, MAb
RMP Training Session
M
MAf
MAb
N
O
P
Place Cell (%)
PC, HPf
PC, HPb
62.7 ± 15.4%
HMP Training Session
Q
P=0.054
HPf
HPb
R
Session 1
Session 7
S
T

**Figure S5. Most place cells exhibit multiple place fields regardless of criteria for field identification and behavioral paradigms.**

To identify place fields, we employed two sets of criteria: rigorous criteria (**A** to **F**) and loose criteria (**G** to **K**) (see Methods).

(**A** to **C)** Under the rigorous criteria, the proportion of place cells with a single field (PCsf) in the open field (7.6 ± 4.8% in Stage 1 and 7.4 ± 3.9% in Stage 2), Maze A (5.6 ± 3.7% in Stage 1, 9.1 ± 5.6% in Stage 2), and Maze B (8.6 ± 4.9%), remained low during Stage 1 (A) and Stage 2 (B).

(**D** to **F**) The average number of fields per place cell exhibited a mild but not significant reduction in Maze A (E, Paired t-test, n = 4 mice: 7.1 ± 2.4 in Stage 1 S1 to 4.6 ± 0.9 in Stage 2 S13, P = 0.096) and a mild but significant increase in Maze B (Paired t-test, n = 4 mice: 3.6 ± 1.0 in S1 to 4.3 ± 0.7 in S13, P = 0.022). The open field also displayed a subtle but significant increase (Paired t-test, n = 4 mice: 3.8 ± 0.2 in Stage 1 S1 to 3.9 ± 0.2 in Stage 2 S13, P = 0.027). (F) Despite these fluctuations, the number of fields in both Maze A and B remains significantly higher than in the open field (Paired t-test with Bonferroni Correction: MA (4.9 ± 1.5) vs. OF (3.9 ± 0.3), df = 99 from 4 mice, P = $1 \times 10^{-9}$; MB (4.4 ± 0.9) vs. OF (3.9 ± 0.3), df = 51 from 4 mice, P = 0.001).

(**G** to **I)** Under the loose criteria, the proportion of PCsf in the open field was lower (1.6 ± 2.0% in Stage 1, 1.2 ± 1.1% in Stage 2), and similar trends were observed in Maze A (1.7 ± 1.5% in Stage 1, 3.2 ± 2.3% in Stage 2) and Maze B (3.0 ± 2.5%) during Stage 1 (G) and Stage 2 (H).

(**J** and **K**) The average number of fields per place cell under the loose criteria showed a significant reduction in Maze A (K, Paired t-test, n = 4 mice: 11.4 ± 3.1 in Stage 1 S1 to 7.4 ± 1.3 in Stage 2 S13, P = 0.040) and a mild but significant increase in Maze B (Paired t-test, n = 4 mice: 6.0 ± 1.0 in S1 to 7.1 ± 1.0 in S13, P = 0.024). The open field exhibited a subtle but not significant increase (Paired t-test, n = 4 mice: 8.0 ± 0.6 in Stage 1 S1 to 8.7 ± 0.8 in Stage 2 S13, P = 0.061). These results demonstrate that most place cells under either the rigorous or loose criteria display undeniable multiple place fields.

(**L** to **O**) display results obtained in reversed maze paradigm. (L) The proportion of place cells encoding both directions, with overlapped colored squares indicating that 81.8 ± 10.9% of place cells encode both directional movements. (M) The proportion of place cells in backward directional movement (85.7 ± 8.5%) is mildly but significantly lower than that of forward movement (Paired t-test, n = 4 mice, df = 37: P = $7 \times 10^{-12}$), possibly due to distinct levels of familiarization. (N) Place cells encoding both directions display multiple place fields at the ensemble level, (O) with the average number of fields for backward direction (7.3 ± 1.8) mildly but significantly greater than that for forward direction (6.4 ± 1.7, Paired t-test, n = 4 mice, df = 37: P = $4.4 \times 10^{-8}$). This suggests that the total number of fields to encode both directions remains relatively constant. Calcium events detected on incorrect paths and during backward movement were excluded. Four mice were trained for session 1 to 7, with two of them receiving additional training (session 8 to 12).

(**P** to **T**) display results obtained in hairpin maze paradigm. (P) The proportion of place cells encoding both directions, with overlapped colored squares indicating that 62.7 ± 15.4% of place cells encode both directional movements. (Q) The proportion of place cells encoding both the forward (76.7 ± 9.9%) and backward (73.8 ± 14.8%) movements showed no significant difference (Paired t-test, n = 4 mice, df = 27: P = 0.054). (R) The proportion of place cells encoding both directions gradually increased over time (Paired t-test, n = 4 mice, forward: from 65.3 ± 7.8% (S1) to 83.2 ± 4.2% (S7), P = 0.005; backward: from 62.1 ± 10.6% (S1) to 83.2 ± 8.7% (S7), P = 0.013). (S to T) Place cells in the hairpin maze displayed evident multi-field coding, with the average number of fields for the forward direction (8.6 ± 2.6) being mildly but significantly greater than that for the backward direction (8.0 ± 2.2, Paired t-test, n = 4 mice, df = 27: P = $2.3 \times 10^{-5}$). Error bars represent 95% confidence intervals. Significance levels:

ns, $P \geq 0.05$; *, $P < 0.05$; **, $P < 0.01$; ****, $P < 0.0001$. All open field sessions were included for visualization, but for statistical analysis, the two open field sessions per block were averaged for paired t-tests.

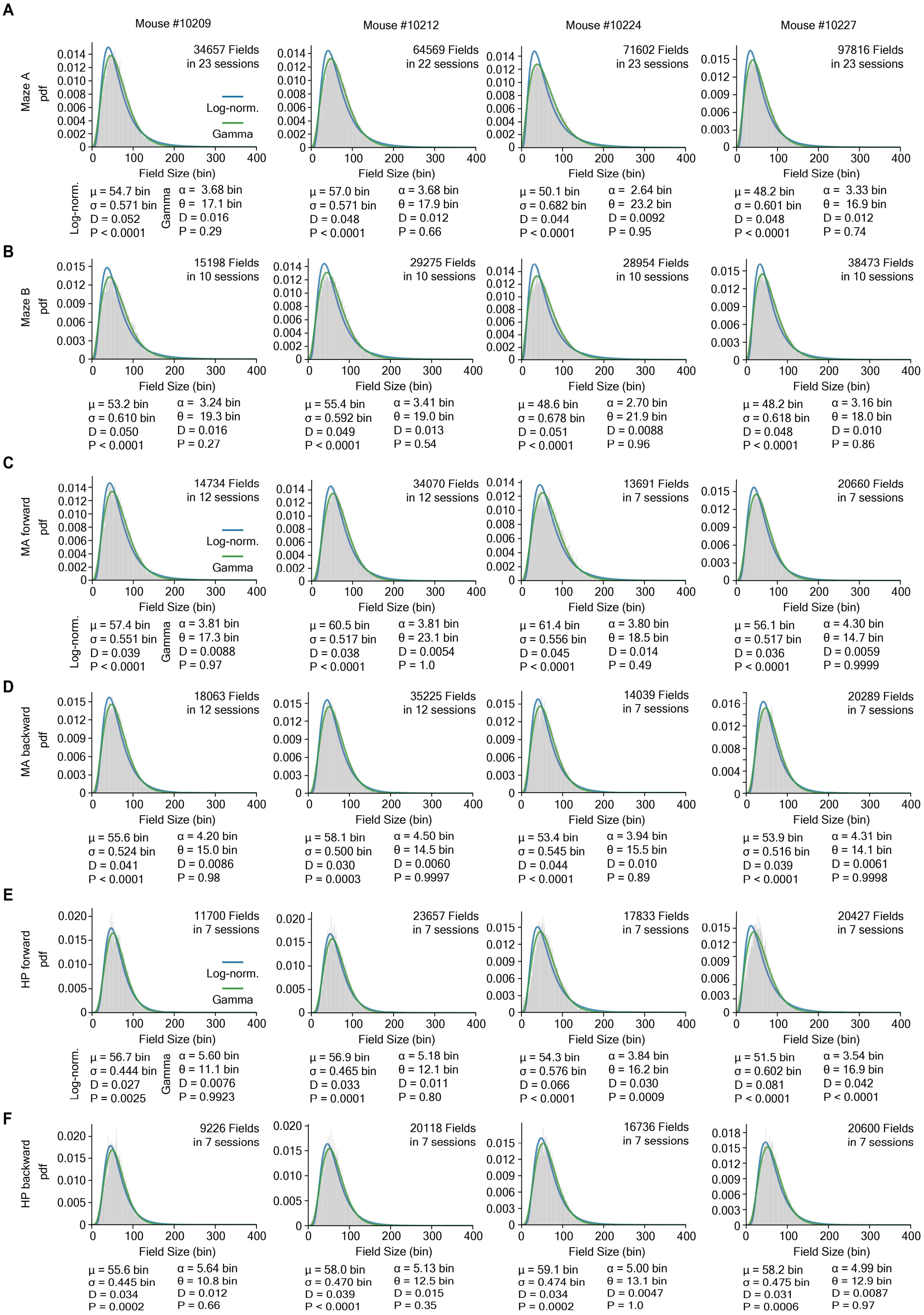
A
Mouse #10209
Mouse #10212
Mouse #10224
Mouse #10227
Maze A
pdf
34657 Fields in 23 sessions
64569 Fields in 22 sessions
71602 Fields in 23 sessions
97816 Fields in 23 sessions
Log-norm.
Gamma
Field Size (bin)
μ = 54.7 bin σ = 0.571 bin D = 0.052 P < 0.0001
α = 3.68 bin θ = 17.1 bin D = 0.016 P = 0.29
μ = 57.0 bin σ = 0.571 bin D = 0.048 P < 0.0001
α = 3.68 bin θ = 17.9 bin D = 0.012 P = 0.66
μ = 50.1 bin σ = 0.682 bin D = 0.044 P < 0.0001
α = 2.64 bin θ = 23.2 bin D = 0.0092 P = 0.95
μ = 48.2 bin σ = 0.601 bin D = 0.048 P < 0.0001
α = 3.33 bin θ = 16.9 bin D = 0.012 P = 0.74
B
Maze B
15198 Fields in 10 sessions
29275 Fields in 10 sessions
28954 Fields in 10 sessions
38473 Fields in 10 sessions
μ = 53.2 bin σ = 0.610 bin D = 0.050 P < 0.0001
α = 3.24 bin θ = 19.3 bin D = 0.016 P = 0.27
μ = 55.4 bin σ = 0.592 bin D = 0.049 P < 0.0001
α = 3.41 bin θ = 19.0 bin D = 0.013 P = 0.54
μ = 48.6 bin σ = 0.678 bin D = 0.051 P < 0.0001
α = 2.70 bin θ = 21.9 bin D = 0.0088 P = 0.96
μ = 48.2 bin σ = 0.618 bin D = 0.048 P < 0.0001
α = 3.16 bin θ = 18.0 bin D = 0.010 P = 0.86
C
MA forward
14734 Fields in 12 sessions
34070 Fields in 12 sessions
13691 Fields in 7 sessions
20660 Fields in 7 sessions
μ = 57.4 bin σ = 0.551 bin D = 0.039 P < 0.0001
α = 3.81 bin θ = 17.3 bin D = 0.0088 P = 0.97
μ = 60.5 bin σ = 0.517 bin D = 0.038 P < 0.0001
α = 3.81 bin θ = 23.1 bin D = 0.0054 P = 1.0
μ = 61.4 bin σ = 0.556 bin D = 0.045 P < 0.0001
α = 3.80 bin θ = 18.5 bin D = 0.014 P = 0.49
μ = 56.1 bin σ = 0.517 bin D = 0.036 P < 0.0001
α = 4.30 bin θ = 14.7 bin D = 0.0059 P = 0.9999
D
MA backward
18063 Fields in 12 sessions
35225 Fields in 12 sessions
14039 Fields in 7 sessions
20289 Fields in 7 sessions
μ = 55.6 bin σ = 0.524 bin D = 0.041 P < 0.0001
α = 4.20 bin θ = 15.0 bin D = 0.0086 P = 0.98
μ = 58.1 bin σ = 0.500 bin D = 0.030 P = 0.0003
α = 4.50 bin θ = 14.5 bin D = 0.0060 P = 0.9997
μ = 53.4 bin σ = 0.545 bin D = 0.044 P < 0.0001
α = 3.94 bin θ = 15.5 bin D = 0.010 P = 0.89
μ = 53.9 bin σ = 0.516 bin D = 0.039 P < 0.0001
α = 4.31 bin θ = 14.1 bin D = 0.0061 P = 0.9998
E
HP forward
11700 Fields in 7 sessions
23657 Fields in 7 sessions
17833 Fields in 7 sessions
20427 Fields in 7 sessions
μ = 56.7 bin σ = 0.444 bin D = 0.027 P = 0.0025
α = 5.60 bin θ = 11.1 bin D = 0.0076 P = 0.9923
μ = 56.9 bin σ = 0.465 bin D = 0.033 P = 0.0001
α = 5.18 bin θ = 12.1 bin D = 0.011 P = 0.80
μ = 54.3 bin σ = 0.576 bin D = 0.066 P < 0.0001
α = 3.84 bin θ = 16.2 bin D = 0.030 P = 0.0009
μ = 51.5 bin σ = 0.602 bin D = 0.081 P < 0.0001
α = 3.54 bin θ = 16.9 bin D = 0.042 P < 0.0001
F
HP backward
9226 Fields in 7 sessions
20118 Fields in 7 sessions
16736 Fields in 7 sessions
20600 Fields in 7 sessions
μ = 55.6 bin σ = 0.445 bin D = 0.034 P = 0.0002
α = 5.64 bin θ = 10.8 bin D = 0.012 P = 0.66
μ = 58.0 bin σ = 0.470 bin D = 0.039 P < 0.0001
α = 5.13 bin θ = 12.5 bin D = 0.015 P = 0.35
μ = 59.1 bin σ = 0.474 bin D = 0.034 P = 0.0002
α = 5.00 bin θ = 13.1 bin D = 0.0047 P = 1.0
μ = 58.2 bin σ = 0.475 bin D = 0.031 P = 0.0006
α = 4.99 bin θ = 12.9 bin D = 0.0087 P = 0.97

**Figure S6. Field sizes of place fields in goal-directed navigation paradigm are well-described by Log-normal and Gamma distributions.**

(**A** to **F**) Field sizes (unit: bins, each bin is 4 $cm^2$) recorded in Maze A (A), Maze B (B), forward movement in reversed maze paradigm (C), backward movement in reversed maze paradigm (D), forward movement in hairpin maze paradigm (E), and backward movement in hairpin maze paradigm (F), were well-fitted by skewed distributions (e.g., gamma and gamma distributions). The goodness of fit for log-normal and gamma distributions was assessed by Lilliefors-corrected Kolmogorov-Smirnov tests (see Methods). The location parameters for both distributions were forcibly set to 0. Gamma distribution provides better estimation for most situations ($P > 0.27$ for 22 of 24 spatial maps, KS statistic $D = 0.012 \pm 0.008$), compared to Log-normal distribution ($P < 0.003$ for all 24 spatial maps, KS statistic $D = 0.044 \pm 0.012$, which is significantly greater than that of gamma distribution, Paired t-test $P = 4 \times 10^{-18}$). $\mu$ and $\sigma$ represent the mean and standard deviation of the log-normal distribution, respectively. $\alpha$ and $\theta$ indicate the shape and scale parameters of the gamma distribution, respectively. D: KS statistic; P: P-values determined by Monte Carlo simulation. Pdf: probability density function of field sizes. Gray bars: histogram of actual field size distribution. Blue and green lines indicate the pdfs of fitted log-normal and gamma distributions, respectively. MA: Maze A; HP: Hairpin maze. Novel sessions of maze navigation paradigm (Maze: Stage 1 S1 to 3; Maze B: Stage 2 S1 to 3) were excluded from analysis. Data from mouse #10209, shown in Fig. 2e, are also included here to correlate with the parameters and statistical results presented.

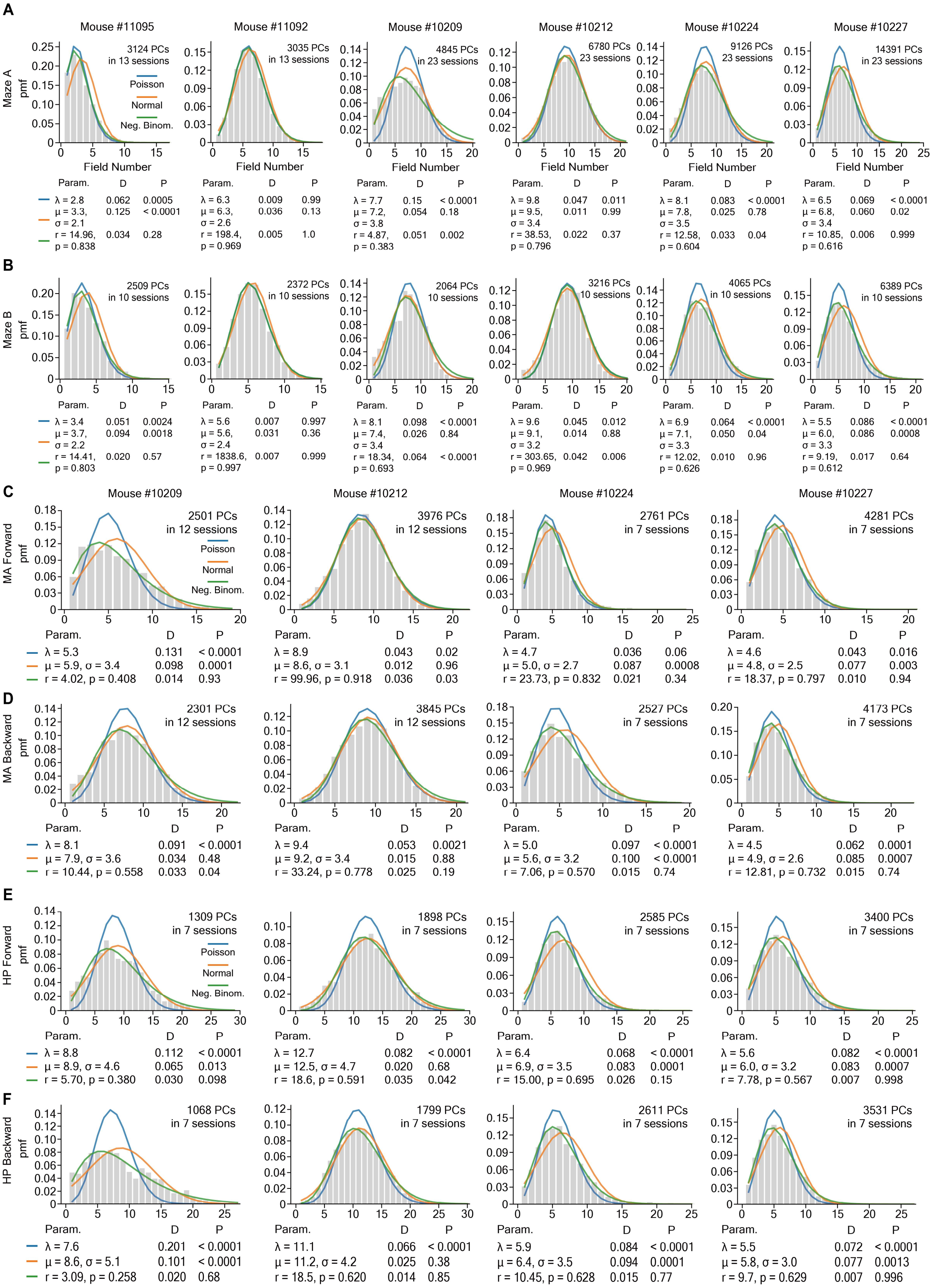
A
Mouse #11095
Mouse #11092
Mouse #10209
Mouse #10212
Mouse #10224
Mouse #10227
Maze A
pmf
Field Number
3124 PCs in 13 sessions
3035 PCs in 13 sessions
4845 PCs in 23 sessions
6780 PCs 23 sessions
9126 PCs 23 sessions
14391 PCs in 23 sessions
Poisson
Normal
Neg. Binom.
Param. D P
λ = 2.8 0.062 0.0005
μ = 3.3, σ = 2.1 0.125 < 0.0001
r = 14.96, p = 0.838 0.034 0.28
λ = 6.3 0.009 0.99
μ = 6.3, σ = 2.6 0.036 0.13
r = 198.4, p = 0.969 0.005 1.0
λ = 7.7 0.15 < 0.0001
μ = 7.2, σ = 3.8 0.054 0.18
r = 4.87, p = 0.383 0.051 0.002
λ = 9.8 0.047 0.011
μ = 9.5, σ = 3.4 0.011 0.99
r = 38.53, p = 0.796 0.022 0.37
λ = 8.1 0.083 < 0.0001
μ = 7.8, σ = 3.5 0.025 0.78
r = 12.58, p = 0.604 0.033 0.04
λ = 6.5 0.069 < 0.0001
μ = 6.8, σ = 3.4 0.060 0.02
r = 10.85, p = 0.616 0.006 0.999
B
Maze B
2509 PCs in 10 sessions
2372 PCs in 10 sessions
2064 PCs 10 sessions
3216 PCs 10 sessions
4065 PCs in 10 sessions
6389 PCs in 10 sessions
Param. D P
λ = 3.4 0.051 0.0024
μ = 3.7, σ = 2.2 0.094 0.0018
r = 14.41, p = 0.803 0.020 0.57
λ = 5.6 0.007 0.997
μ = 5.6, σ = 2.4 0.031 0.36
r = 1838.6, p = 0.997 0.007 0.999
λ = 8.1 0.098 < 0.0001
μ = 7.4, σ = 3.4 0.026 0.84
r = 18.34, p = 0.693 0.064 < 0.0001
λ = 9.6 0.045 0.012
μ = 9.1, σ = 3.2 0.014 0.88
r = 303.65, p = 0.969 0.042 0.006
λ = 6.9 0.064 < 0.0001
μ = 7.1, σ = 3.3 0.050 0.04
r = 12.02, p = 0.626 0.010 0.96
λ = 5.5 0.086 < 0.0001
μ = 6.0, σ = 3.3 0.086 0.0008
r = 9.19, p = 0.612 0.017 0.64
C
Mouse #10209
Mouse #10212
Mouse #10224
Mouse #10227
MA Forward
2501 PCs in 12 sessions
3976 PCs in 12 sessions
2761 PCs in 7 sessions
4281 PCs in 7 sessions
Param. D P
λ = 5.3 0.131 < 0.0001
μ = 5.9, σ = 3.4 0.098 0.0001
r = 4.02, p = 0.408 0.014 0.93
λ = 8.9 0.043 0.02
μ = 8.6, σ = 3.1 0.012 0.96
r = 99.96, p = 0.918 0.036 0.03
λ = 4.7 0.036 0.06
μ = 5.0, σ = 2.7 0.087 0.0008
r = 23.73, p = 0.832 0.021 0.34
λ = 4.6 0.043 0.016
μ = 4.8, σ = 2.5 0.077 0.003
r = 18.37, p = 0.797 0.010 0.94
D
MA Backward
2301 PCs in 12 sessions
3845 PCs in 12 sessions
2527 PCs in 7 sessions
4173 PCs in 7 sessions
Param. D P
λ = 8.1 0.091 < 0.0001
μ = 7.9, σ = 3.6 0.034 0.48
r = 10.44, p = 0.558 0.033 0.04
λ = 9.4 0.053 0.0021
μ = 9.2, σ = 3.4 0.015 0.88
r = 33.24, p = 0.778 0.025 0.19
λ = 5.0 0.097 < 0.0001
μ = 5.6, σ = 3.2 0.100 < 0.0001
r = 7.06, p = 0.570 0.015 0.74
λ = 4.5 0.062 0.0001
μ = 4.9, σ = 2.6 0.085 0.0007
r = 12.81, p = 0.732 0.015 0.74
E
HP Forward
1309 PCs in 7 sessions
1898 PCs in 7 sessions
2585 PCs in 7 sessions
3400 PCs in 7 sessions
Param. D P
λ = 8.8 0.112 < 0.0001
μ = 8.9, σ = 4.6 0.065 0.013
r = 5.70, p = 0.380 0.030 0.098
λ = 12.7 0.082 < 0.0001
μ = 12.5, σ = 4.7 0.020 0.68
r = 18.6, p = 0.591 0.035 0.042
λ = 6.4 0.068 < 0.0001
μ = 6.9, σ = 3.5 0.083 0.0001
r = 15.00, p = 0.695 0.026 0.15
λ = 5.6 0.082 < 0.0001
μ = 6.0, σ = 3.2 0.083 0.0007
r = 7.78, p = 0.567 0.007 0.998
F
HP Backward
1068 PCs in 7 sessions
1799 PCs in 7 sessions
2611 PCs in 7 sessions
3531 PCs in 7 sessions
Param. D P
λ = 7.6 0.201 < 0.0001
μ = 8.6, σ = 5.1 0.101 < 0.0001
r = 3.09, p = 0.258 0.020 0.68
λ = 11.1 0.066 < 0.0001
μ = 11.2, σ = 4.2 0.025 0.38
r = 18.5, p = 0.620 0.014 0.85
λ = 5.9 0.084 < 0.0001
μ = 6.4, σ = 3.5 0.094 0.0001
r = 10.45, p = 0.628 0.015 0.77
λ = 5.5 0.072 < 0.0001
μ = 5.8, σ = 3.0 0.077 0.0013
r = 9.7, p = 0.629 0.007 0.996

**Figure S7. Field number per place cell in goal-directed navigation paradigm are well-described by negative binomial distribution but with a Poisson-like shape.**

(**A** to **F**) The number of fields per place cell recorded in Maze A (A), Maze B (B), forward movement in the reversed maze paradigm (C), backward movement in the reversed maze paradigm (D), forward movement in the hairpin maze paradigm (E), and backward movement in the hairpin maze paradigm (F) were well-fitted by negative binomial and discrete normal distributions but poorly fitted by the Poisson distribution. The goodness of fit for each distribution was assessed by Lilliefors-corrected Kolmogorov-Smirnov tests (see Methods). The negative binomial distribution provides the best estimation for most situations ($P > 0.09$ for 19 of 26 spatial maps and $P > 0.03$ for 24 of 26 spatial maps, KS statistic $D = 0.023 \pm 0.014$), compared to the normal distribution ($P > 0.05$ for 12 of 26 spatial maps, KS statistic $D = 0.060 \pm 0.033$, which is significantly greater than that of the negative binomial distribution, Paired t-test with Bonferroni correction: $P = 1.2 \times 10\text{-}4$) and the Poisson distribution ($P > 0.05$ for 3 of 26 spatial maps, KS statistic $D = 0.075 \pm 0.039$, Paired t-test with Bonferroni correction: $P = 2.2 \times 10\text{-}7$). $\lambda$ is the rate parameter of the Poisson distribution; $\mu$ and $\sigma$ represent the mean and standard deviation of the normal distribution, respectively; r and p indicate the shape and scale parameters of the negative binomial distribution, respectively. D: KS statistic; P: P-values determined by Monte Carlo simulation. pmf: probability mass function of the field number per place cell. Gray bars: histogram of the actual field size distribution. Blue, orange, and green lines indicate the pmfs of fitted Poisson, normal, and negative binomial distributions, respectively. Param.: parameters; PC: place cells; MA: Maze A; HP: Hairpin maze. Novel sessions of the maze navigation paradigm (Maze A: Stage 1 S1 to S3; Maze B: Stage 2 S1 to S3) were excluded from analysis. Data from mouse #10227, shown in Fig. 2f, are also included here to correlate with the parameters and statistical results presented.

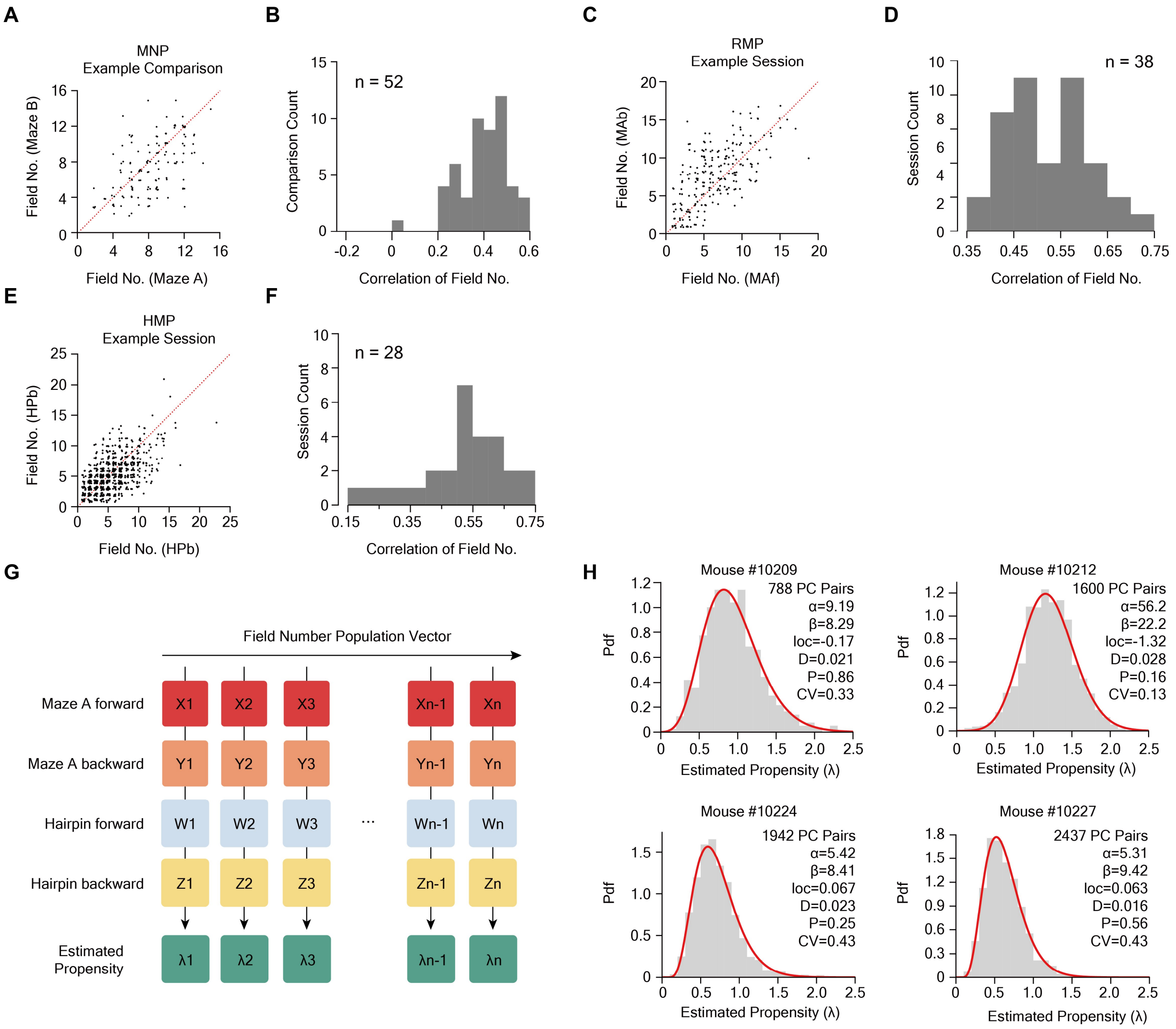


**Figure S8. Statistical analysis reveals limited neuronal propensity in hippocampal place cells for forming place fields.**

(**A** and **B**) We tracked the same neurons across Maze A and Maze B within each training block during Stage 2 of the maze navigation paradigm (MNP), obtaining 52 comparisons in total (4 mice, each trained for 13 blocks during Stage 2). An example comparison (mouse #10224, Block 7) with a Pearson correlation of 0.56 ($P = 2.2 \times 10^{-12}$) for field number is shown in (A), where each black dot represents a tracked neuron in both mazes, with added jitters for visualization due to integer field numbers. The histogram in (B) summarizes all 52 comparisons (mean ± std.: 0.39 ± 0.11; range: 0.00 to 0.58; $P < 0.05$ for n = 48 of 52 comparisons).

(**C** and **D**) Place cells encoding both forward and backward directions in each session of the reversed maze paradigm (RMP) training were used to compute Pearson correlations between field numbers for both directions. An example session (mouse #10209, RMP session 5) with a correlation of 0.65 ($P = 3.6 \times 10^{-28}$) is displayed in (C). The histogram in (D) shows the correlations for all 38 RMP sessions (mean ± std.: 0.53 ± 0.08; range: 0.38 to 0.73; $P < 3.6 \times 10^{-12}$ for all n = 38 sessions). (**E** and **F**) Similarly, place cells encoding both directions in each session of the hairpin maze paradigm (HMP) training were analyzed. An example session (mouse #10227, HMP session 4) with a correlation of 0.58 ($P = 2.5 \times 10^{-63}$) is shown in (E). The histogram in (F) summarizes all 28 HMP sessions (mean ± std.: 0.51 ± 0.14; range: 0.17 to 0.75; P < 0.0005 for all n = 28 sessions).

(**G** and **H**) The process for estimating neuronal propensity is schematically demonstrated (see Methods for details). By tracking neurons across Maze A and the hairpin maze, with each neuron having two distinct spatial maps for forward and backward movement in each environment, we obtain four field numbers $(X_i, Y_i, W_i, Z_i)$ for each neuron $i$. These values are used to calculate the estimated propensity $\lambda_i$ (unit: No. of place field per meter) for each neuron. The Gamma-Poisson model assumes that this propensity follows a gamma distribution, which was observed in all four mice (Kolmogorov-Smirnov test, P > 0.15 for all mice), as shown in (H). The fitted gamma distributions are indicated by red lines. The coefficient of variance (CV), which is the inverse of the square root of the shape parameter α, is used to assess the relative variance of the fitted gamma distribution. The lower the variance, the more limited the heterogeneity of neuronal propensity. The CV computed from a previously reported value (α = 0.571) is 1.32, while even the highest CV calculated from our data is more than three times lower, suggesting much more limited neuronal heterogeneity. Only place cells detected with at least one place field in each of the four spatial maps were included in the estimation.

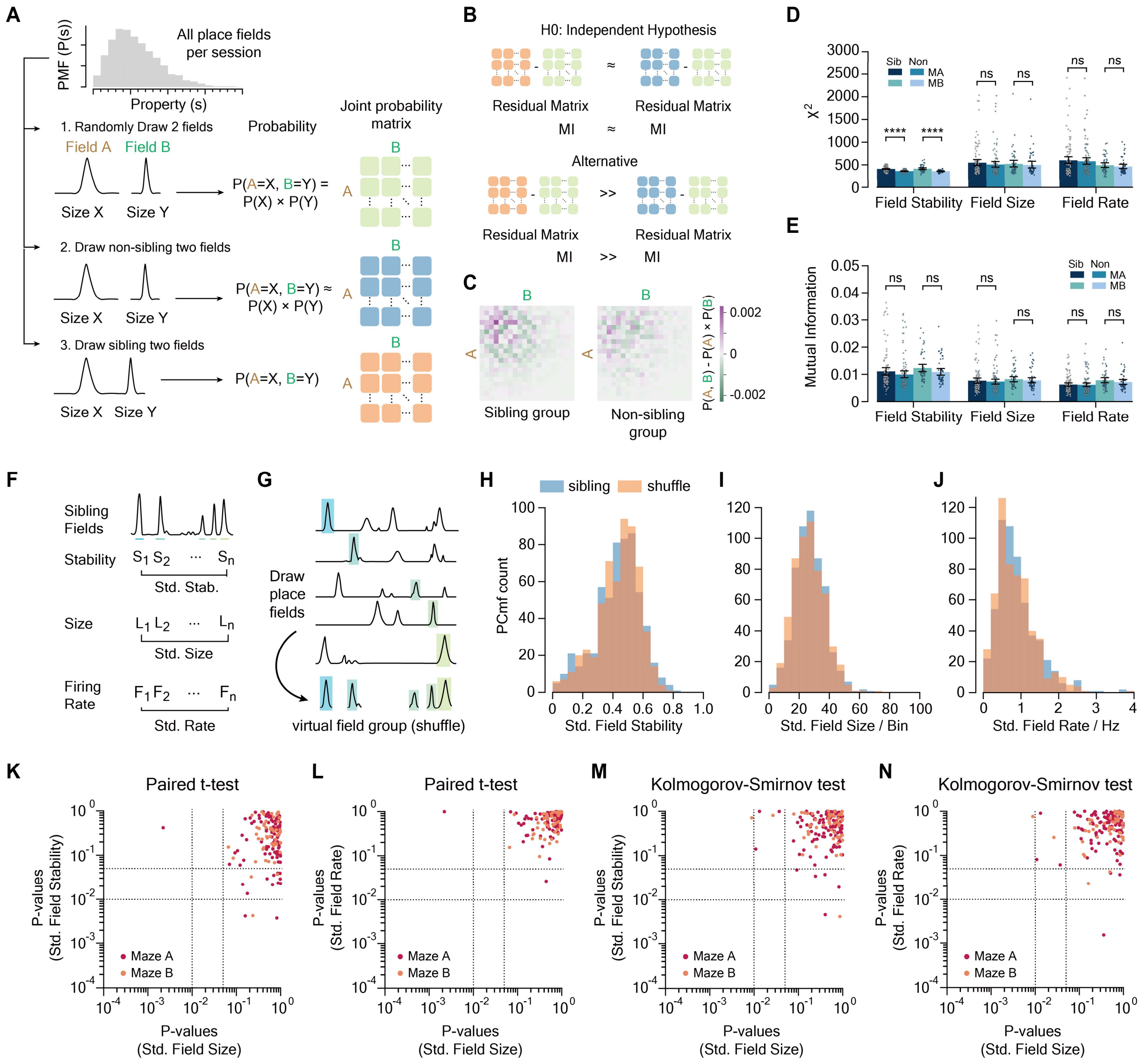


**Figure S9. Analysis reveals high independence of sibling place fields' properties in complex mazes.**

(**A**) Illustration of the probabilistic approach to assess the independence of sibling place fields' properties (see Methods). The top left panel displays an example probability mass function (PMF, denoted as P(s)) of place fields' properties per session. We calculated three joint probability matrices per session: the expected joint probability ($P^{(S)}$, green matrix), the joint probability of non-sibling fields ($P^{(S_{\mathrm{non}})}$, blue matrix), and that of sibling fields ($P^{(S_{\mathrm{sib}})}$, orange). Entries in these matrices represent the probability that two fields exhibit specific property values within predefined ranges of bins for each property. Field properties were binned into 20 bins, resulting in 20 × 20 matrices.

(**B**) The null hypothesis assumes that properties are independent among sibling fields, implying that $P^{(S_{\mathrm{sib}})}$ should not deviate significantly from $P^{(S)}$ beyond the deviation observed between $P^{(S_{\mathrm{non}})}$ and $P^{(S)}$. $\chi^2$ and mutual information (MI) are metrics to assess these deviations. Fields smaller than 16 bins were excluded from stability analysis.

(**C**) Example residual matrix calculated by subtracting either $P^{(S_{\text{sib}})}$ or $P^{(S_{\text{non}})}$ to $P^{(S)}$ (mouse #10227, Maze A, Stage 2 S13).

(**D** and **E**) both $\chi^2$ (D) and MI (E) metrics indicate high independence of field properties among sibling fields, with no significant differences between sibling (Sib) and non-sibling (Non) groups in most comparisons (two-sample two-sided t-test, $P > 0.3$ for field rate and sizes in both mazes), except a subtle relationship in their within-session stability ($\chi^2$, $P < 1 \times 10^{-6}$ for both mazes; MI, $P > 0.13$ for both mazes). Each point represents a session (n = 4 mice, 103 Maze A sessions, 52 Maze B sessions). MA: Maze A; MB: Maze B.

(**F** and **G**) Schematic of the third approach to assess sibling fields' property independence (see Methods). Only place cells with more than 2 fields were considered for this analysis. The null hypothesis posits that the standard deviation (std.) of the properties of a sibling field group (**F**) equals a chance level determined by a shuffle test (**G**), which randomly draws fields from different place cells to form a virtual group.

(**H** to **J**) Distribution of the std. of stability (H), field size (I), and field center rate (J) for an example session (mouse #10227, Maze A, Stage 2 S13). Blue bars represent real sibling fields' std. distribution, and orange bars represent virtual groups' std. distribution. No significant differences were observed in either mean values or empirical cumulative distributions in each comparison (Paired t-test for means: stability, P = 0.75; field size, P = 0.29; field rate, P = 0.22. Two-sample Kolmogorov-Smirnov test for distributions: stability, P = 0.79; field size, P = 0.33; field rate, P = 0.33).

(**K** to **N**) P-values from Paired t-tests (K and L) and two-sample KS tests (M and N) for all Maze A and B sessions during maze navigation paradigm training (n = 4 mice). For each property's std., most sessions showed no significant deviations from chance levels in either mean (Paired t-test: stability, P ≥ 0.05 for 91 of 103 Maze A sessions and 49 of 52 Maze B sessions; field size, P ≥ 0.05 for 101 of 103 Maze A sessions and 52 of 52 Maze B sessions; field rate, P ≥ 0.05 for 102 of 103 Maze A sessions and 52 of 52 Maze B sessions) or empirical cumulative distribution (two-sample KS test: stability, P ≥ 0.05 for 98 of 103 Maze A sessions and 51 of 52 Maze B sessions; field size, P ≥ 0.05 for 99 of 103 Maze A sessions and 50 of 52 Maze B sessions; field rate, P ≥ 0.05 for 100 of 103 Maze A sessions and 50 of 52 Maze B sessions). Dotted lines mark P = 0.05 and P = 0.01. Significance levels: ns, P ≥ 0.05; ****, P < 0.0001.

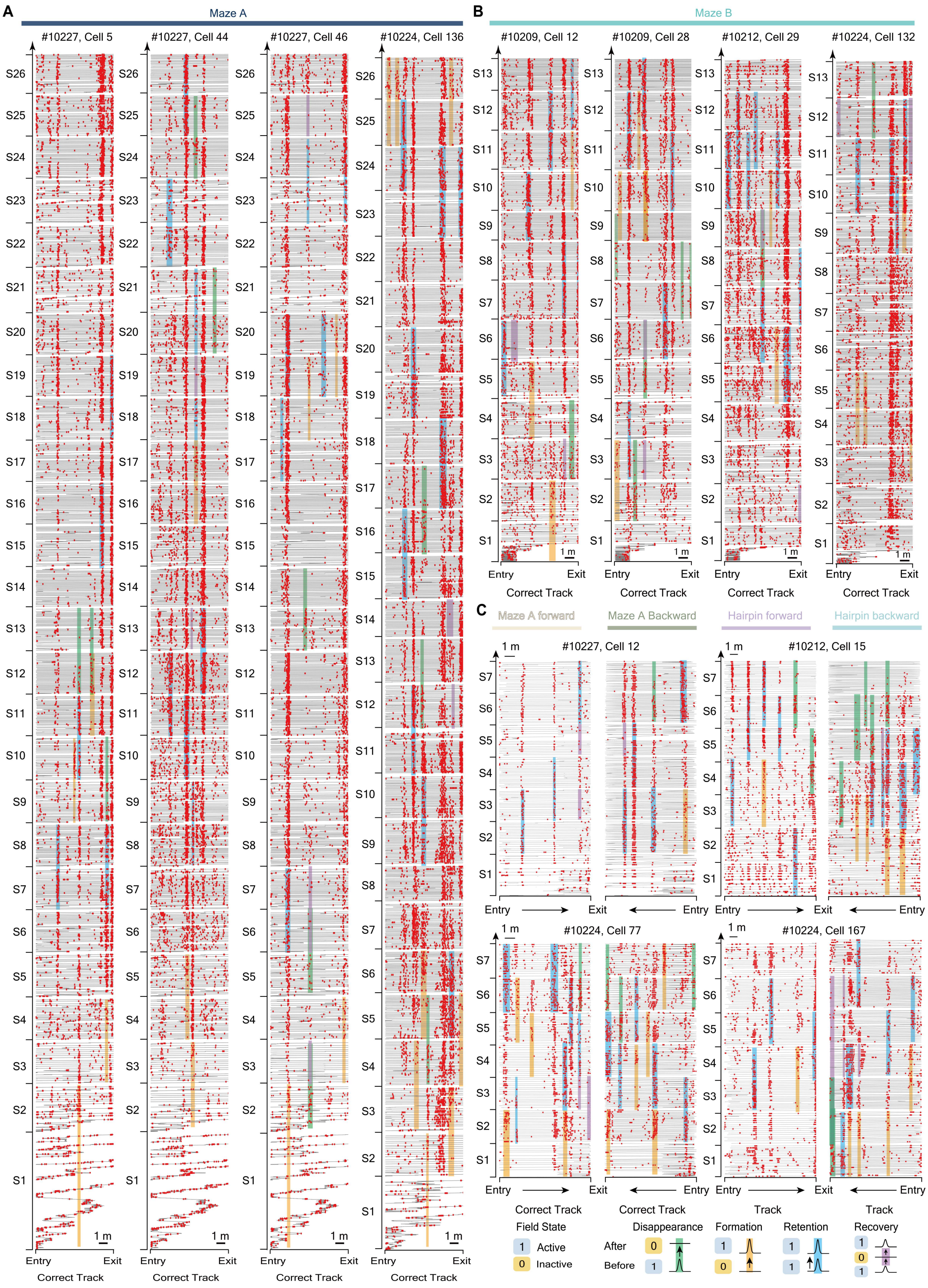

A
Maze A
#10227, Cell 5
#10227, Cell 44
#10227, Cell 46
#10224, Cell 136
Entry
Exit
Correct Track
1 m
B
Maze B
#10209, Cell 12
#10209, Cell 28
#10212, Cell 29
#10224, Cell 132
C
Maze A forward
Maze A Backward
Hairpin forward
Hairpin backward
#10227, Cell 12
#10212, Cell 15
#10224, Cell 77
#10224, Cell 167
Track
Field State
1 Active
0 Inactive
Disappearance
Formation
Retention
Recovery
After
Before

**Figure S10. Examples place cells demonstrating diverse evolution events in the long-term dynamics of hippocampal multi-field spatial maps**

(**A to C**) 12 exemplary place cells demonstrate a spectrum of intricate evolution of place fields in their spatial maps, which were widely recorded across distinct paradigms and spatial maps. These changes were classified into three fundamental categories of evolution events: disappearance, formation, and retention of place fields, marked with green, yellow, and blue shadows, respectively. Beyond these basic categories, we documented the recovery of previously weakened place fields, highlighted by purple shadows to indicate their periods of dormancy. Red bars indicate individual calcium events, and the background gray lines depict the mice's trajectories on the correct track of Maze A throughout the maze navigation paradigm training. Sessions are labeled as S1 to S26 (corresponding to Stage 2 S13).

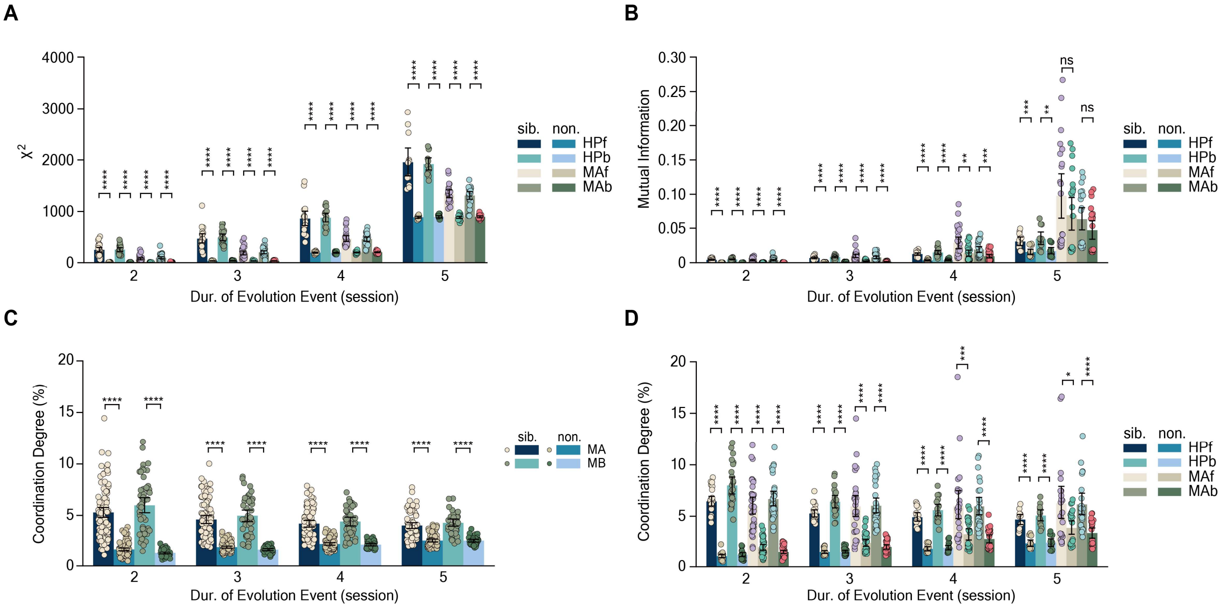


**Figure S11. Coordination in the evolution of sibling place fields in reversed and hairpin maze paradigms.**

(**A** and **B**) Analysis of $\chi^2$ (**A**), mutual information (MI) (**B**), and degree of coordination (CD)

(**C** and **D**) metrics reveals the presence of some extent of coordination in the evolution of sibling place fields during forward (HPf and MAf) and backward (HPb and MAb) movements in the hairpin maze (HP) and Maze A (MA), respectively. Statistical comparisons using Welch's t-test show significant differences in $\chi^2$, MI, and CD values between sibling (sib.) and non-sibling (non.) groups for all durations (dur.) of evolution events (all $\chi^2$ comparisons, $P < 0.001$; all MI comparisons, $P < 0.02$; all CD comparisons, $P < 0.02$). The degree of coordination, albeit present, is relatively subtle (for duration = 2: MA, 5.3 ± 2.5%; MB, 6.0 ± 2.7%; MAf, 6.4 ± 1.2%; MAb, 8.0 ± 2.0%; HPf, 6.0 ± 2.4%; HPb, 6.7 ± 2.0%. For duration = 3: MA, 4.6 ± 1.9%; MB, 5.0 ± 1.9%; MAf, 5.3 ± 0.8%; MAb, 6.4 ± 1.3%; HPf, 6.0 ± 2.9%; HPb, 6.0 ± 2.0%. For duration = 4: MA, 4.2 ± 1.5%; MB, 4.4 ± 1.4%; MAf, 4.9 ± 0.8%; MAb, 5.6 ± 1.1%; HPf, 6.0 ± 3.5%; HPb, 6.0 ± 2.2%. For duration = 5: MA, 4.0 ± 1.3%; MB, 4.3 ± 1.1%; MAf, 4.7 ± 0.9%; MAb, 5.1 ± 1.0%; HPf, 6.2 ± 3.9%; HPb, 6.2 ± 2.5%). Significance levels: ns, $P \geq 0.05$; *, $P < 0.05$; **, $P < 0.01$; ***, $P < 0.001$; ****, $P < 0.0001$.

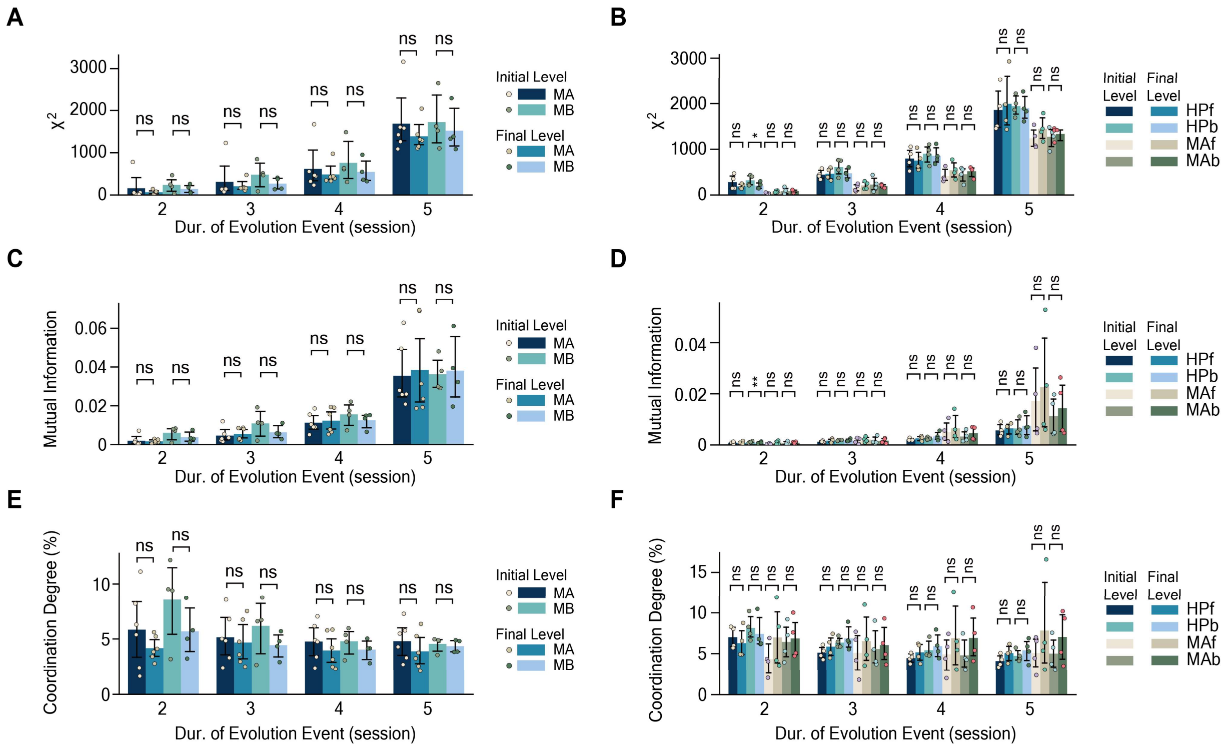


**Figure S12. The degree of coordination preserves throughout familiarization.**

Comparisons of $\chi^2$ (**A** and **B**), mutual information (MI) (**C** and **D**), and the degree of coordination (CD) (**E** and **F**) confirmed that the extent of coordination among sibling place fields preserves throughout familiarization, observed across multiple behavioral paradigms, irrespective of the duration selected to define evolutionary events (Paired t-test for the comparison between the initial level and final level of $\chi^2$, MI, and CD for any given duration of evolution events). Significance levels: ns, $P \geq 0.05$; *, $P < 0.05$; **, $P < 0.01$; ***, $P < 0.001$; ****, $P < 0.0001$.

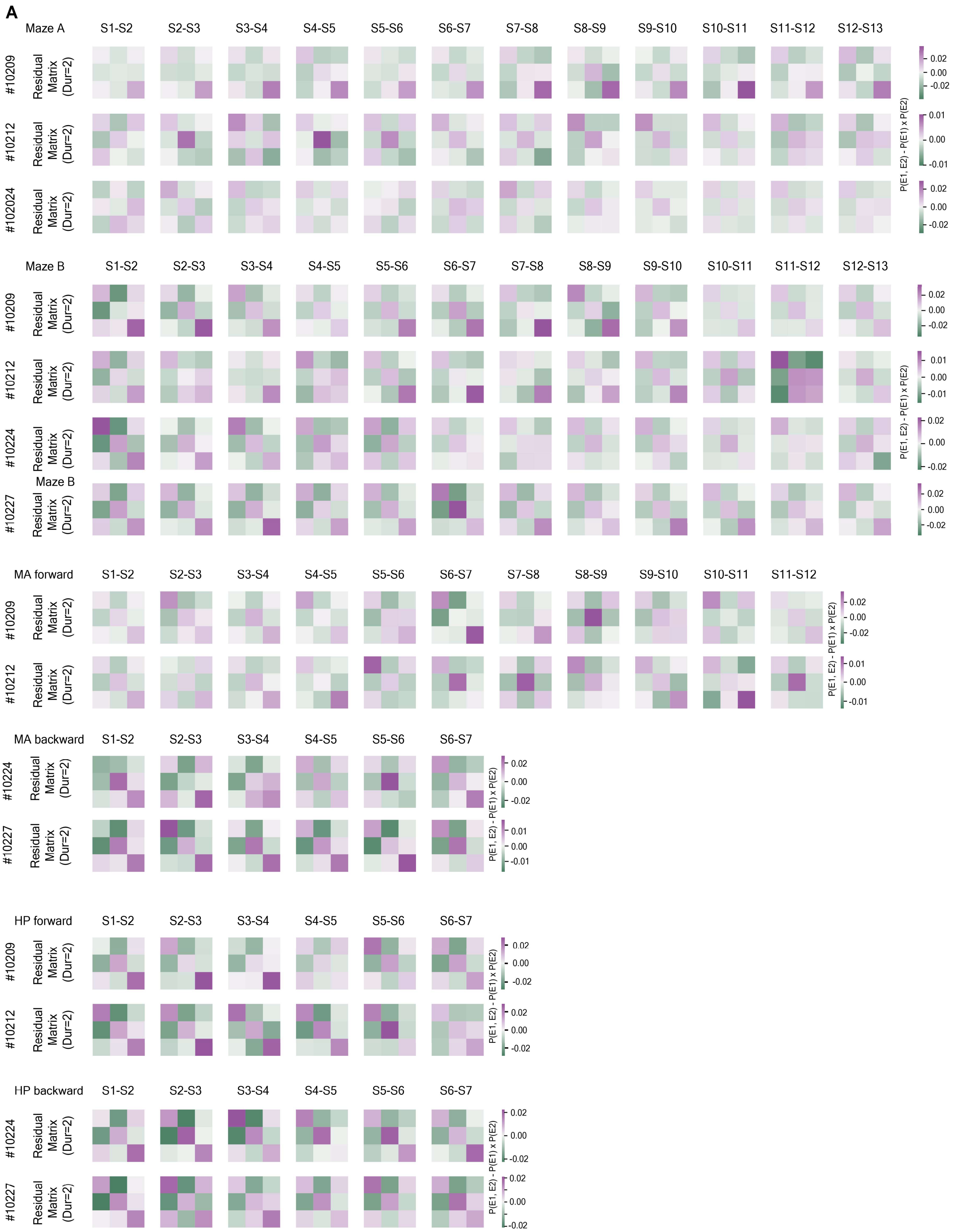
A
Maze A
S1-S2
S2-S3
S3-S4
S4-S5
S5-S6
S6-S7
S7-S8
S8-S9
S9-S10
S10-S11
S11-S12
S12-S13
#10209
#10212
#102024
Residual Matrix (Dur=2)
Maze B
#10224
#10227
MA forward
MA backward
HP forward
HP backward
P(E1, E2) - P(E1) x P(E2)
0.02
0.00
-0.02
0.01
-0.01

**Figure S13. Analysis of residual matrices suggest an endured preference for synchronized evolution of sibling place fields.**

The residual matrices for evolution events with a duration of two are 3 × 3 square matrices. Each entry at row i and column j in the residual matrix represents the extent to which the simultaneous occurrence of evolution events i and j for two sibling place fields deviates from what would be expected if the evolution of two sibling fields is independent. Darker colors indicate greater deviations from expectation. This panel visually presents some residual matrices from four mice across different behavioral paradigms in temporal order. Entries along the principal diagonal of these matrices tend to be positive (purple-colored), implying a preference for synchronized evolution of sibling place fields. Sessions S1 to S26 correspond to Session 1 to Session 26 (i.e., Stage 2 S13).

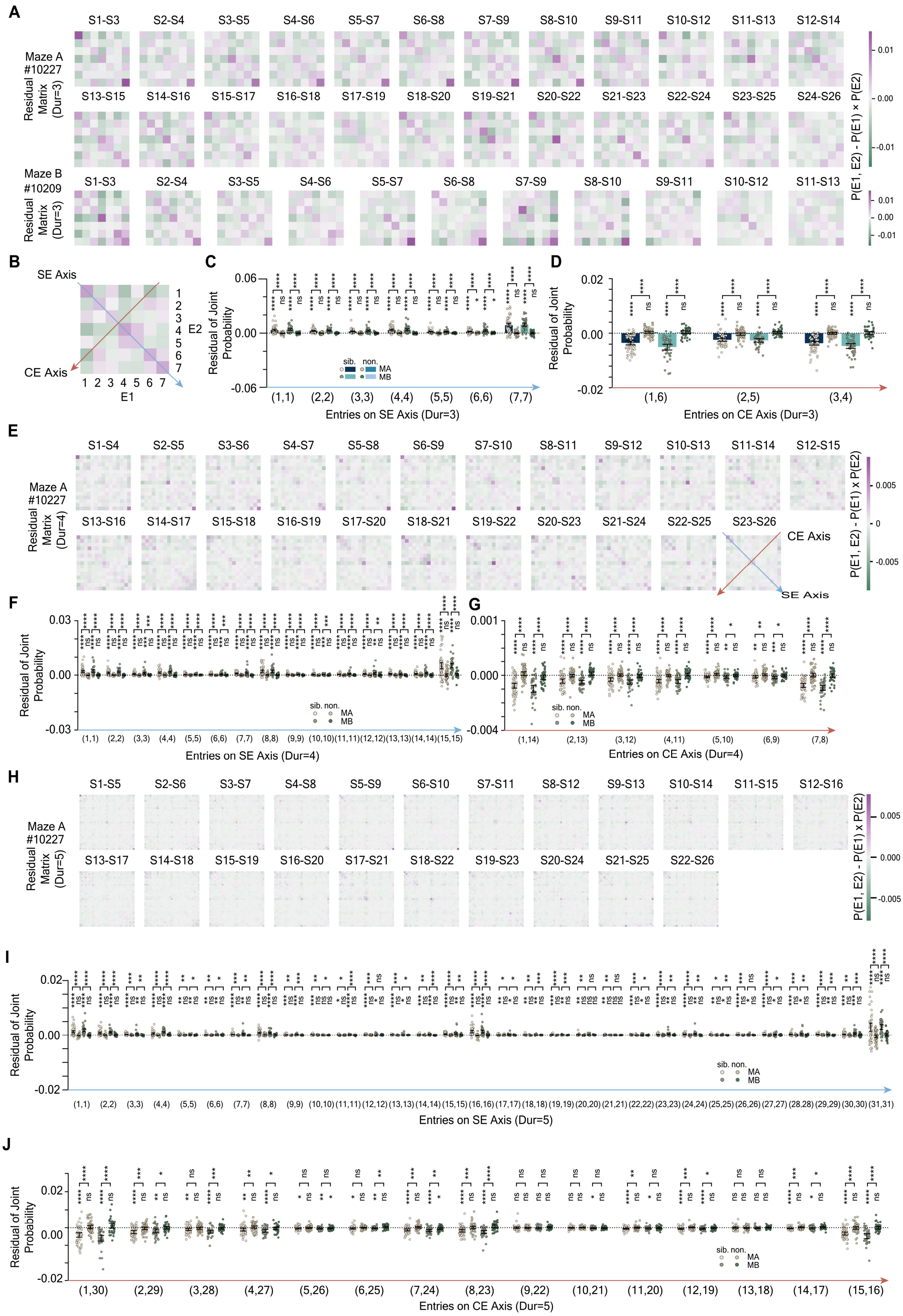
A
Maze A
#10227
Residual Matrix (Dur=3)
S1-S3
S2-S4
S3-S5
S4-S6
S5-S7
S6-S8
S7-S9
S8-S10
S9-S11
S10-S12
S11-S13
S12-S14
S13-S15
S14-S16
S15-S17
S16-S18
S17-S19
S18-S20
S19-S21
S20-S22
S21-S23
S22-S24
S23-S25
S24-S26
Maze B
#10209
Residual Matrix (Dur=3)
P(E1, E2) - P(E1) × P(E2)
0.01
0.00
-0.01
B
SE Axis
CE Axis
E1
E2
C
Residual of Joint Probability
Entries on SE Axis (Dur=3)
(1,1) (2,2) (3,3) (4,4) (5,5) (6,6) (7,7)
sib. non.
MA
MB
D
Entries on CE Axis (Dur=3)
(1,6) (2,5) (3,4)
E
Maze A
#10227
Residual Matrix (Dur=4)
S1-S4
S2-S5
S3-S6
S4-S7
S5-S8
S6-S9
S7-S10
S8-S11
S9-S12
S10-S13
S11-S14
S12-S15
S13-S16
S14-S17
S15-S18
S16-S19
S17-S20
S18-S21
S19-S22
S20-S23
S21-S24
S22-S25
S23-S26
CE Axis
SE Axis
0.005
0
-0.005
F
Entries on SE Axis (Dur=4)
G
Entries on CE Axis (Dur=4)
(1,14) (2,13) (3,12) (4,11) (5,10) (6,9) (7,8)
H
Maze A
#10227
Residual Matrix (Dur=5)
S1-S5
S2-S6
S3-S7
S4-S8
S5-S9
S6-S10
S7-S11
S8-S12
S9-S13
S10-S14
S11-S15
S12-S16
S13-S17
S14-S18
S15-S19
S16-S20
S17-S21
S18-S22
S19-S23
S20-S24
S21-S25
S22-S26
I
Entries on SE Axis (Dur=5)
J
Entries on CE Axis (Dur=5)
(1,30) (2,29) (3,28) (4,27) (5,26) (6,25) (7,24) (8,23) (9,22) (10,21) (11,20) (12,19) (13,18) (14,17) (15,16)

**Figure S14. Sibling place fields exhibit a preference for synchronized evolution, contributing to limited coordination in MNP.**

(**A**) Illustrated here are two sets of example residual matrices with durations of three (7 × 7 matrices) from Maze A (first two rows) and Maze B (last row). Each entry at row i and column j, entry (i, j), in the residual matrix represents the extent to which the simultaneous occurrence of evolution events i and j for two sibling place fields deviates from what would be expected by chance. Darker colors denote larger deviations from the expected values. Notably, entries along the principal diagonal tend to be positive (purple-colored), suggesting a tendency for synchronized evolution among sibling place fields. Session labels S1 to S26 correspond to Sessions 1 to 26 (Stage 2 S13).

(**B**) For further analysis, entries along the axes of synchronized events (SE, blue arrow) and complementary events (CE, dark red arrow) were extracted.

(**C** and **D**) Residual values along the SE axis (C) and CE axis (D) are plotted in the order indicated by their respective arrows in (B). We performed statistical comparisons between sibling and non-sibling field groups (two-sample two-sided t-test, significance indicated by stars above brackets) and tested each group's residuals against zero (one-sample two-sided t-test, significance indicated by stars below brackets). The tuple (i, j) denotes entries located at row i and column j of the residual matrices.

(**E** to **J**) This analysis was extended to evolution events with durations of four (**E** to **G**) and five (**H** to **J**). Most entries along the SE axis remain significantly positive across various durations of evolution events (Fig. 4I; fig S14C, F, and I), indicating a mild but consistent preference for synchronized evolution among sibling place fields. In contrast, most entries along the CE axis exhibit significantly lower levels than expected for durations of two (Fig. 4J) to five (fig S14D, G and J) sessions. Significance levels: ns, $P \geq 0.05$; *, $P < 0.05$; **, $P < 0.01$; ***, $P < 0.001$; ****, $P < 0.0001$.

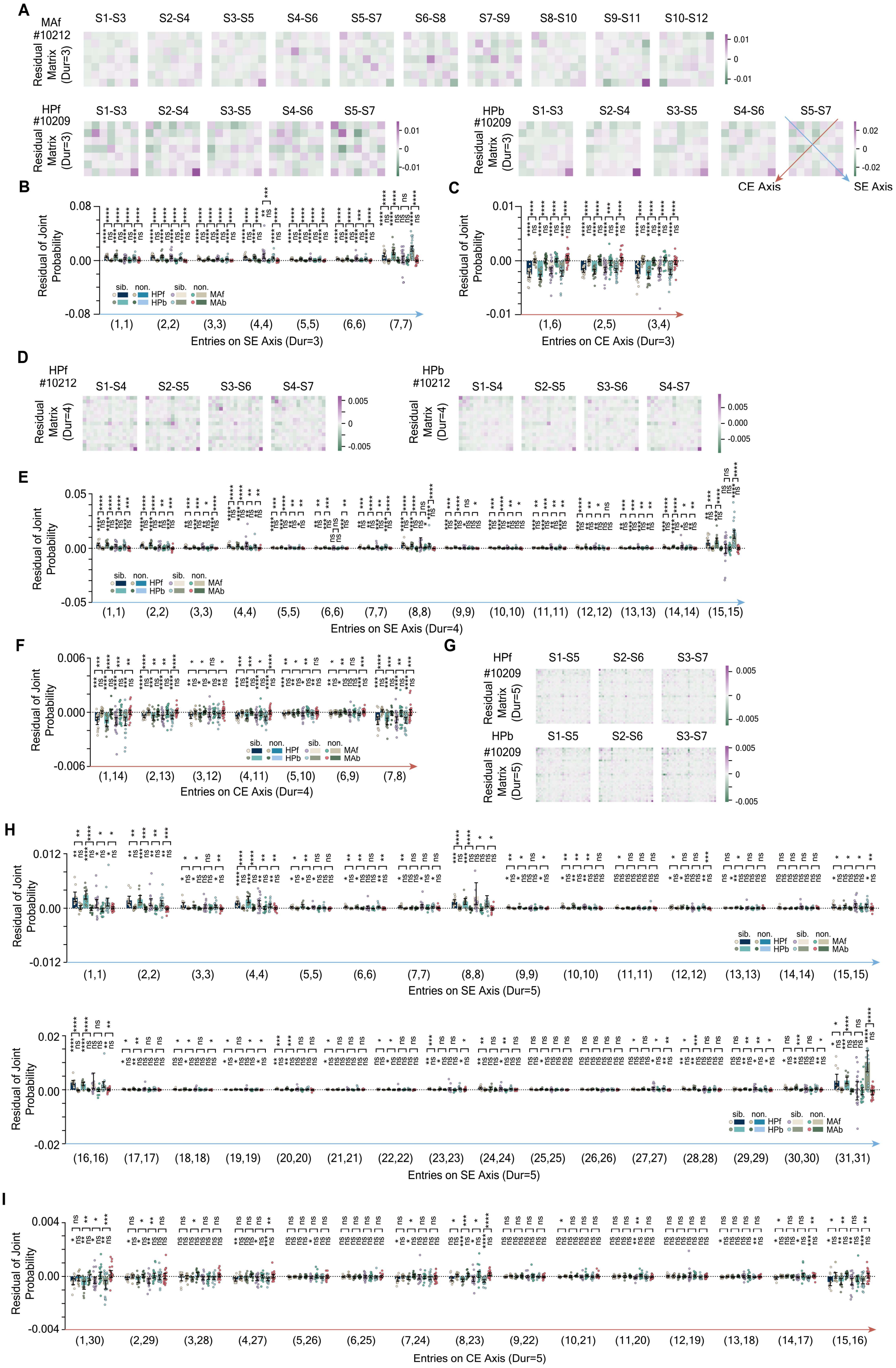
A
MAf #10212
Residual Matrix (Dur=3)
S1-S3 S2-S4 S3-S5 S4-S6 S5-S7 S6-S8 S7-S9 S8-S10 S9-S11 S10-S12
0.01 0 -0.01
HPf #10209
S1-S3 S2-S4 S3-S5 S4-S6 S5-S7
HPb #10209
CE Axis
SE Axis
0.02 0 -0.02
B
Residual of Joint Probability
Entries on SE Axis (Dur=3)
sib. non. HPf HPb MAf MAb
C
Entries on CE Axis (Dur=3)
D
HPf #10212
HPb #10212
Residual Matrix (Dur=4)
S1-S4 S2-S5 S3-S6 S4-S7
E
Entries on SE Axis (Dur=4)
F
Entries on CE Axis (Dur=4)
G
HPf #10209
HPb #10209
Residual Matrix (Dur=5)
S1-S5 S2-S6 S3-S7
H
Entries on SE Axis (Dur=5)
I
Entries on CE Axis (Dur=5)

**Figure S15. Sibling place fields exhibit a preference for synchronized evolution, contributing to limited coordination in RMP and HMP.**

(**A**) Illustrated here are three sets of example residual matrices with durations of three (7 × 7 matrices) from forward movement in Maze A (first row), forward (bottom left) and backward movement (bottom right) in the hairpin maze. Each entry at row i and column j in the residual matrix represents the extent to which the simultaneous occurrence of evolution events i and j for two sibling place fields deviates from what would be expected by chance. Darker colors denote larger deviations from the expected values. Notably, entries along the principal diagonal tend to be positive (purple-colored), suggesting a tendency for synchronized evolution among sibling place fields. Session labels S1 to S12 correspond to Sessions 1 to 12. (**B** to **C**) Residual values along the SE axis (B) and CE axis (C) are plotted in the order indicated by their respective arrows. We performed statistical comparisons between sibling and non-sibling field groups (two-sample two-sided t-test, significance indicated by stars above brackets) and tested each group's residuals against zero (one-sample two-sided t-test, significance indicated by stars below brackets). The tuple (i, j) denotes entries located at row i and column j of the residual matrices. This analysis was extended to evolution events with durations of four (**D** to **F**) and five (**G** to **I**). Most entries along the SE axis remain significantly positive across various durations of evolution events (Figures 4K, S15B, S15E and S15H), indicating a mild but consistent preference for synchronized evolution among sibling place fields. In contrast, most entries along the CE axis exhibit significantly lower levels than expected for durations of two (Figure 4L) to five (Figures S15C, S15F and S15I). Consistent with observations in Maze A and Maze B, this suggests that the observed coordination is featured by a subtle but enduring preference for synchronized evolution and a disfavor for alternate evolution among sibling place fields. Significance levels: ns, $P \geq 0.05$; *, $P < 0.05$; **, $P < 0.01$; ***, $P < 0.001$; ****, $P < 0.0001$.

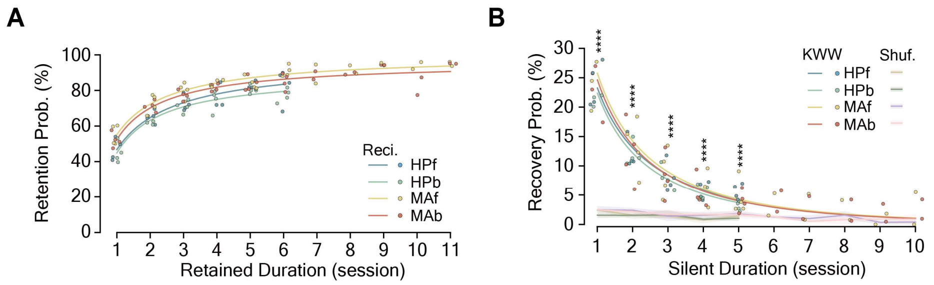


**Figure S16. Single field evolution rule is consistent across distinct environments and goal-directed navigation paradigms.**

(**A**) Field retention probability measured in both movement directions during reversed and hairpin maze paradigms, fitted with the 'reci.' function, showing a similar trend to Maze A and B during maze navigation paradigm (Figure 5D).

(**B**) Field recovery probability fitted with the Kohlrausch-Williams-Watts (KWW) function. Chance levels computed by re-locate shuffle test (see Methods) are indicated by lines with shadows, and significance levels are denoted by colored stars. Significance levels: ***, P < 0.001; ****, P < 0.0001; No stars indicate no significant differences between real data and shuffled results. Abbreviations: MA, Maze A; MB, Maze B; HP, hairpin maze; f, forward; b, backward.

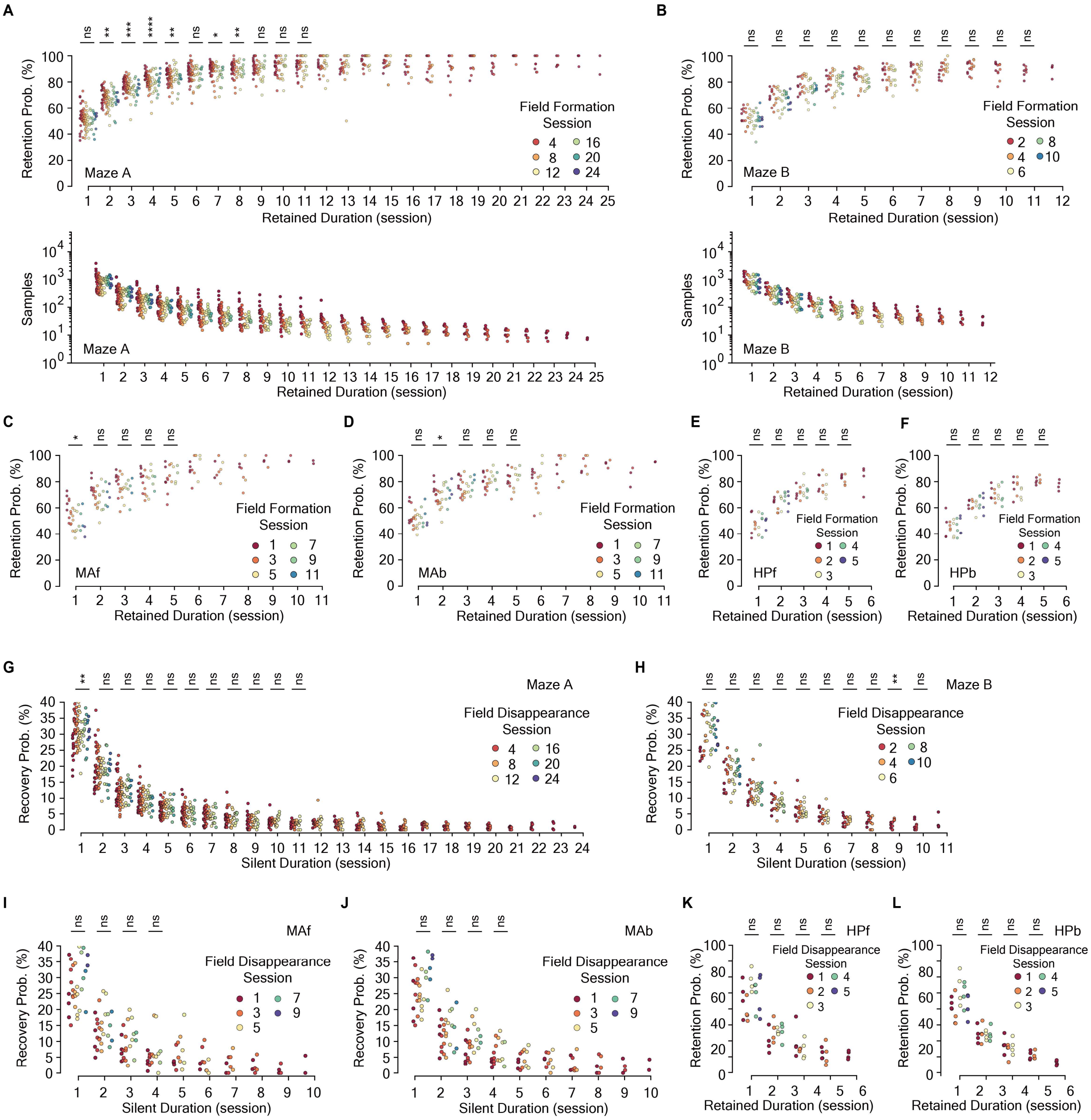


**Figure S17. Single field evolution rule is learning-independent.**

(**A** to **F**) Place fields were grouped based on the sessions during which they formed (field formation session), and the retention probability was calculated for (A), MB (B), MAf (C), MAb (D), HPf (E), and HPb (F). The results are presented as circles with different colors corresponding to the field formation sessions. The bottom figures in (A) and (B) displayed the number of samples used for probability calculations. We compared the related probability values across different groups of place fields using one-way ANOVA to determine if the field formation session significantly impacts the retention probability. Only fields formed during the first 11 sessions in MA and MB, and during the first 5 sessions in MAf, MAb, HPf, and HPb, were eligible for comparison to maintain a consistent number of data points (n = 6 in MA and n = 4 in the rest).

(**G** to **L**) Similarly, place fields were grouped based on the sessions during which they disappeared, and the recovery probability was calculated for MA (G), MB (H), MAf (I), MAb (J), HPf (K), and HPb (L). The results were presented in circles with different colors corresponding to the field disappearance session. Similarly, only fields that disappeared during the first 10 sessions in MA and MB, and during the first 4 sessions in MAf, MAb, HPf, and HPb, were eligible for comparison. The data show no significant differences across place fields of distinct groups, suggesting that SFER operates independently of the spatial learning process. Significance levels: ns, P ⩾ 0.05; *, P < 0.05; **, P < 0.01; ***, P < 0.001; ****, P < 0.0001.

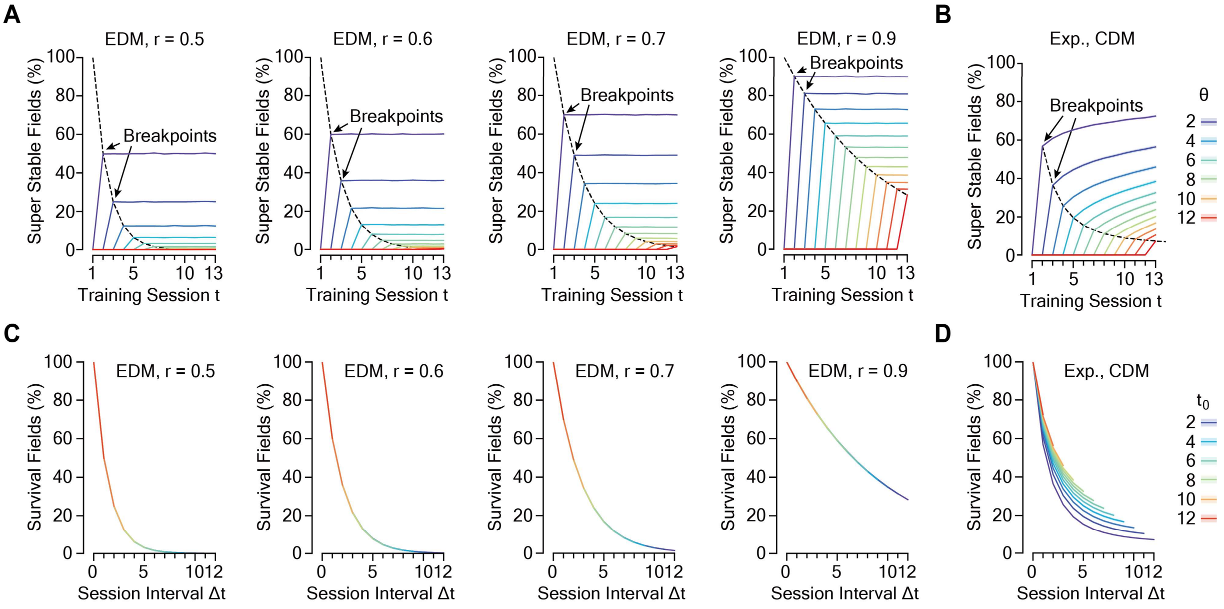


**Figure S18. Single field evolution rule is sufficient to account for the dynamics of hippocampal spatial representation at the ensemble level.**

(**A** to **D**) Dynamics of the proportion of super stable place fields (SupS) for equal-rate drift models (EDMs) with distinct parameter r (A), and convergent drift models (CDMs) with 'exp.' function (B). In EDMs (A), the proportion of SupS, defined by a threshold θ, is a function of training session t and drift rate parameter r. The black dotted line represents the distribution of breakpoints as a function of θ. Colored lines show the dynamics of SupS for different θ values, with redder lines indicating higher θ. In CDMs (B), SupS is influenced by session t and the parameters of the convergent function, with breakpoints converging to a non-negative value. (**C** and **D**) Dynamics of the proportion of survival place fields (Surv) over daily drift for EDMs with distinct parameter r (C) and CDMs with 'exp.' function (D). In EDMs (C), Surv is an exponential function of session intervals Δt, regardless of the start session $t_0$, with different start sessions visualized in distinct colors. The decay rate is controlled by parameter r. In CDMs (D), Surv is a function of session intervals Δt and the parameters of the convergent function, with a black arrow indicating the cumulative effect.
